# A zebrafish leprosy model identifies interferon and plasminogen signaling as survival determinants in mycobacterial infection

**DOI:** 10.64898/2026.07.31.739979

**Authors:** Beatrice Junqueira de Souza, Robin Rio, Giovanni Galletti, William Morrill, Cressida A. Madigan

## Abstract

Although many genes are associated with leprosy, a skin and nerve infection by *Mycobacterium leprae*, the function of most of these genes in infection remains unknown. This is partly due to a paucity of animal models that are genetically malleable and recapitulate features of the human disease. Zebrafish, a recent leprosy model, have human-like responses to *M. leprae*, including macrophage-mediated inflammation and neurodegeneration. We confirm this at the transcriptional level, using RNA sequencing (RNAseq) of chronic *M. leprae* infection of adult zebrafish. This identified regulated zebrafish orthologs of human leprosy-associated genes, including β- and γ-interferon, IL-6, IL-10, IL-4, and TNF superfamily members. Other regulated genes have not been previously associated with leprosy. Genes associated with tuberculoid leprosy (T-lep) were largely downregulated, while lepromatous leprosy (L-lep) genes were upregulated. Pathways relevant to leprosy, such as phagocytosis and antiviral responses, differed between early and late stages of infection. In infected rag1 mutant zebrafish, which lack T and B cells, T-lep genes were downregulated, suggesting that adaptive immunity is required for their expression. Key pathways and genes were validated by qPCR using zebrafish infection with *M. marinum:PGL-1,* a model pathogen that can express *M. leprae* genes. This system allowed for the identification of validated *M. leprae*-regulated genes that alter infection outcomes, by using multiplex CRISPR to simultaneously mutate many zebrafish genes. 17 genes were screened and 2 were identified that increased mortality infection: isg15 (interferon-stimulated gene 15) and serpine1 (plasminogen activator inhibitor-1). This work confirms that, in humans and zebrafish, type I interferon and plasminogen activation are required for survival of mycobacterial disease, and demonstrates that multiplex CRISPR can identify mediators of host defense in vivo.

## INTRODUCTION

Clinical data have identified many genes associated with leprosy in humans^1^. Some of these genes have been connected to the divergent immune responses to *M. leprae* that influence the presentation and symptoms of the disease. For example, mediators of Th1 responses, including IFNγ, TNF, IL-12 and IL-6, are associated with efficient *M. leprae* killing in tuberculoid leprosy (T-lep). In contrast, Th2 mediators such as IFNβ, IL-10, and IL-4 are associated with *M. leprae* proliferation in lepromatous leprosy (L-lep). Immunologically unstable patients can shift between L-lep and T-lep responses during type 1 (R1) or type 2 (R2) leprosy reactions, which are associated with a dramatic exacerbation of inflammation and nerve injury. These are associated with IFNγ and TNF in R1, and IL-1β, NLRP3, and IL-10 in R2^1^. It seems clear that disparate immune responses to *M. leprae* are critical mediators of the immunological and neurological outcomes of leprosy in humans.

Beyond these classic immune response genes, the function of most other leprosy-associated genes in infection is unknown. This is partly because *M. leprae* cannot replicate in cultured cells^2^, instead requiring growth in an animal that is maintained at ∼33°C. For the study of *M. leprae* beyond a few days, a “cold-blooded” animal model is needed that can be maintained at ∼33°C. While it lacks most molecular and genetic tools, the armadillo has been used for this purpose^3, 4^. However, a recent model host for *M. leprae* infection, the zebrafish, is maintained at cooler temperatures (∼28°C), can be genetically modified, and allows for screening and imaging approaches that are not feasible in armadillos, mice, and cell culture^5, 6^.

A genetically malleable animal model that recapitulates aspects of human leprosy would contribute to identifying disease-causing pathways by infection of a genetically modified host. To this end, we have employed an unusual model of *M. leprae* infection^5, 6^: zebrafish. The similarities between zebrafish and humans outweigh their differences: over 70% of human genes have a known zebrafish ortholog^7^. Their innate responses to infection are similar, too: polarized macrophages phagocytose bacteria^8^, neutrophils deploy NETs^9^, and dendritic cells present antigen^10^. Zebrafish form inflammatory granulomas (macrophage aggregates) in response to *M. leprae* and *M. marinum,* a model pathogen of fish. Granulomatous inflammation predicts infection severity in primates and zebrafish^11^, and *M. leprae* granulomas in zebrafish bear a striking histological similarity to those of human leprosy. In adult zebrafish, granulomas containing *M. leprae* bacilli persist through at least 5 months post-infection. Therefore, zebrafish models of infection by *M. leprae*, or by *M. marinum* expressing *M. leprae* genes, recapitulate aspects of human leprosy, and can contribute to elucidating previously-unknown mechanisms that drive leprosy pathogenesis.

Here, we use transcriptomics to follow the course of a chronic *M. leprae* infection over 8 months. We identify infection-induced regulation of 347 zebrafish orthologs of human genes associated with leprosy. These findings are confirmed using the larval zebrafish model infected with recombinant *M. marinum* expressing an *M. leprae* glycolipid, *M. marinum:PGL1^5^*. Then, multiplex CRISPR gene editing is used to mutate pools of genes in zebrafish and assess their contribution to infection. Two genes were identified in this way, isg15 and serpine1, which worsened survival in infection. These findings demonstrate the utility of this method to identify genes required for the long-term immune response to *M. leprae*.

### Zebrafish kill *M. leprae* within weeks, regardless of adaptive immunity

To identify differentially regulated genes (DEGs) in *M. leprae*-infected zebrafish, 4-month-old adult zebrafish were infected with 10^7^ colony forming units (CFU) of *M. leprae* Thai-53, as described^6^. Sibling rag1(+/-) heterozygotes, which have intact immunity, and rag1(-/-) mutants, which lack T and B cells, were infected and maintained for 240 days (Supplemental Table 1). Infected and uninfected zebrafish were sacrificed at 0, 7, 30, 120, 180, and 240 days post infection (dpi) for RNAseq.

In human leprosy, efficient antimicrobial responses of macrophages kill *M. leprae,* which primarily reside in the macrophages of inflammatory granulomas. To detect quantify viable *M. lepra*e over time in adult zebrafish, qPCR was used to detect 16S rRNA and RLEP, which are indicators of transcriptional activity^12^ (Figure 1a,b). The ratio of RLEP to 16S indicated *M. leprae* viability through the first few weeks of infection, with 63% of the initial inoculum remaining at 7 dpi, and nearly 0% at 180 dpi (Figure 1a). Therefore, early responses to infection (0-30 dpi) are associated with viable *M. leprae,* while late responses (120-240 dpi) are associated with killed *M. leprae* (Figure 1a). The kinetics of *M. leprae* killing were similar regardless of the zebrafish rag1 genotype (Figure 1b), suggesting that zebrafish killing of *M. leprae* early in infection does not require adaptive immunity.

**Figure 1.**
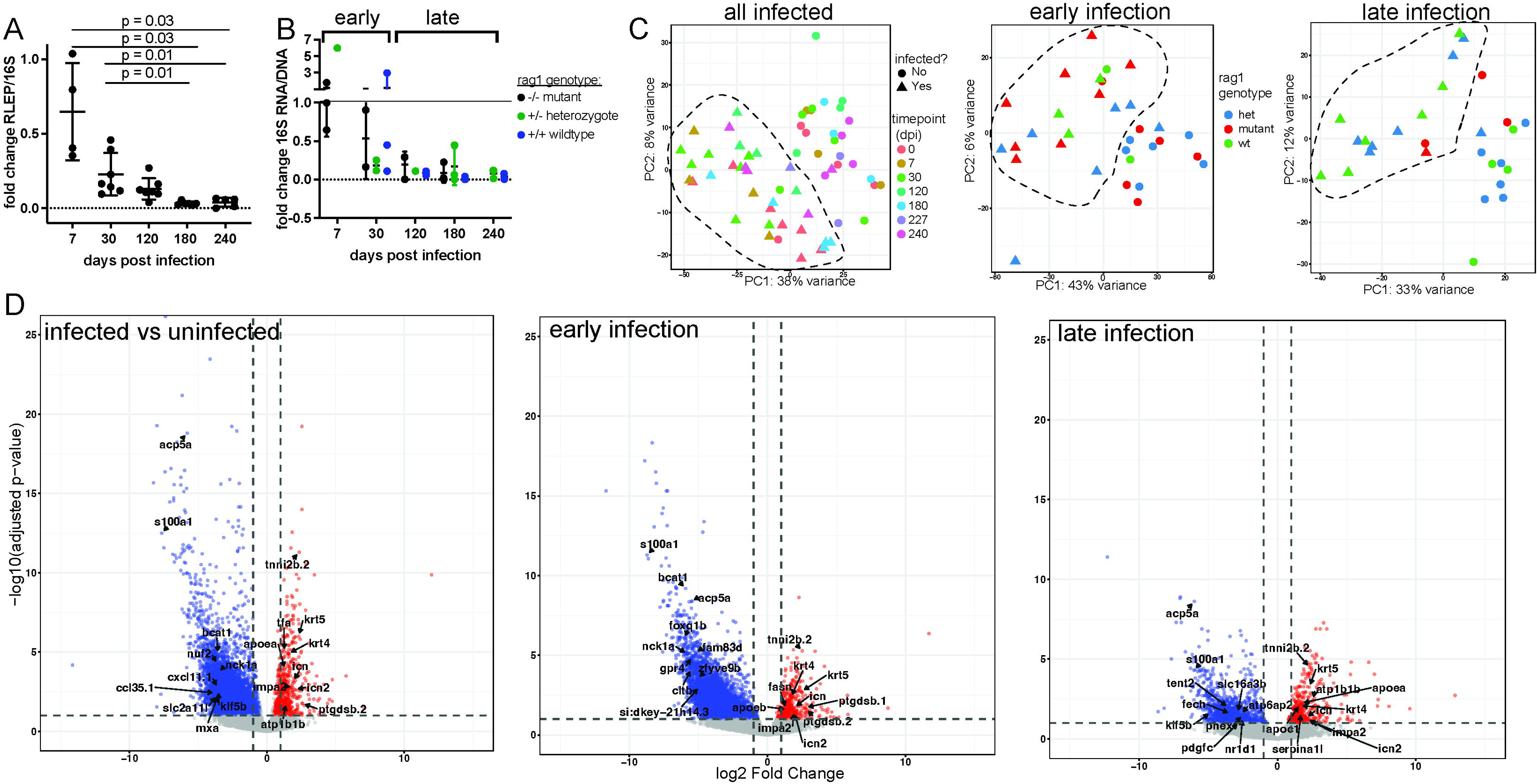
*M. leprae* viability is associated with zebrafish gene expression. (A, B) *M. leprae* viability in infected adult zebrafish was quantified by measuring the ratio of 16S rRNA to 16S or RLEP DNA. Viability decreases from 7 dpi (days post infection) to 240 dpi, when nearly all bacteria are non-viable. The fold change over time in 16S rRNA:DNA was unaffected by the zebrafish rag1 genotype. Students’s t test. (C) PCA plots show a loose association between infected samples at all timepoints; this was independent of rag1 genotype (Supplemental Figure 1). (D) Volcano plots labeling the top 10 down- and upregulated zebrafish orthologs for human genes associated with leprosy (Supplemental Table 2). Significant DEGs have a fold change > +/-2, and adjusted p-value of <0.05.

### *M. leprae* infection modifies zebrafish gene expression over months

RNA extracted from the zebrafish was sequenced to a depth of ∼60 million reads, and principle component analysis (PCA) was used to detect the sources of variance in the samples. Samples loosely clustered by infection status, but not by rag1 genotype or timepoint (Figure 1C). Further, sex was associated with 19% variance comparing infected and uninfected zebrafish (Supplemental Figure 1), likely due to unequal sex ratios between the groups (Supplemental Table 1). To accommodate this, only zebrafish of the same sex were compared, or sex-regulated genes were identified and removed before analysis (see Methods).

Genes that were differentially expressed (DEGs) during infection were identified (adjusted p-value <0.05 and fold change > +2 or -2) (Figure 1d and Supplemental Table 2), with most downregulated early in infection. To demonstrate that this downregulation was genuine, the expression of 4 zebrafish housekeeping genes was assessed and found to be unchanged by infection status, timepoint, or genotype (Supplemental Figure 2). Volcano plots confirm that many genes are regulated at early timepoints, and fewer at later timepoints, and that more genes are downregulated than upregulated (Figure 1d).

### Chronic *M. leprae* infection regulates genes associated with human leprosy

To identify similarities between human and zebrafish responses to *M. leprae*, zebrafish orthologs for 347 human genes associated with leprosy were identified in the zebrafish RNAseq dataset (Supplemental Table 3). The top regulated leprosy DEGs include s100a1 (human S100A1), ccl35.1 (CCL4l1), mxa (MX2), apoea (APOE), and ptgdsb.2 (PTGDS) (Supplemental Tables 2 and 3). Most of these were expressed in early infection (Figure 2a), and overall, about 25% were associated with L-lep, R1 or R2 (Figure 2a). Further, 50% of the top 10 up-regulated leprosy DEGs were associated with L-lep, including ccl38a.4, ptgdsb.2, snrpb, and wasb, while 12-20% were associated with other forms (Figure 2c and Supplemental Table 2). This distribution changed by late timepoints, when most leprosy DEGs (33%) were associated with L-lep or R1 (Figure 2d).

**Figure 2.**
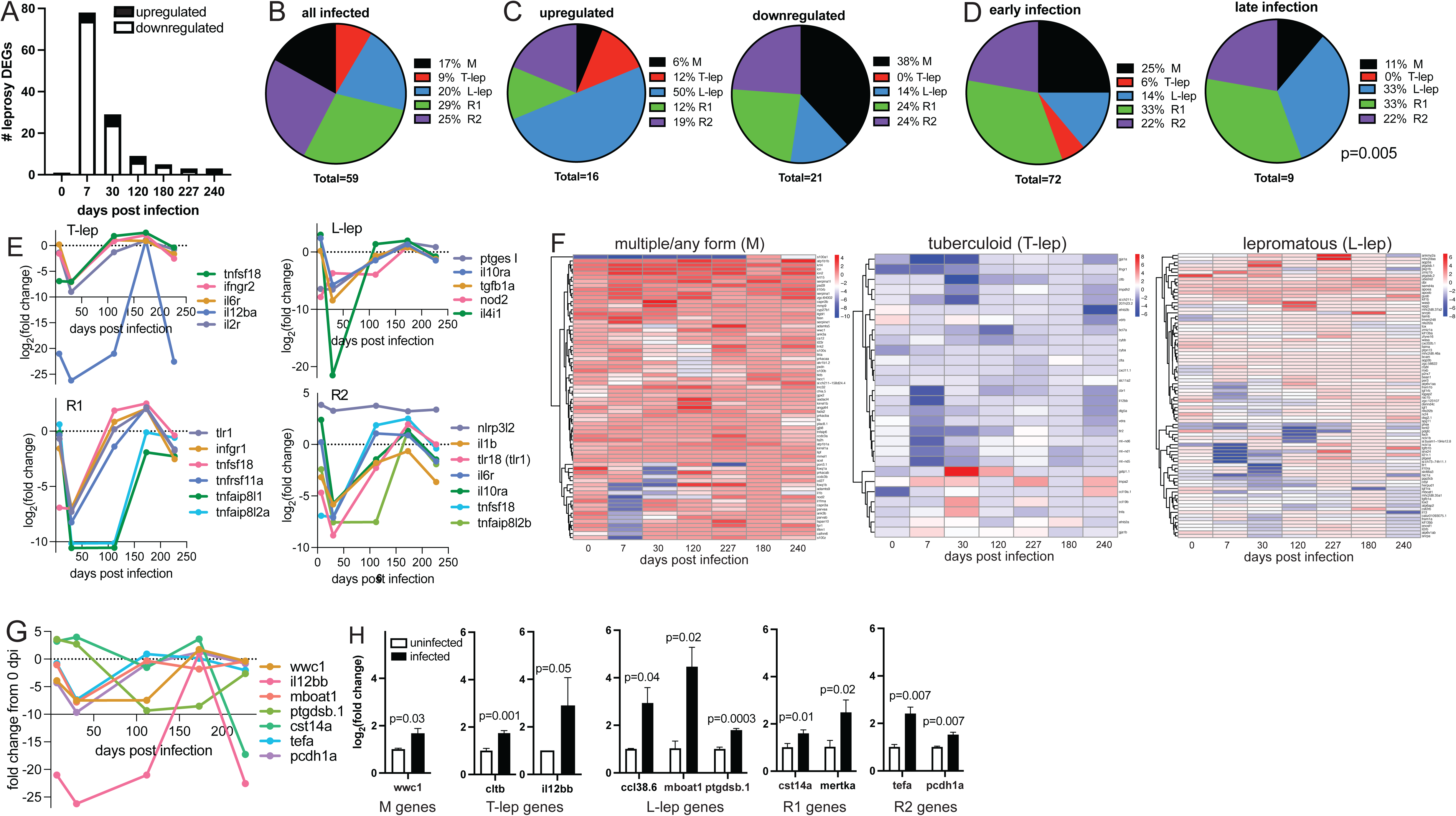
Orthologs of human leprosy-associated genes are regulated in *M. leprae*-infected zebrafish. (A) Zebrafish orthologs of human genes associated with leprosy were differentially expressed in infected zebrafish. (B) Comparing all infected to all uninfected zebrafish identified leprosy DEGs associated with multiple forms of leprosy (M), tuberculoid leprosy (T-lep), lepromatous leprosy (L-lep), type 1 reactions (R1), or type 2 reactions (R2). (C) The distribution of leprosy forms in the top 10-20 upregulated and downregulated leprosy DEGs. (D) The distribution of leprosy DEGs as in C, comparing early infection (7 dpi) to late infection (120 dpi) (p=0.005, Chi squared). (E) Gene trajectories showing detection of DEGs associated with each form of leprosy in humans. (F) Heatmaps of DEGs associated with M, T-lep, or L-lep, showing their regulation over time (0-240 dpi). (G, H) Expression over time of specific leprosy DEGs in RNAseq data, and (H) their expression in 4-5 dpi zebrafish larvae infected with ∼100 colony forming units *M. marinum:PGL-*1 (black), compared to uninfected (white). N=11-30 larvae per group. Student’s t-test.

Many leprosy-associated DEGs were downregulated at 7 dpi, which decreased through 240 dpi (Figure 2a). Cumulatively, 17% of leprosy DEGs were associated with unspecified or multiple forms of leprosy (M), 29% with type 1 reactions (R1), 25% with type 2 reactions (R2), 20% with lepromatous leprosy (L-lep), and 9% with tuberculoid leprosy (T-lep) (Figure 2b and Supplemental Table 2). 50% of the top regulated leprosy genes were associated with L-lep, including ccl38a.4, ptgdsb.2, snrpb, and wasb (Figure 2c). The top downregulated genes (38%) were associated with M leprosy.

The number and distribution of leprosy-associated genes differed over time. In early infection (7 dpi), most genes (33%) were associated with R1, while late infection was dominated by 33% L-lep genes and 33% R1 genes (Figure 2d and Supplemental Table 3). Examining the gene trajectories of typical leprosy genes, T-lep genes were largely inhibited in early infection, and recovered expression ∼100 dpi, as occurred with ifngr2, il12ba, and il6 (Figure 2e). Many typical L-lep genes followed the same pattern, such as il10ra, tgfb1a, and il4il. Genes associated with R1 (tlr1, ifng1, tnsf18 and tnfaip8l1) and R2 (nlrp3, il1b, il6) had more sustained downregulation, lasting through ∼180 dpi. Heatmaps indicated that non-specific leprosy genes (M) are largely induced by infection, except for when a subset are downregulated at 0-30 dpi (Figure 2f). In contrast, T-lep genes are largely downregulated, except for a brief upregulation of a subset of genes at 30 dpi. L-lep genes were intermediate, with mixed up- and down-regulation.

To confirm the regulation of these genes using another approach, 7 genes were identified that were regulated by *M. leprae* infection (Figure 2g) and associated with different forms of the disease (Figure 2g). Expression of these genes was then assessed in zebrafish larvae infected with ∼100 CFU *M. marinum:*PGL-1. At 4-5 dpi, infected and uninfected larvae were sacrificed for RNA extraction and subsequent qPCR, which detected regulation of wwc1 (M), cltb and il12bb (T-lep), ptgsb.1, mboat1, and ptgdsb.1 (L-lep), cst14a and mertka (R1), and tefa and pcdh1a (R2) (Figure 2h).

### Zebrafish respond to *M. leprae* by regulating known mycobacterial defense pathways

To assess the signaling pathways that are altered over time in M. leprae infection, KEGG pathways analysis was performed, which identified 161 regulated pathways in *M. leprae*-infected zebrafish (Figure 3a). Many are infection and inflammation pathways that are known or suspected to contribute to leprosy pathogenesis, including phagocytosis, mitophagy, antiviral responses, and signaling through innate receptors such as NOD-like receptors and TLRs (Figure 3a-b). Most pathway regulation occurred in early infection, and was absent or substantially decreased in late infection (Figure 3a-b). Pathways regulated in late infection include “herpes virus infection”, “Salmonella infection,” and others associated with mycobacterial infection, such as autophagy, cytokine-cytokine receptor interactions, and arginine metabolism. Comparing the trajectories for pathways over time revealed that 5 pathways are upregulated in early infection (endocytosis, arachidonic acid metabolism, steroid biosynthesis, lysosome biogenesis, and glycerolipid metabolism), while 2 are downregulated (oxidative phosphorylation and phagocytosis) (Figure 3c). This suggests that *M. leprae* infection results in a sustained immune response in zebrafish that persists for 6 months or more.

**Figure 3.**
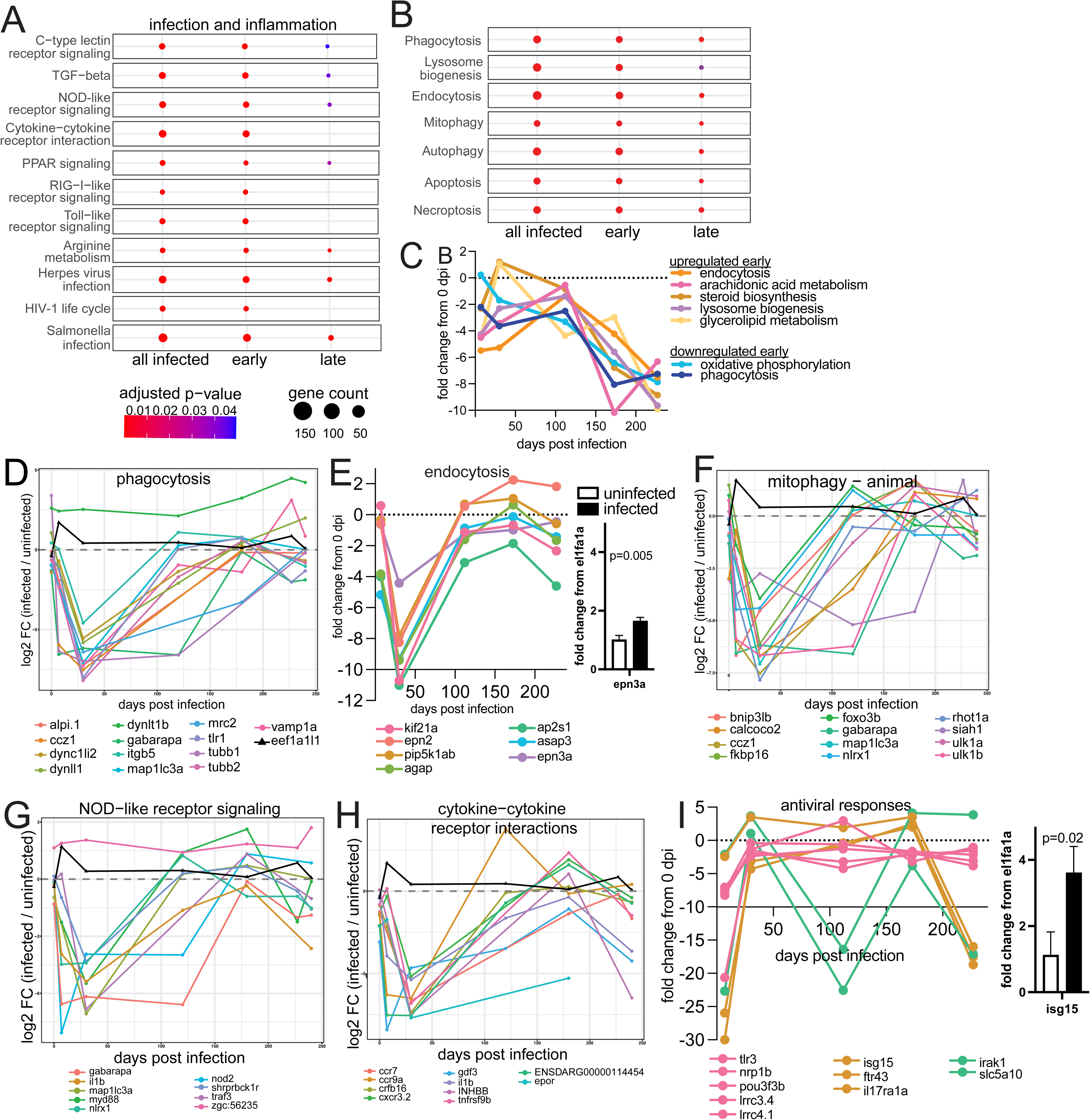
Infection and inflammation pathways are regulated in *M. leprae*-infected zebrafish. (A, B) KEGG analysis identified 103 pathways regulated by *M. leprae* infection in zebrafish, including those known to contribute to leprosy in humans (endocytosis, phagocytosis, arachidonic acid metabolism, necroptosis) and physiological pathways (calcium signaling, MAPK signaling, protein processing). (C) Regulation of *M. leprae*-induced pathways, which upregulated (endocytosis, arachidonic acid metabolism, steroid biosynthesis, lysosome biogenesis, and glycerolipid metabolism) or down regulated (oxidative phosphorylation, phagocytosis) in early infection. (C-I) Expression of DEGs in *M. leprae*-induced pathways, including phagocytosis (D), endocytosis (E), mitophagy (F), NOD-like receptor signaling (G), cytokine-cytokine receptor interactions (H), and antiviral responses (I). In E and I, pathway DEGs are detected by qPCR in uninfected zebrafish larvae or those infected with ∼100 CFU *M. marinum:PGL-1* for 4-5 days. E, expression of epn3a, an endocytosis DEG. I, expression of isg15, an antiviral DEG. Student’s t-test.

An examination of the expression of pathway genes reveals that gene subsets are similarly regulated during *M. leprae* infection. For example, many pathway genes are downregulated immediately after infection and remain downregulated through 30 dpi, before recovering expression in late infection. This was seen with phagocytosis and endocytosis genes (Figure 3d-e). To validate the regulation of these pathways, RNA from larval zebrafish infected with *M. marinum*:PGL-1 was used to confirm the regulation of selected pathway genes by qPCR. As with *M. leprae*, epn3a was regulated by *M. marinum:PGL-1* infection (Figure 3e). In contrast to endocytosis, some genes related to mitophagy and NOD-like receptor signaling displayed prolonged downregulation into late infection; this occurred with nod2, gabarapa, ulk1b and siah1 (Figure 3f-g). A third expression pattern was seen with some cytokine-cytokine receptor interaction genes, such as il1b, tnfrsf9b, and cxcr3.2, which recovered expression but were downregulated again through late infection (Figure 3h). Some antiviral signaling genes follow this late downregulation pattern, including isg15, il17ra1a and ftr43, while others maintained expression over time (tlr3, nrp1b, pou3f3b, lrrc3.4 and lrrc4.1), or were downregulated at ∼100 dpi (irak1, slc5a10). qPCR of *M. marinum*:PGL-1 infection confirmed that antiviral isg15 was upregulated 4-fold (p=0.02) (Figure 3i).

Beyond the infection and inflammation pathways, other physiological pathways were regulated that are less typically associated with leprosy. These include oxidative phosphorylation, and signaling through calcium, MAP kinase, Wnt, and mTOR (Figure 4a-b). Overtime, the number of regulated pathway genes decreased from early to late infection. Patterns of gene expression differed between pathways, and some regulated pathways featured subsets of genes with related expression patterns. For example, although genes involved in both positive and negative regulation of Wnt were downregulated early in infection, inhibitors of Wnt signaling recovered expression first, around 100 dpi. Positive regulators recovered months later, ∼173 dpi (Figure 4c). To confirm Wnt regulation in vivo, expression of THBS1a, an upstream regulator of Wnt signaling, was detected by qPCR in *M. marinum*:PGL-1-infected larvae (Figure 4c). In contrast to the Wnt pathway, genes that regulate TGF-β signaling were expressed in two patterns, with up-regulation starting in early (tgfb1a, THBS1a, grem1b) or late infection (smad6a, acvr1ba, nrros) (Figure 4d). Regulation of calcium signaling by *M. leprae* was validated with detection of ppp3ca in *M. marinum*:PGL-1 infection (Figure 4e). In contrast to most other pathways, many genes involved in mTOR signaling were upregulated in early infection, such as wnt5a, lamtor2, and rnf152 (Figure 4f). Finally, a pathway associated with intracellular infection by Salmonella bacteria was significantly regulated, with several genes upregulated in early infection (myl9b, myl10, and dynlt1b), and many others strongly downregulated (arpc4, cyfip2, il1b) (Figure 4g). While this pathway is not known to be associated with leprosy in humans, its regulation may indicate the use of similar defense strategies to combat intracellular bacteria.

**Figure 4.**
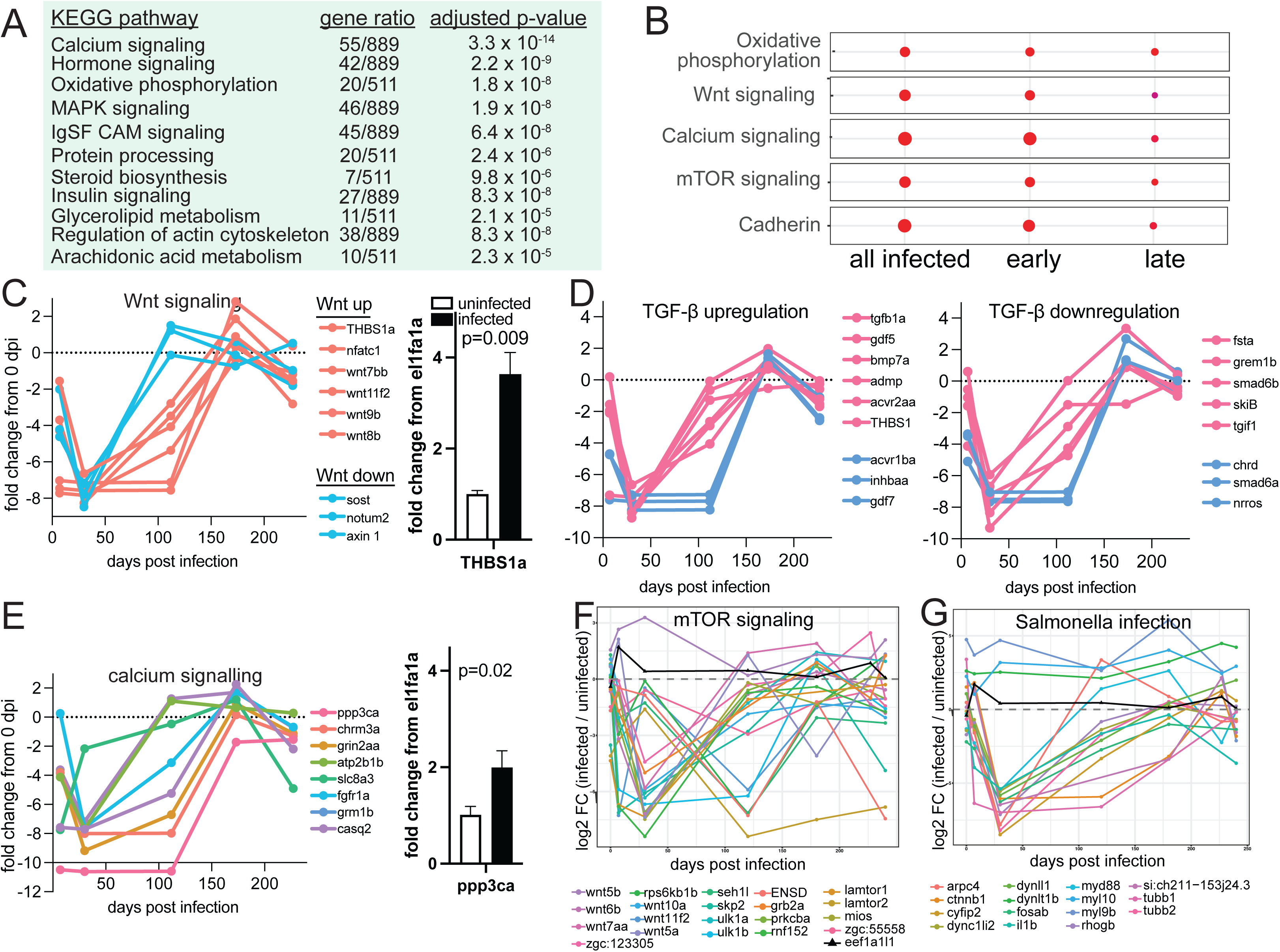
*M. leprae* infection regulates physiological pathways. (A) 11 physiological KEGG pathways that are induced by *M. leprae* in zebrafish but not typically associated with leprosy in humans, in all infected zebrafish compared to uninfected. (B) Physiological KEGG pathways are differentially regulated in early versus late infection. (C-G) Expression of infection-induced pathway DEGs in *M. leprae,* including Wnt signaling (C), regulation of TGF-β (D), calcium signaling (E), mTOR signaling (F), and Salmonella infection (G), as in Figure 3. In C and E, upstream pathway DEGs are detected by qPCR in *M. marinum:PGL-1*-infected zebrafish. C, expression of THBS1a, a Wnt signaling DEG. E, ppp3ca, associated with calcium signaling. N=11-30 larvae per group. Student’s t-test.

### Rag1 mutation alters the expression of genes associated with leprosy in humans

Given that T cells are likely key determinants of the human immune response to *M. leprae*, particularly in T-lep, we compared infected to uninfected rag1 mutants and heterozygotes (Figure 5a). The top regulated genes in each genotype were most associated with L-lep (40-43%), while downregulated genes were associated with R1 and R2. More leprosy-associated genes were regulated in heterozygotes (27) than mutants (6) (Figure 5b). 8 DEGs were regulated in heterozygotes but not in rag1 mutants but, suggesting that these genes required rag1 for wildtype expression (Figure 5c). Heatmaps were used to visualize the regulation of genes associated with each form of human leprosy in the rag1 heterozygotes and mutants, which revealed that rag1 mutation alters the regulation of genes associated with several forms of leprosy. Regulation of genes associated with multiple forms of leprosy (M) was curtailed in rag1 mutants (Figure 5d). In contrast, the downregulation of T-lep genes in rag1 heterozygotes was exacerbated in rag1 mutants (Figure 5d).

**Figure 5.**
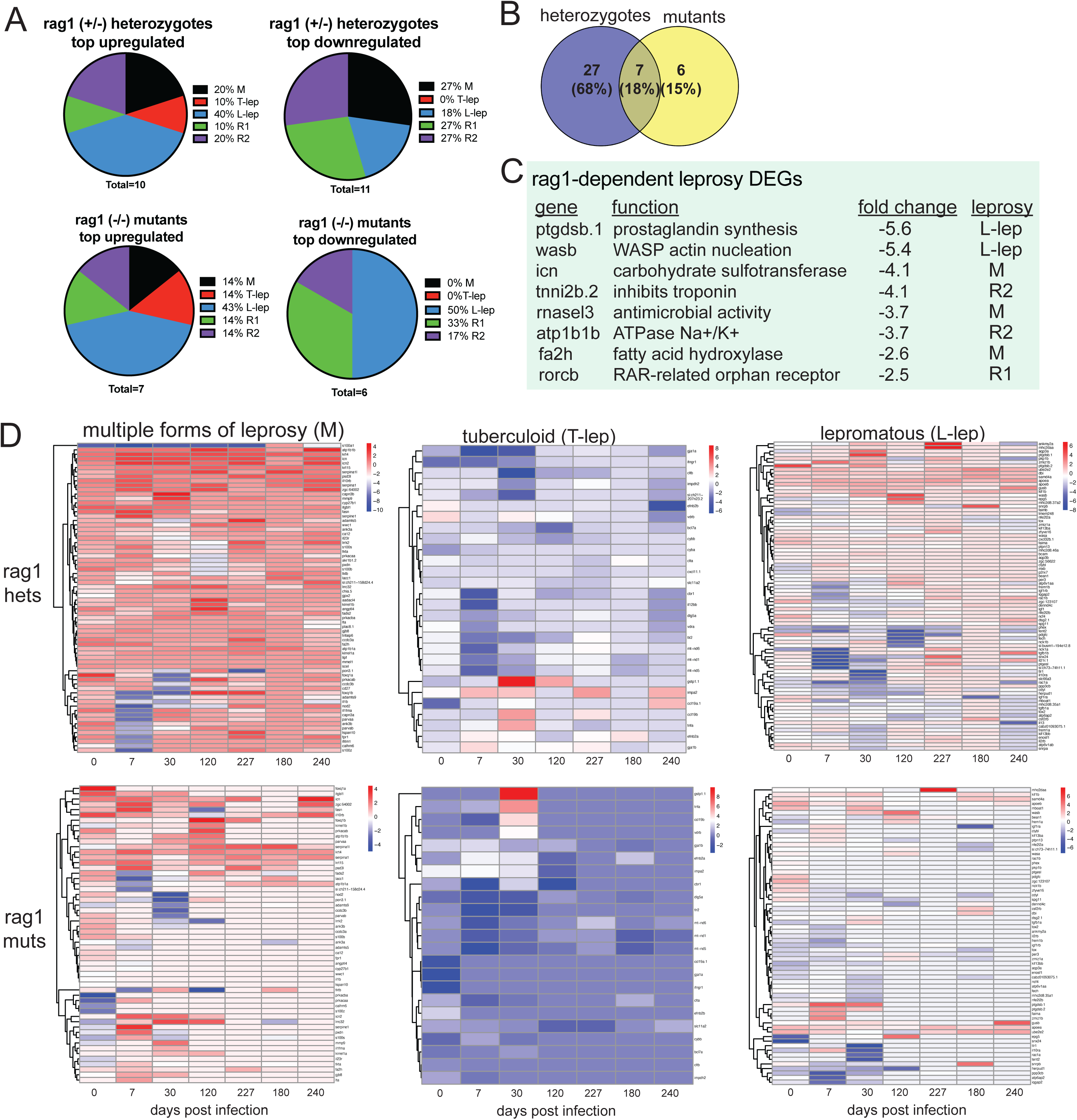
The regulation of leprosy genes decreases in rag1 mutant zebrafish. (A) The distribution of the top 10 upregulated and downregulated leprosy DEGs in rag1 (+/-) heterozygotes (top) and (-/-) mutants (bottom), comparing infected to uninfected. (B) There are 27 leprosy DEGs in rag1 (+/-) heterozygotes (purple), but most are absent in mutants (yellow). (C) Infection-induced leprosy DEGs absent in rag1 (-/-) mutants. (D) Heatmaps showing regulation of leprosy DEGs in infected compared to uninfected heterozygotes (top), or infected compared to uninfected mutants (bottom). Red is upregulated, blue is downregulated.

### Genes regulated by *M. leprae* infection influence the outcome of mycobacterial infection

To demonstrate *M. leprae*-regulated genes impact infection, zebrafish larvae with mutations in many genes were generated using multiplex CRISPR. 17 genes from the RNAseq dataset that were regulated early in infection were selected, including isg15, serpine1, ifit16, and tlr1 (TLR6) (Figure 6a). After confirming their regulation in *M. marinum:PGL-1* infection (Figure 6b), F0 CRISPR mutants (crispants) were generated by injecting zebrafish eggs with guide RNAs (gRNA). At 3 dpf, crispant and control larvae were infected with fluorescent *M. marinum:PGL-1,* and monitored for mortality and bacterial load by fluorescence imaging. Infected triple crispants lacking isg15, ifit16, and trl1 (TLR6) had decreased survival compared to controls (Figure 6c). Single mutants were generated, which revealed that isg15 mutation was associated with decreased survival (Figure 6d). Repeating this approach identified a similar increase in mortality in infected serpine1 crispants (Figure 6e), although paradoxically, the bacterial load was decreased, as shown by fluorescence imaging (Figure 6f).

**Figure 6.**
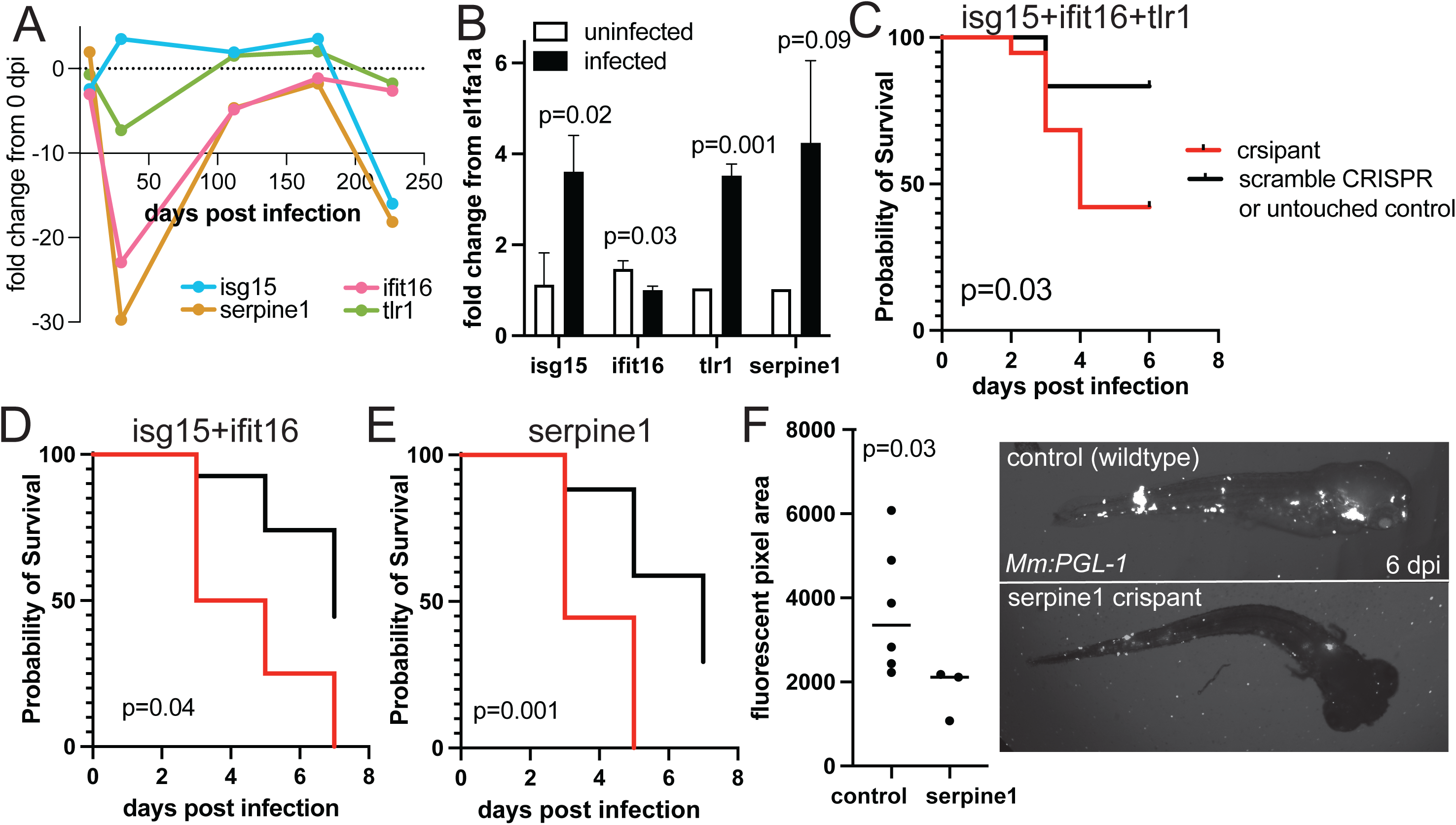
Multiplex CRISPR identifies *M. leprae*-induced genes that alter infection outcomes. (A) Expression of DEGs targeted by CRISPR in the *M. leprae* RNAseq data, and (B) qPCR detection of the gene transcripts in zebrafish infected with *M. marinum:PGL-*1 as in Figure 1. (C-E) Survival curves of crispant zebrafish (red) compared to a wildtype control (black), either a scramble CRISPR or untouched larvae. (F) crispants and wildtype larvae (red) were infected with fluorescent *M. marinum:PGL-*1 as described, and bacterial load was quantified by measuring fluorescence pixel area. Representative images at 6 dpi of infected larvae. Student’s t-test.

## Discussion

We showed that *M. leprae* infection of adult zebrafish mimics many transcriptional responses of humans to *M. leprae*, including the genes typically associated with T-lep, L-lep, R1 and R2. Transcription evolves over time, from 0 to 240 dpi, defining an early response featuring many L-lep genes, versus a late response that is dominated by L-lep and R1 genes. Over 240 days, T-lep genes are increasingly suppressed. Regulation of known and previously-unknown leprosy-associated genes was confirmed with *M. marinum:PGL1* infection. The larval system demonstrates many genes can be simultaneously screened using multiplex CRISPR to identify mutants with infection phenotypes.

Comparing the RNAseq zebrafish data to *M. leprae* viability, the immune response can be framed as an early response to viable *M. leprae* (0-30 dpi), and a late response to nonviable *M. leprae* (180-240 dpi). Given that viable *M. leprae* become non-viable over the course of infection, it is possible that early infection is typified by an inefficient immune response, while bacterial killing in later infection may be associated with an efficient immune response. Despite efficient *M. leprae* killing, paradoxically, the zebrafish response most resembles L-lep, which is typically linked to inefficient cell-mediated immunity and high bacterial load.

No prior study has tracked the host transcriptional response to *M. leprae* over a months-long chronic infection in a living vertebrate. Most existing work examines host responses over a few days, or bacterial transcription. Our data reveals that initial bacterial killing is independent of rag1, suggesting that innate immunity alone is capable of controlling *M. leprae* burden. However, the transcriptional signatures that resemble human leprosy forms is curtailed in rag1(-/-) mutants, suggesting that adaptive immunity shapes the character of the immune response more than it drives bacterial killing per se.

As in zebrafish, ISG15 was highly induced in human macrophages, by the type I interferon response triggered by *M. leprae* infection^13^. This points to the same interferon-driven response being conserved from fish to humans, suggesting that type I interferon a core piece of the antimycobacterial response in leprosy, and likely to other mycobacterial pathogens. Serpine1 (PAI-1) is a canonical downstream effector of TGF-β signaling that promotes fibrosis by suppressing matrix metalloproteinase activity, and that we identified both TGF-β pathway regulation and serpine1 induction in our dataset. We speculate that the zebrafish transcriptional response to *M. leprae* may reflect early and deleterious activation of the same fibrotic program that ultimately underlies irreversible nerve damage in human leprosy^14-16^.

In sum, our data demonstrate the utility of the zebrafish model to identify known and previously-unknown mediators of pathogenesis in *M. leprae* infection. This approach may advance the understanding of how immune responses to *M. leprae* are initiated and evolve over the course of chronic infection. Genes that are regulated in response to *M. leprae* may be useful as new targets for host-directed therapies, which could be tailored to combat pathogenesis in acute versus chronic infection. In the future, combining transcriptional analysis with genetic and pharmacological screening in zebrafish could provide a rapid means of identifying genes with in vivo phenotypes in *M. leprae* infection, immune responses, and neurological injury.

## Supplemental Figures

**Supplemental Figure 1:**
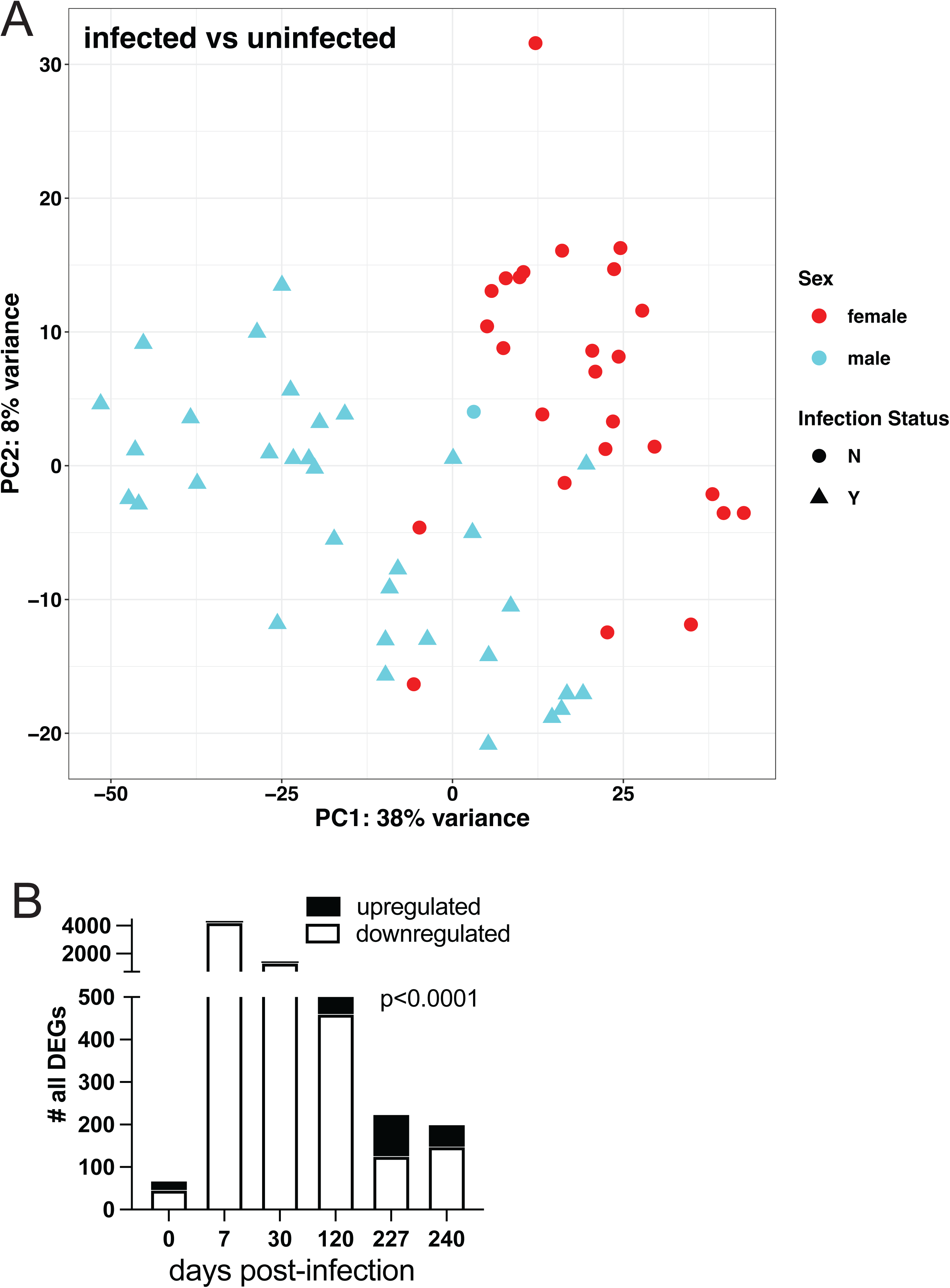
PCA scatter plot showing variance due to zebrafish sex. (A) PCA plot. (B) Up- and down-regulated genes over time. Chi squared.

**Supplemental Figure 2:**
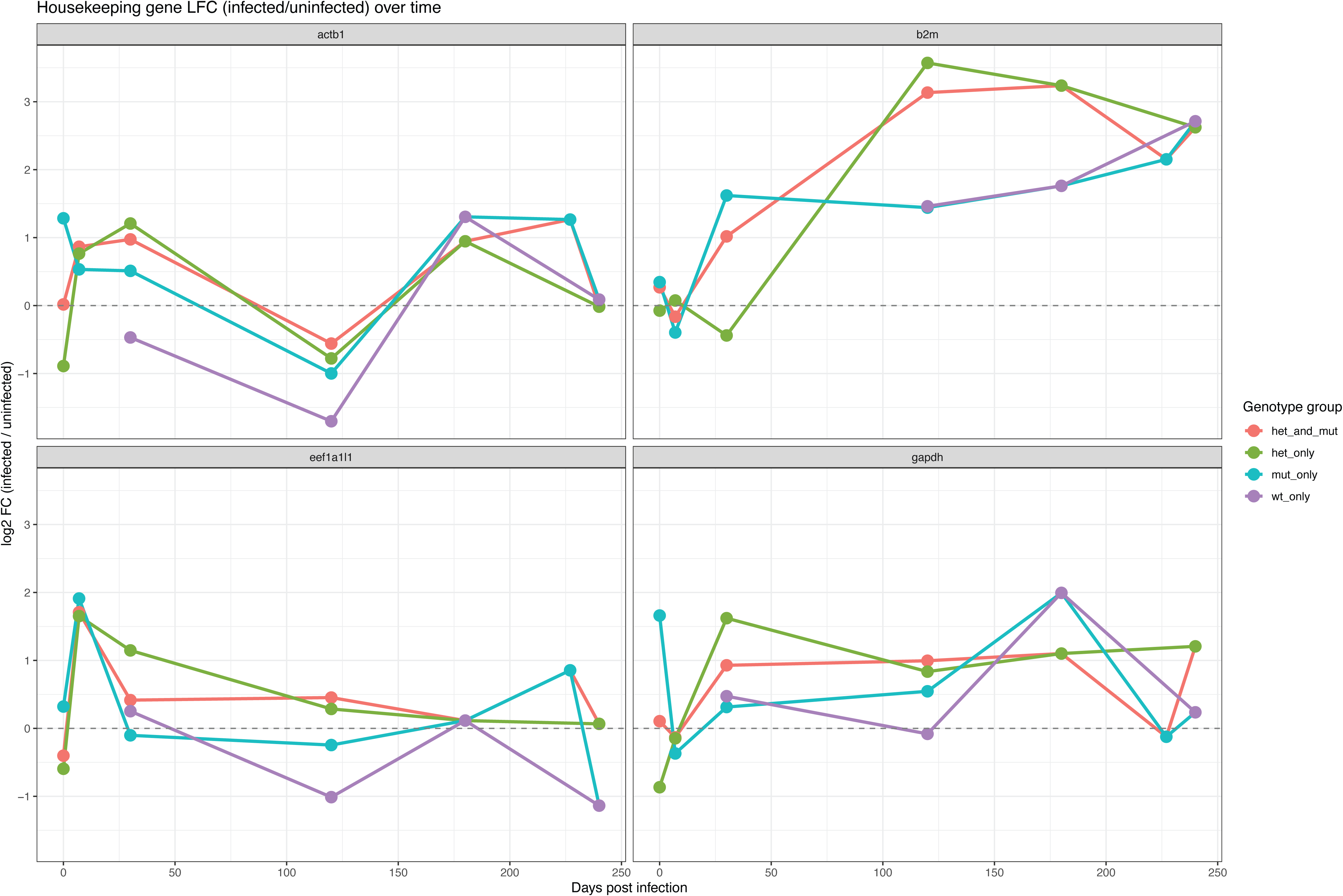
Expression trajectories of typical housekeeping genes (actb1, b2m, eef1l1, and gapdh) over the infection time course.

**Supplemental Figure 3:** Crispants lacking tlr1 survive infection as well as wildtype larvae.

## Supplemental Tables

**Table 1:** Details of the *M. leprae*-infected zebrafish experiment

**Table 2:** Human leprosy-associated genes and their zebrafish orthologs

**Table 3:** DEGs regulated by *M. leprae* infection (all infected compared to all uninfected)

**Table 4:** DEGs regulated by *M. leprae* infection (infected compared to 0 dpi)

**Table 5:** DEGs regulated by *M. leprae* infection of rag1 (+/-) heterozygotes

**Table 6:** DEGs regulated by *M. leprae* infection of rag1 (-/-) mutants

**Table 7:** Pathways regulated by infection (all infected compared to all uninfected)

**Table 8:** R code used in this study

## METHODS

### Zebrafish husbandry

Fertilized embryos of the wildtype AB genotype were placed into a petri dish and randomly selected for infection. During the duration of the study, larvae were housed at 28.5°C, in an excess of zebrafish water containing ddH2O, 14.6 g/l sodium chloride (JT Baker #3628-F7), 0.63 g/l potassium chloride (Sigma-Aldrich #P3911), 1.83 g/l calcium chloride (G-Biosciences #RC-030), 1.99 g/l magnesium sulfate heptahydrate (MP Biomedicals #194833), methylene blue chloride (Millipore Sigma #284), and 0.003% 1-phenyl-2-thiourea (PTU, Sigma-Aldrich #189235) to prevent melanocyte development. Larvae were anesthetized with 2.8% Syncaine (Syndel #886-86-2) prior infection or physical manipulation. Study groups of approximately 30 larvae were infected, for a total of ∼100 larvae per experiment, totaling ∼600 larvae for the study. Zebrafish husbandry and experiments were conducted in compliance with guidelines from the U.S. National Institutes of Health and approved by the University of California San Diego Institutional Animal Care and Use Committee (#S18135) and the Institutional Biosafety Committee of the University of California San Diego (#2452). All zebrafish work was performed by lab personnel that were trained and supervised by a University of California San Diego Animal Care Staff supervisor.

### Larval zebrafish infections

Larvae are infected by injection of 10 nL into the caudal vein at 2-3 days post fertilization (dpf) using a capillary needle containing bacteria diluted in PBS + 2% phenol red (Sigma #P3532). After caudal vein injections, the same needle was used to inject onto 7H10 (Sigma-Aldrich # M199) agar plates containing 50 μg/mL hygromycin B (ThermoFisher #10687010) or 50 μg/ml kanamycin (TCI #K0047) in triplicate to determine the CFUs in the inoculum. ∼ 100−500 CFUs of *M. marinum:PGL-1* were administered to the larvae for experiments unless otherwise specified. Adult zebrafish were infected as described^6^.

### Microscopy and image analysis

To determine bacterial fluorescent pixel count, larval groups were placed n=1 per well of a 96-well imaging plate and placed on ice for approximately 45 minutes for anesthetization. Once anesthetized, standard hand soap was spread on the bottom of the 96 well plate to prevent condensation and larvae were manipulated to lay on their side, horizontal to the imaging plane.

Wide-field imaging was done using the Nikon Eclipse Ti2 inverted microscope with a 2X objective and Chroma 49008 ET-mCherry, Tx Red or SMZ-25/18 P2-EFLC EGFP BP HC filter sets. NIS-Elements (v. 5.20.01) software was used to batch capture images in both greyscale and the specified fluorescent channel (mCherry or GFP as determined by the injected bacteria’s fluorescence). Fluorescent images were taken at 0-7 days post infection (dpi) unless otherwise stated with larvae returned to 28.5 °C between imaging sessions. Captured images were then processed using Adobe Photoshop to mask both non-bacterial larval autofluorescence and bacterial colonies in the zebrafish water outside of the body. Total fluorescent pixel count for each larva was determined using macros in FIJI/Image J. Statistics were analyzed using Prism 11 (GraphPad).

### Transcriptomics

RNAseq data analysis was conducted in R using the DESeq2 package (v1.52.0) [1]. To evaluate temporal dynamics, multiple comparisons were executed within the generalized model framework by contrasting post-infection time points back to the day post-infection 0 (DPI 0) baseline controls. For these time-course analyses, genes were defined as statistically significant if they met a Benjamini-Hochberg adjusted p-value (padj) < 0.05 and an absolute log2 fold change (|log2FC|) >= 0.5. For the infected versus uninfected baseline and genotype comparisons, a significance threshold of padj < 0.10 and a |log2FC| >= 0.5 was applied to classify genes as upregulated, downregulated, or non-significant. To account for potential confounding biological variables, an adjusted log2FC was calculated by removing sex-specific differentially expressed genes, utilizing comparisons between uninfected male and uninfected female samples at the same timepoint. Downstream functional enrichment was conducted to identify regulated biological pathways using Gene Ontology (GO) terms, the Reactome knowledgebase, and Kyoto Encyclopedia of Genes and Genomes (KEGG) pathway analysis mapped to the Danio rerio reference annotation [2] [3] [4]. Gene functions were pulled from the UniProt Knowledgebase [5]. All statistical plotting and data visualizations were generated using the ggplot2 package (v4.0.3) in R [6].

### qPCR

Quantitative PCR (qPCR) was performed using the AzuraView Green qPCR Mix (Azura Genomics, Cat. No. AZ-2320) according to the manufacturer’s instructions. All reactions were executed on a CFX Opus 384 Real-Time PCR System (Bio-Rad). Target-specific primers were designed using the IDT PrimerQuest Tool and synthesized by Integrated DNA Technologies (IDT) [7]. Each 10 uL reaction consisted of 1x AzuraView Green qPCR Mix, forward and reverse primers at a final concentration of 10 uM each, and 1 uL of cDNA template (yielding a final mass of 50 ng of cDNA per reaction).

---

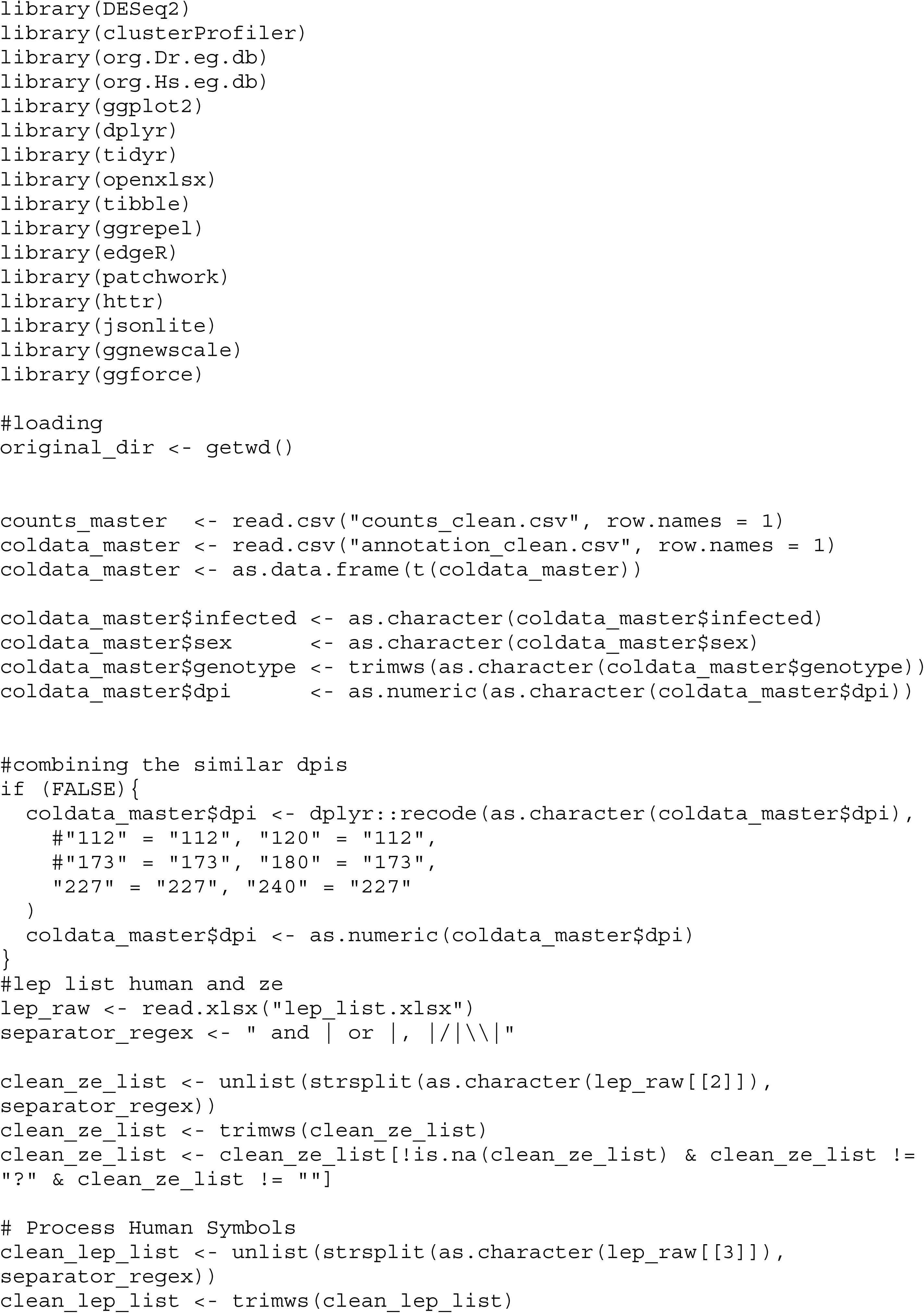

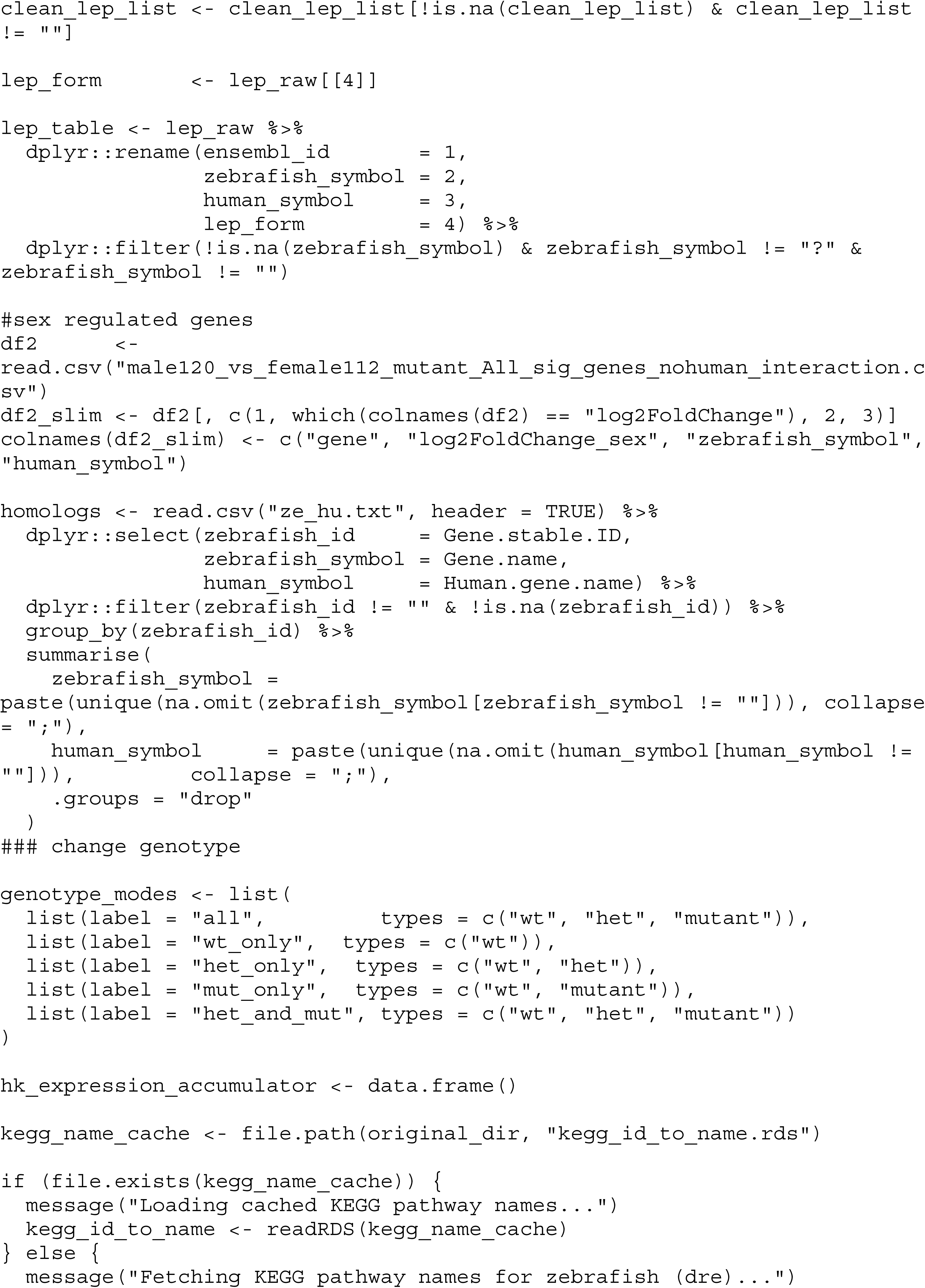

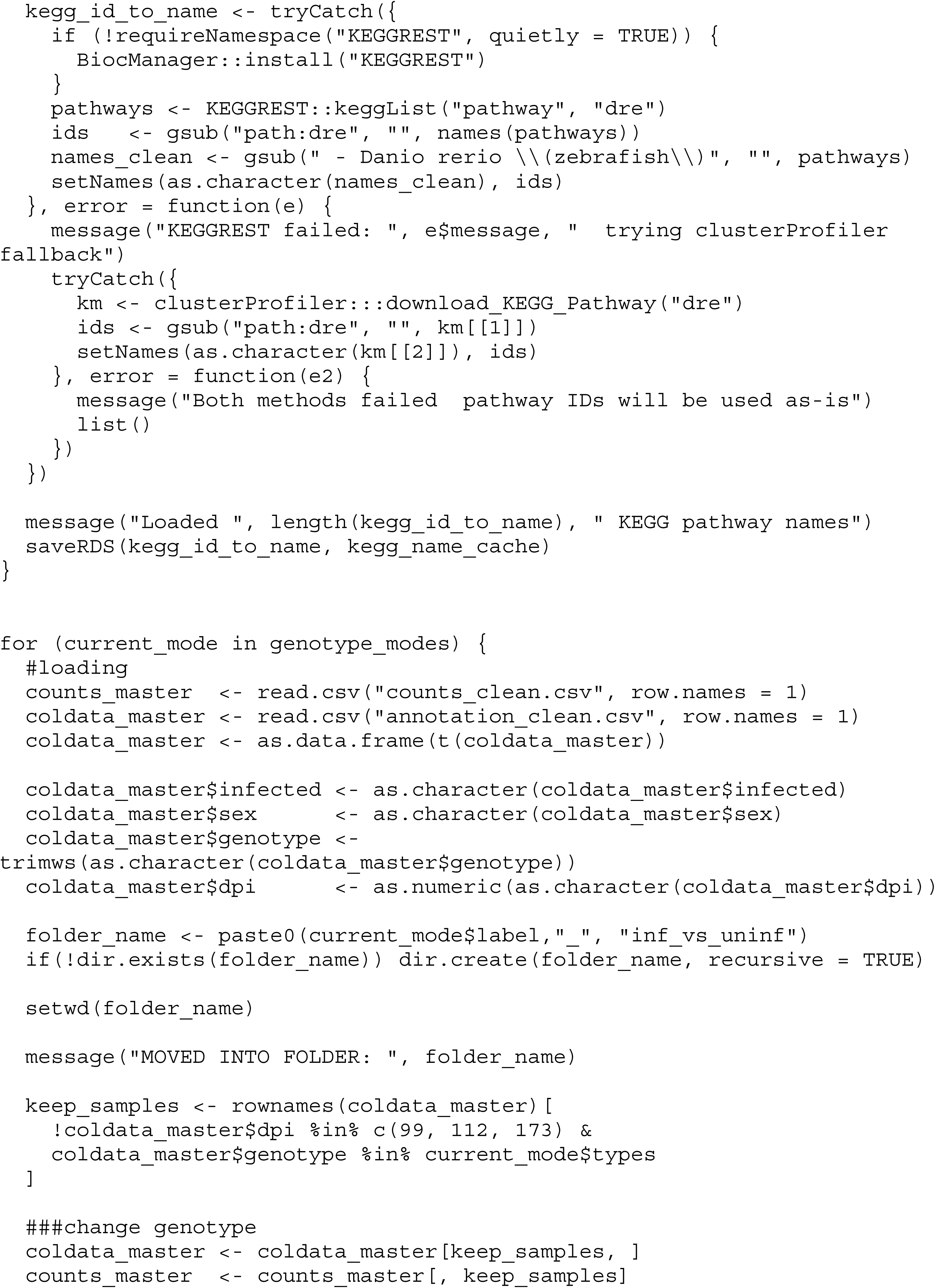

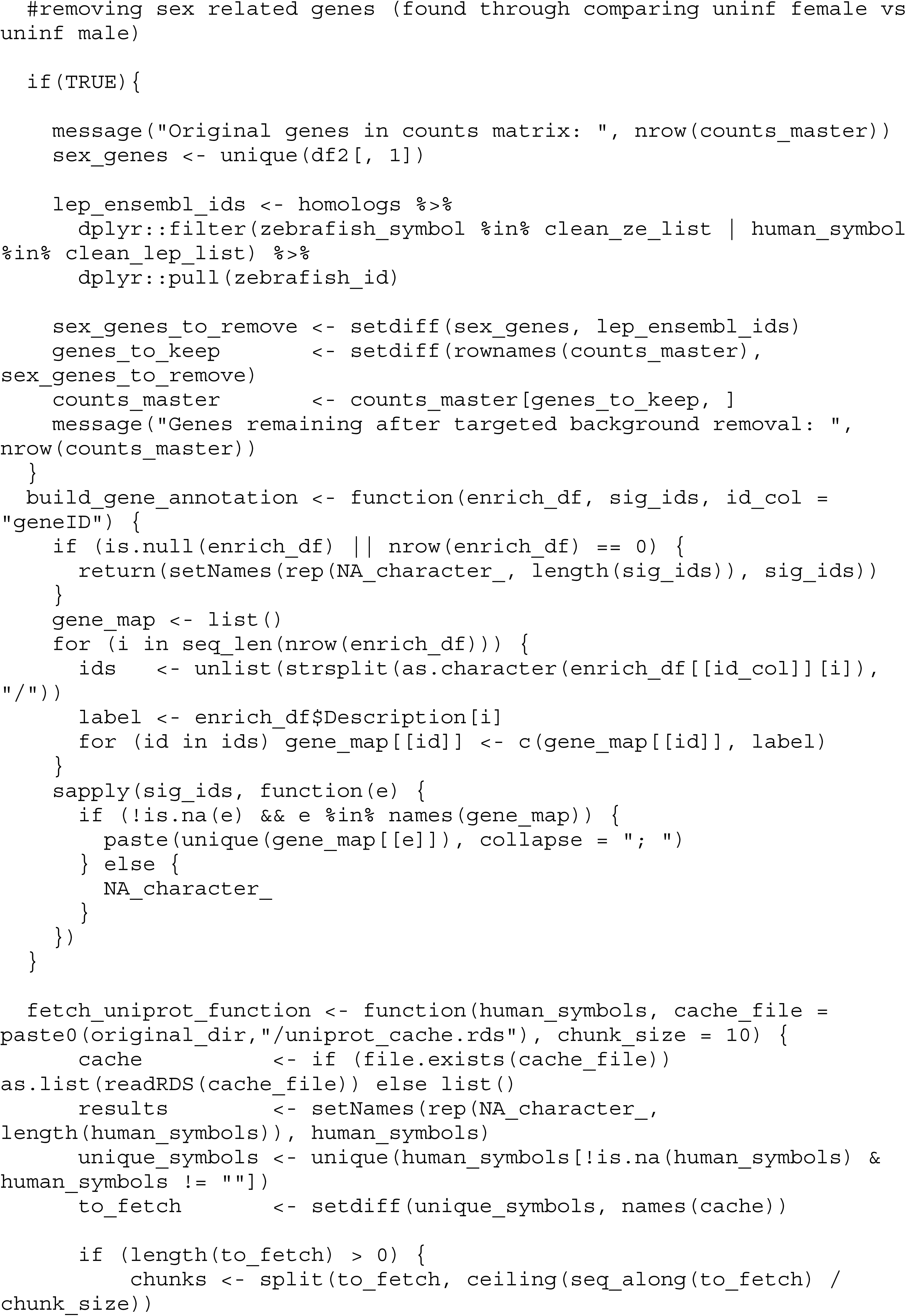

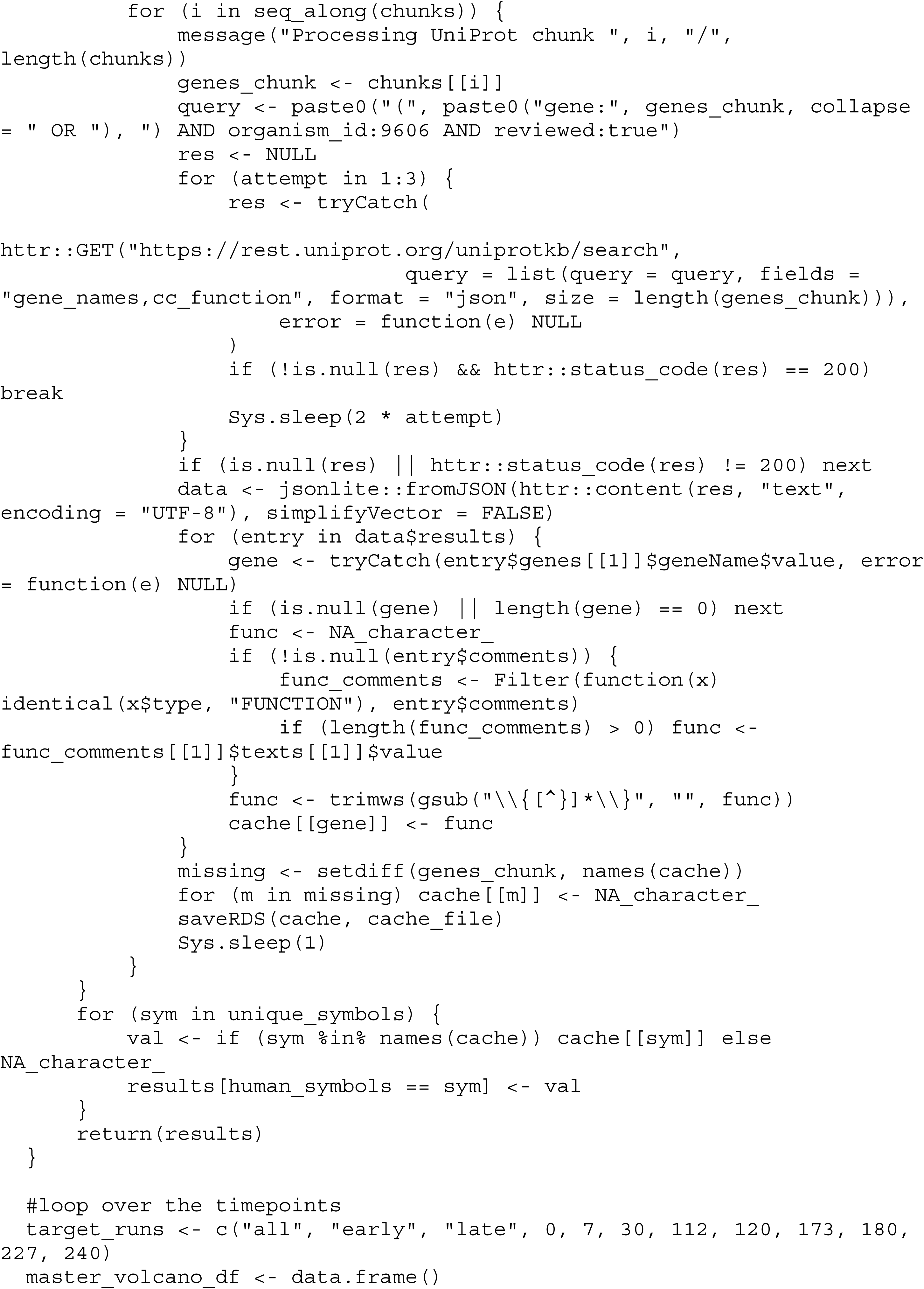

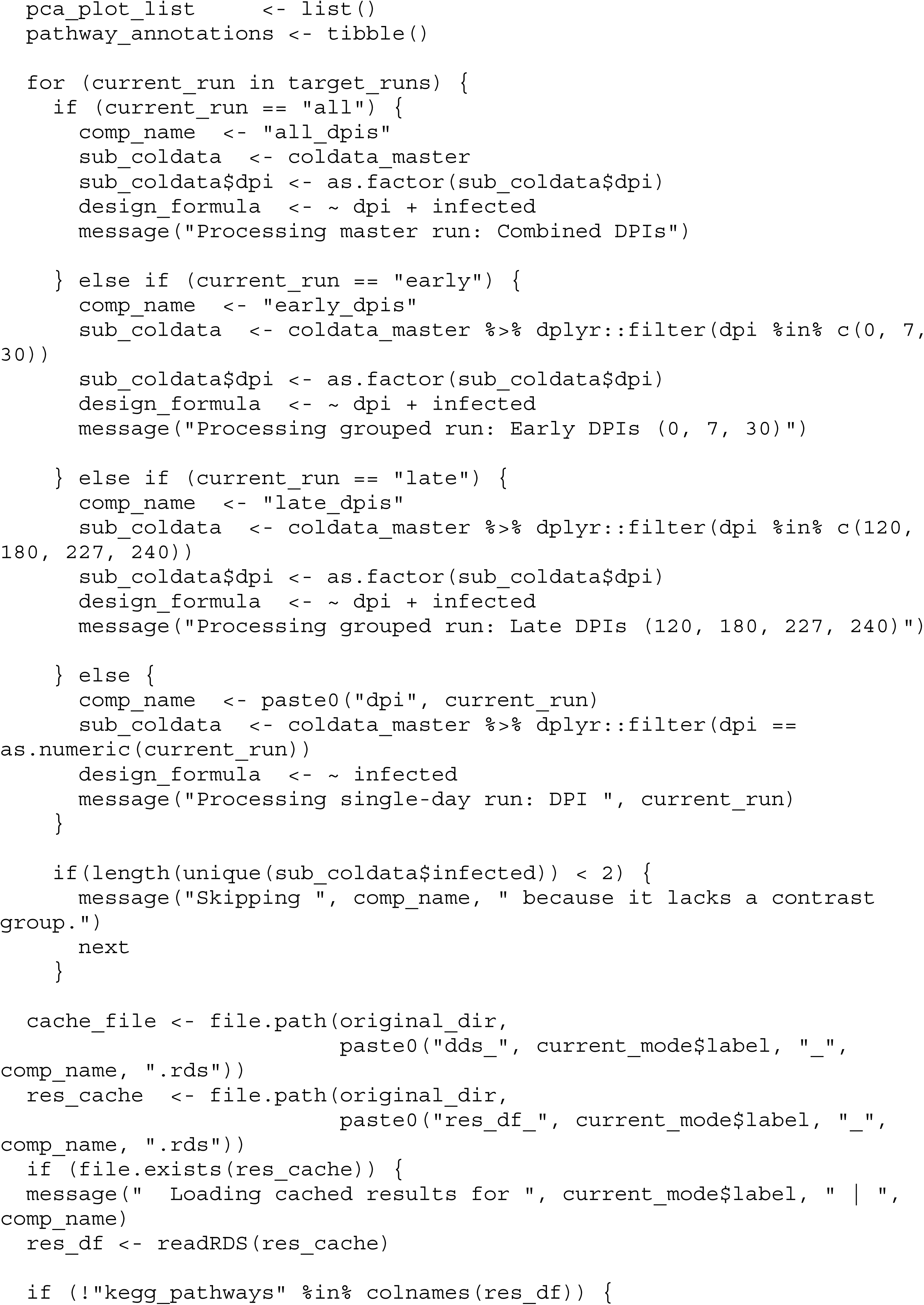

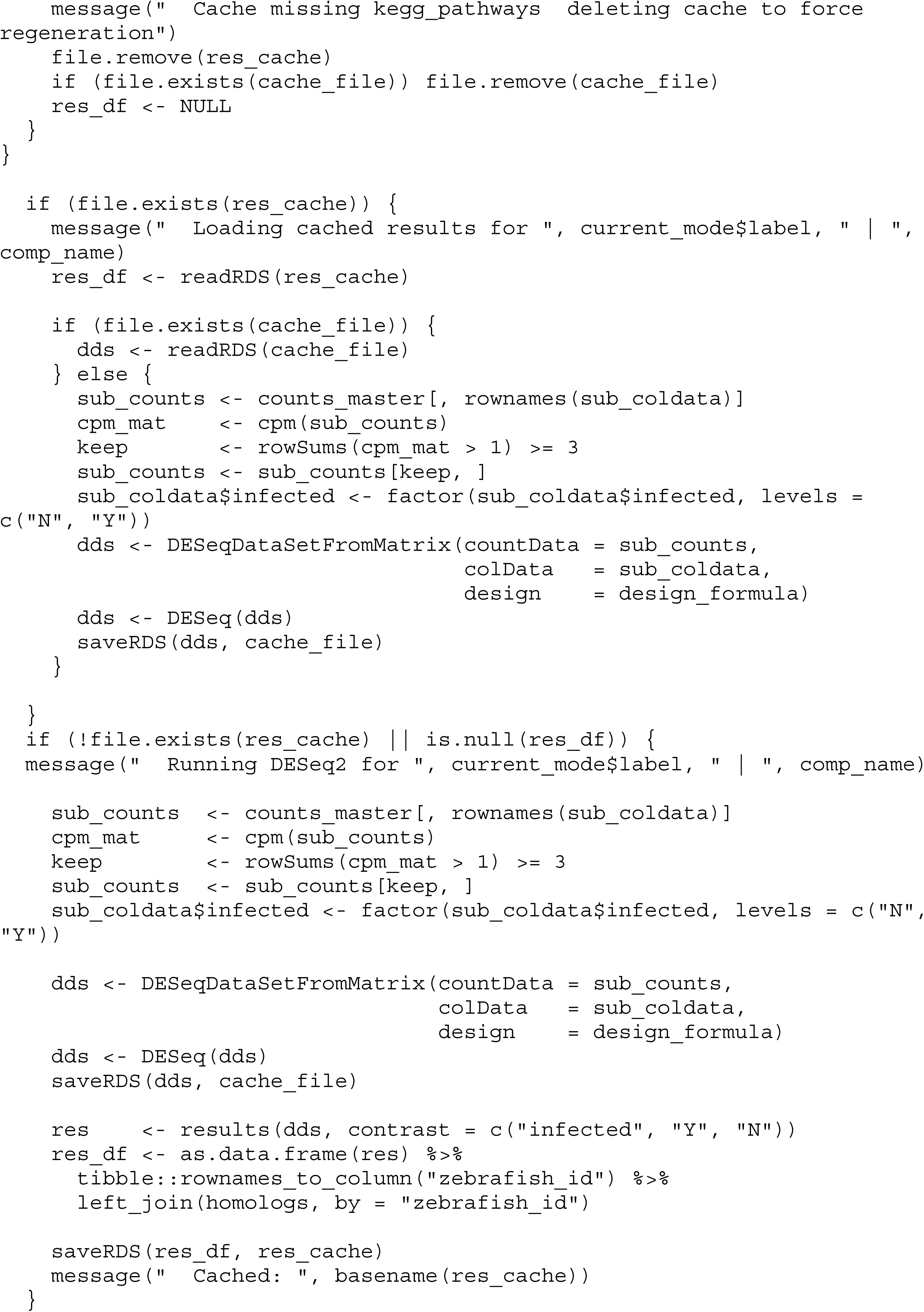

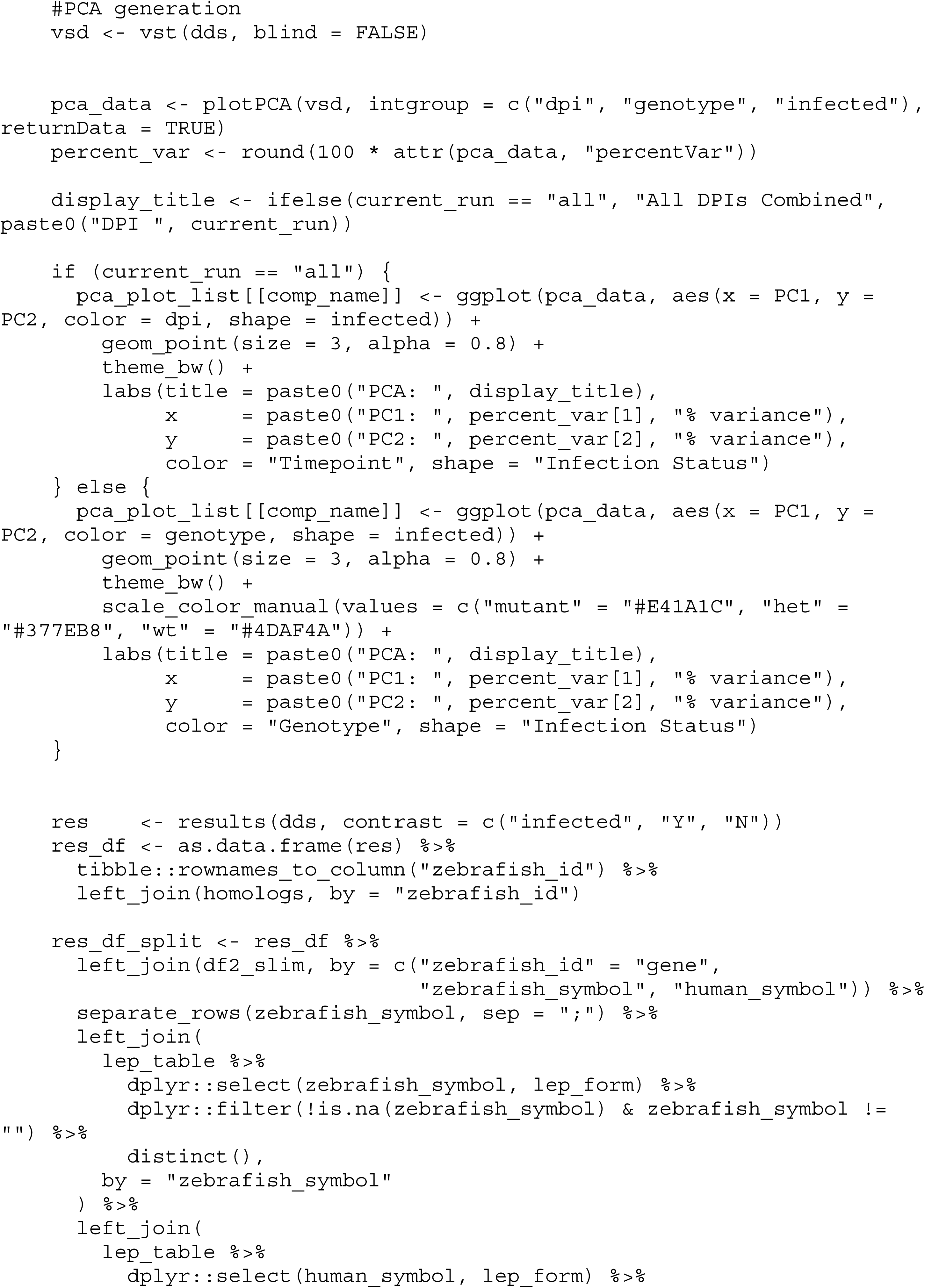

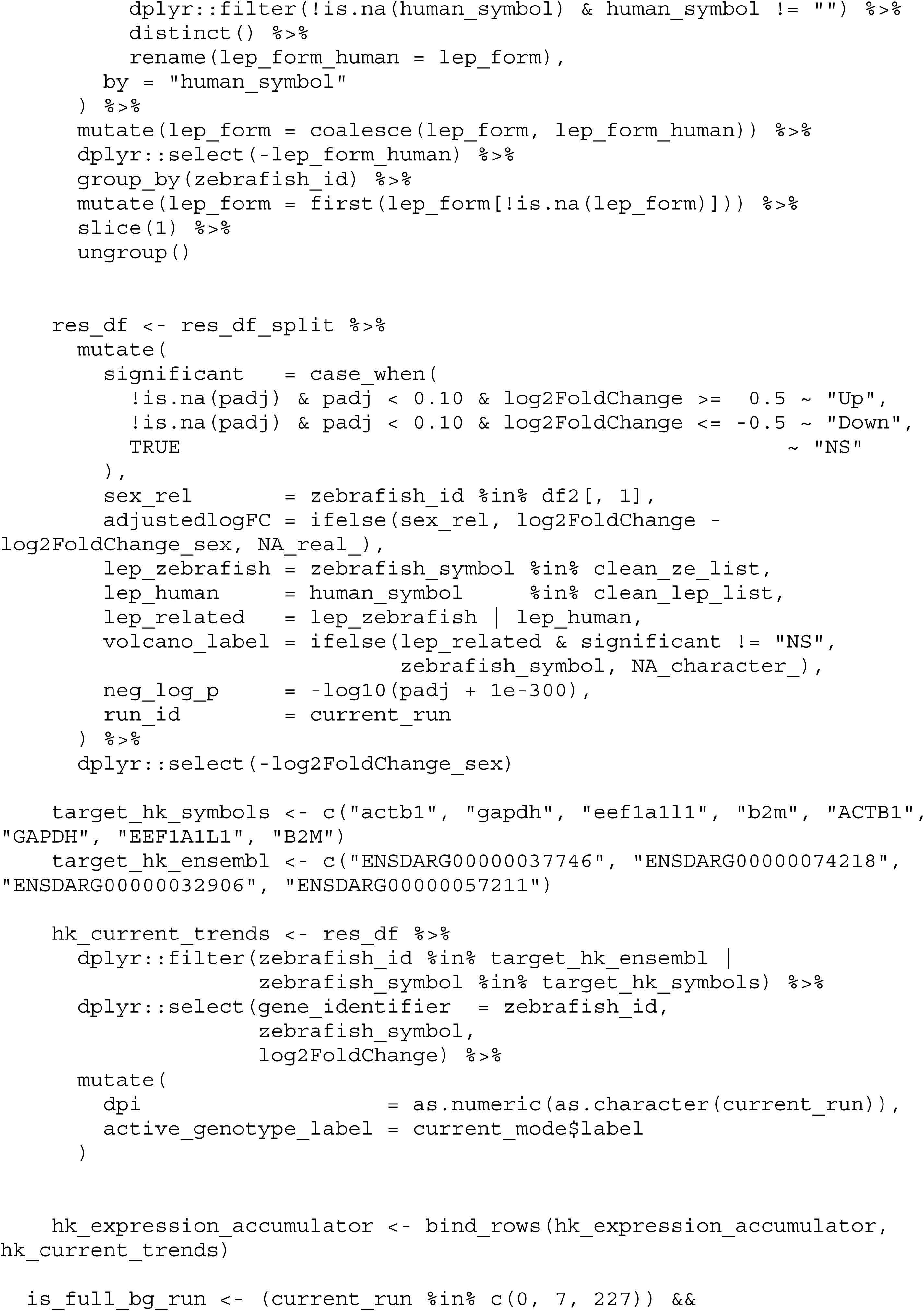

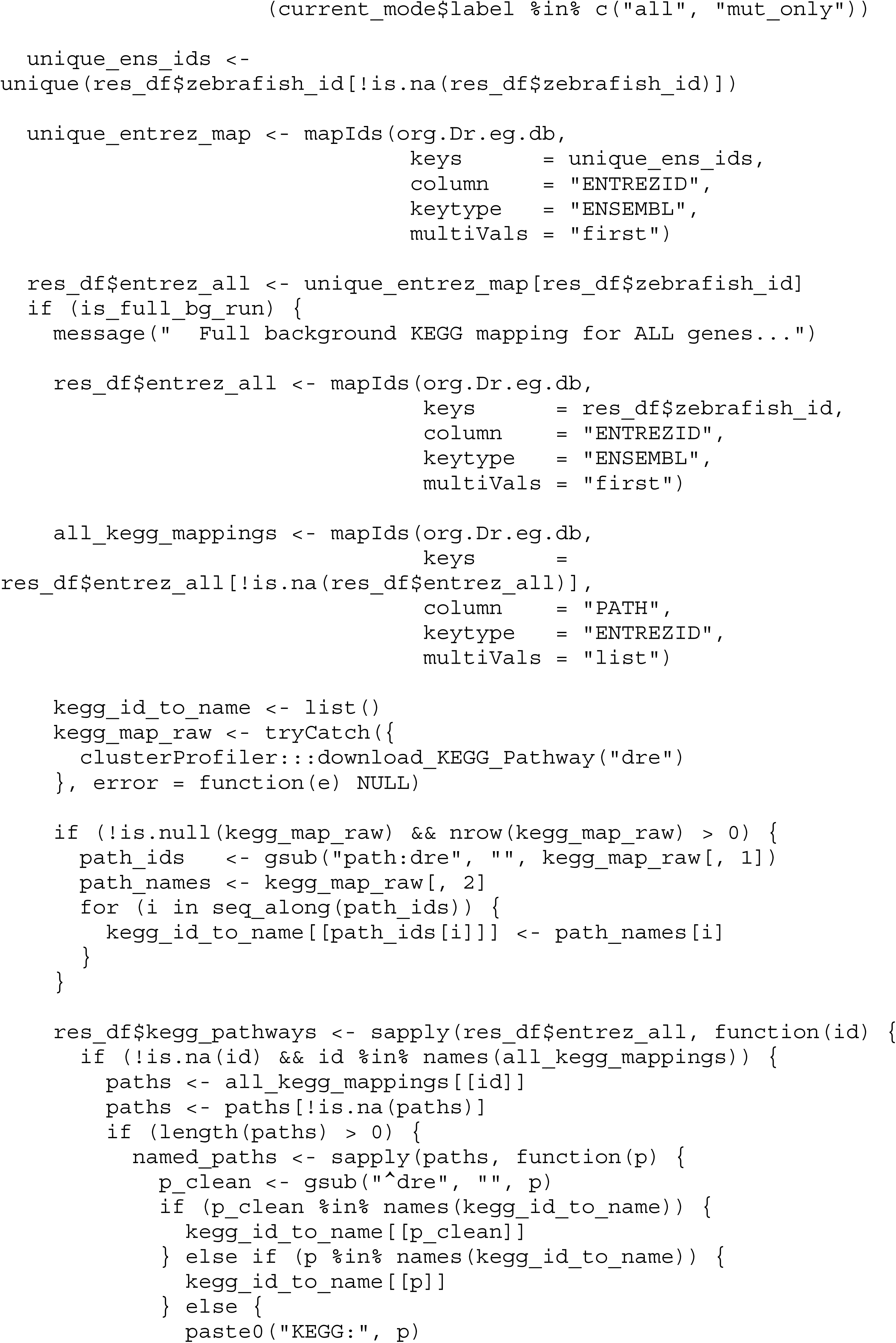

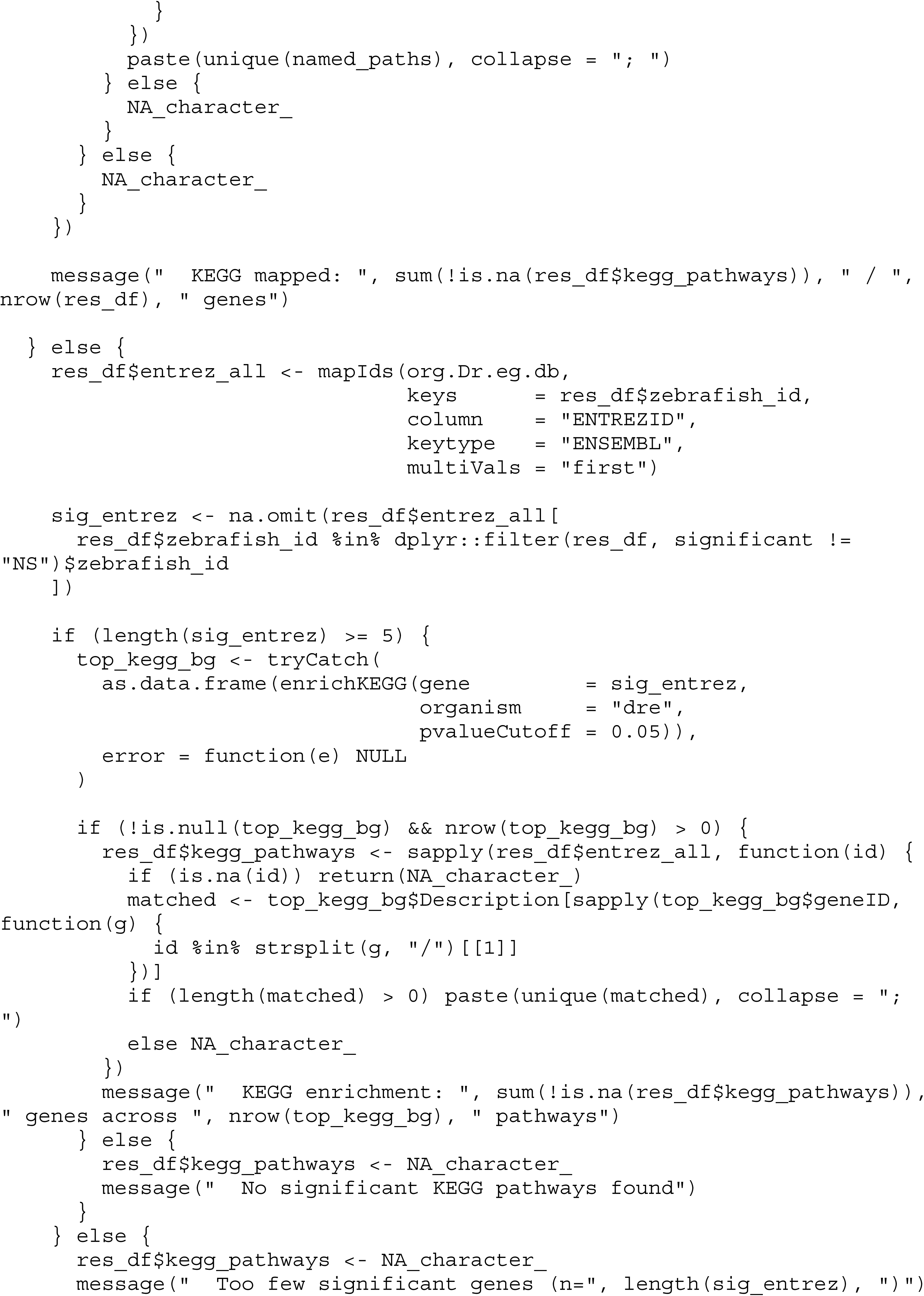

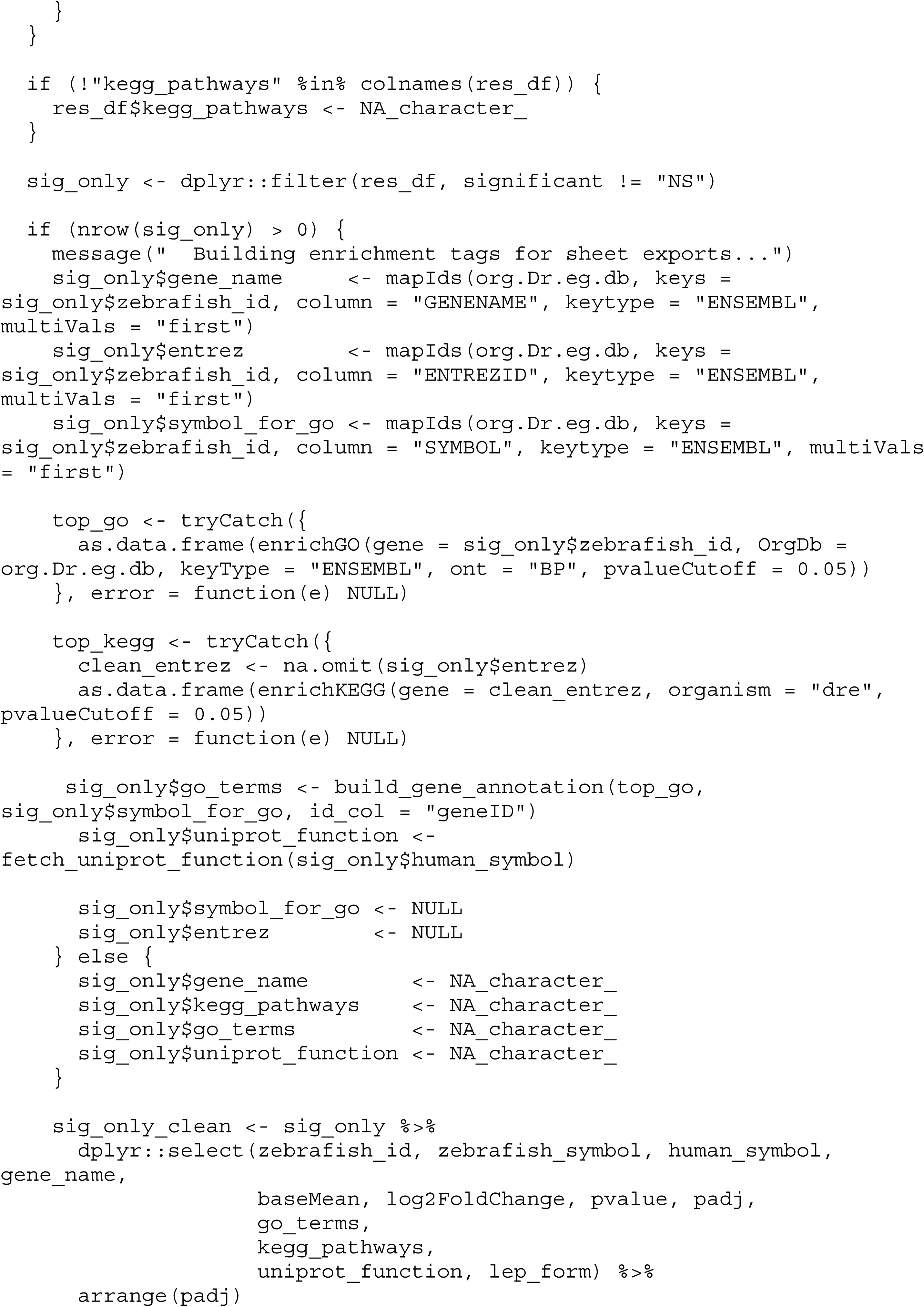

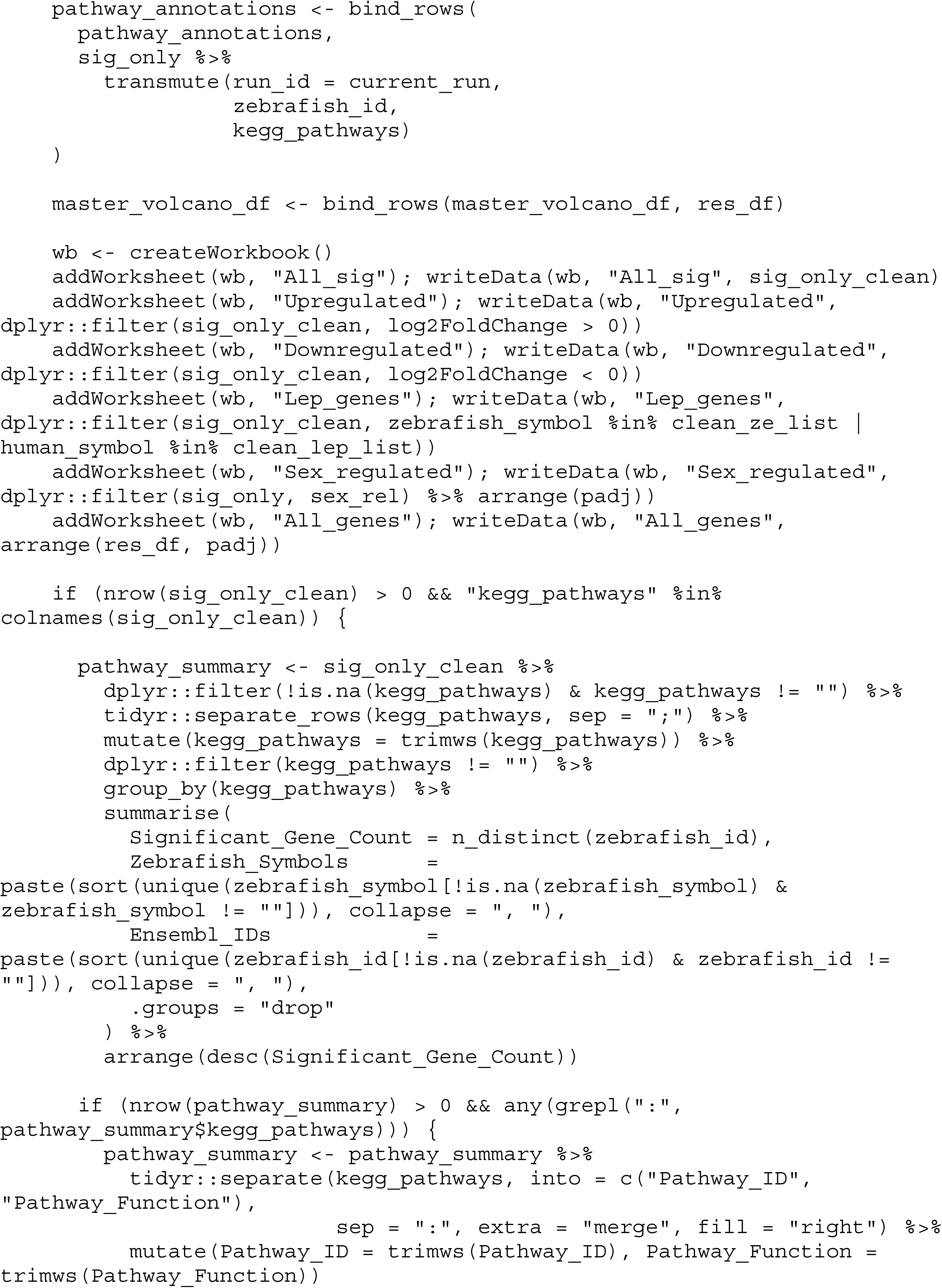

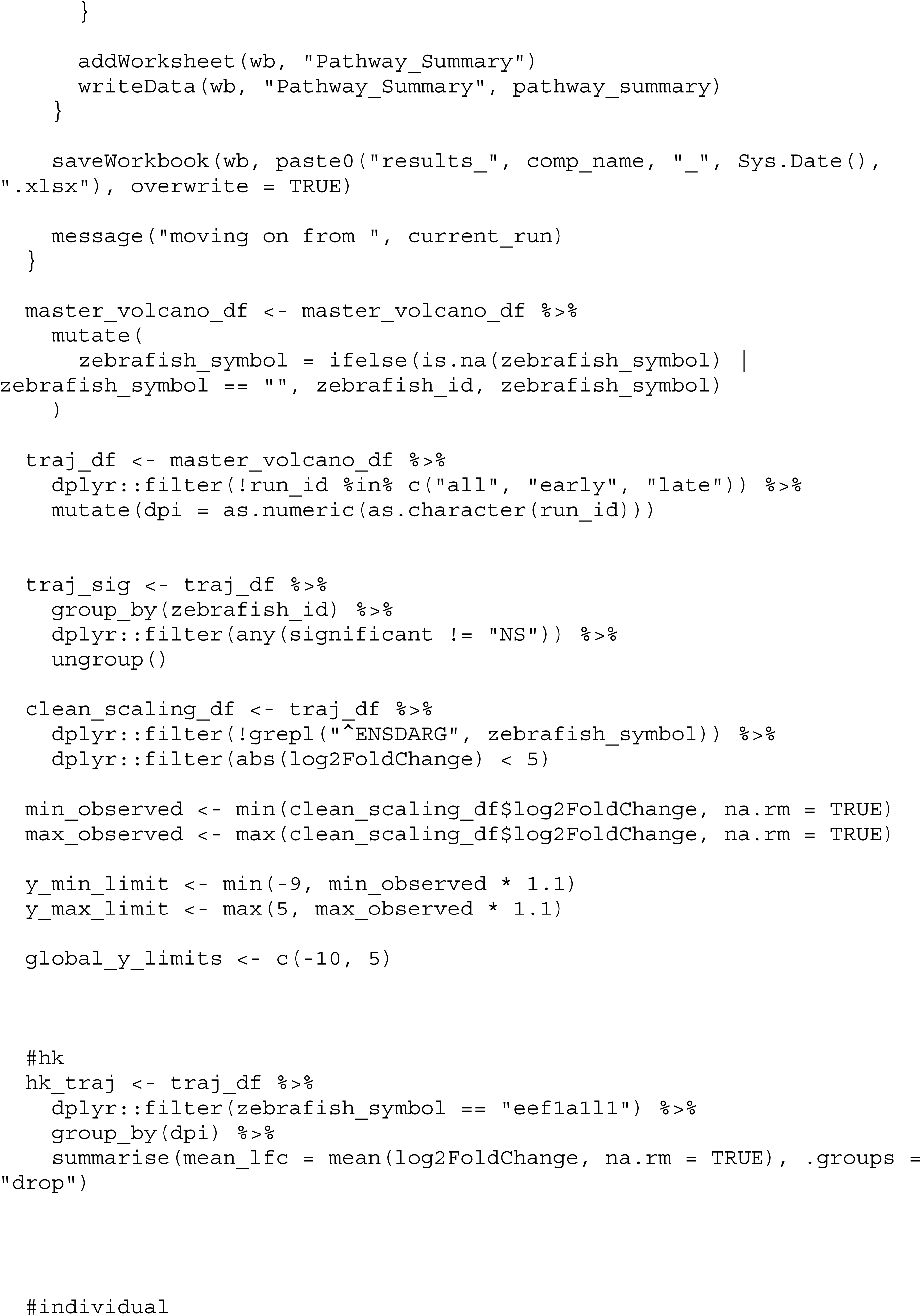

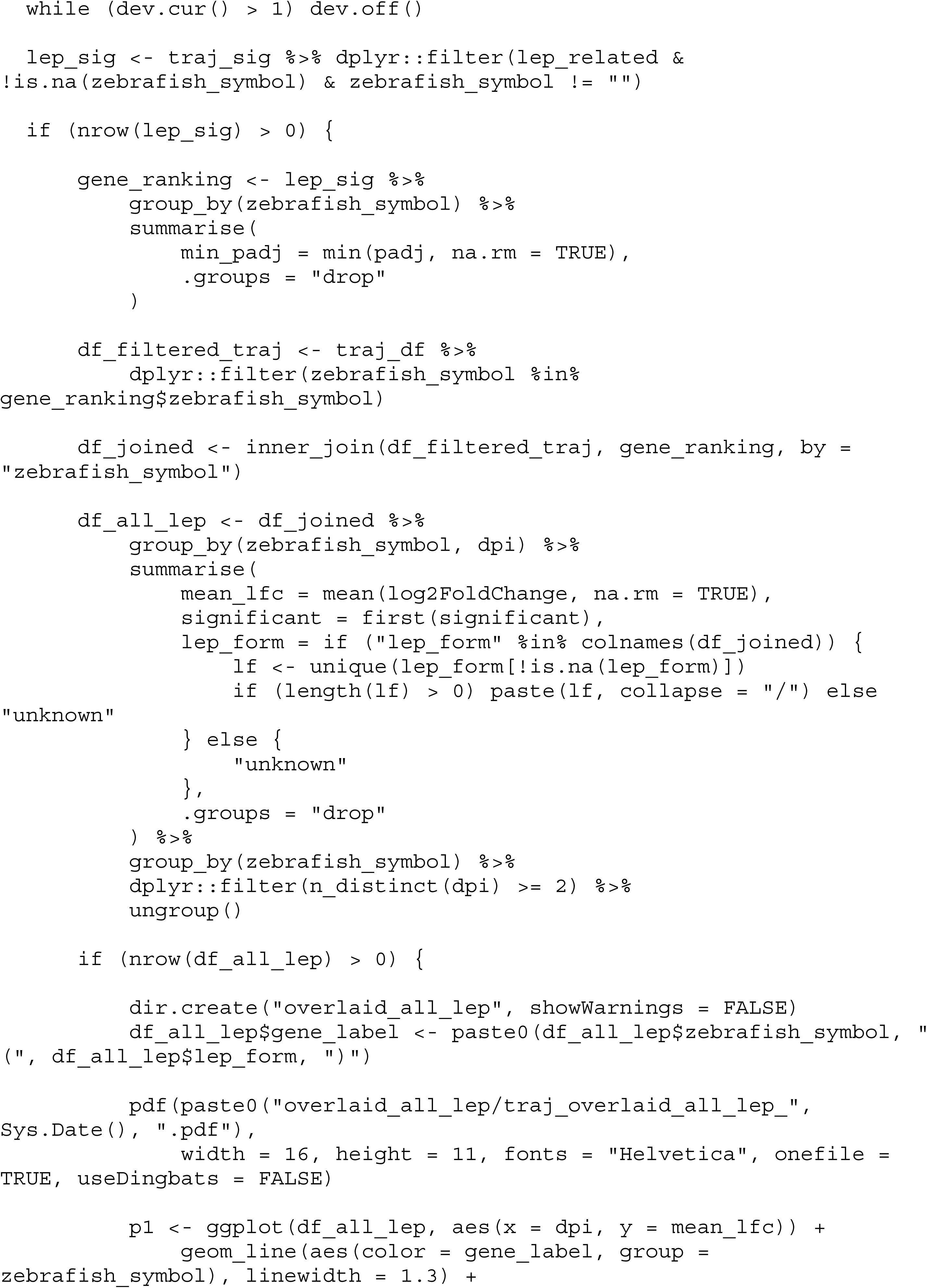

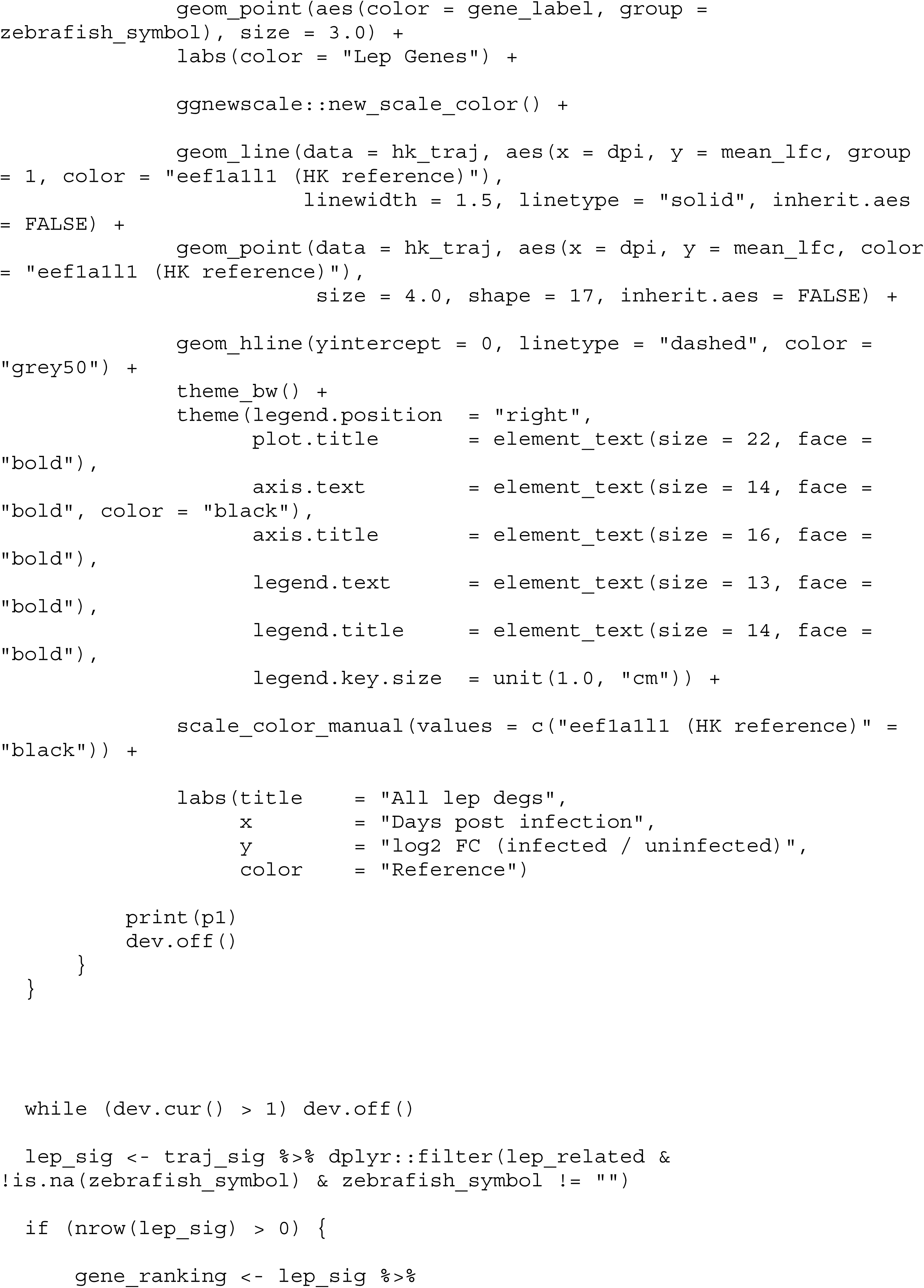

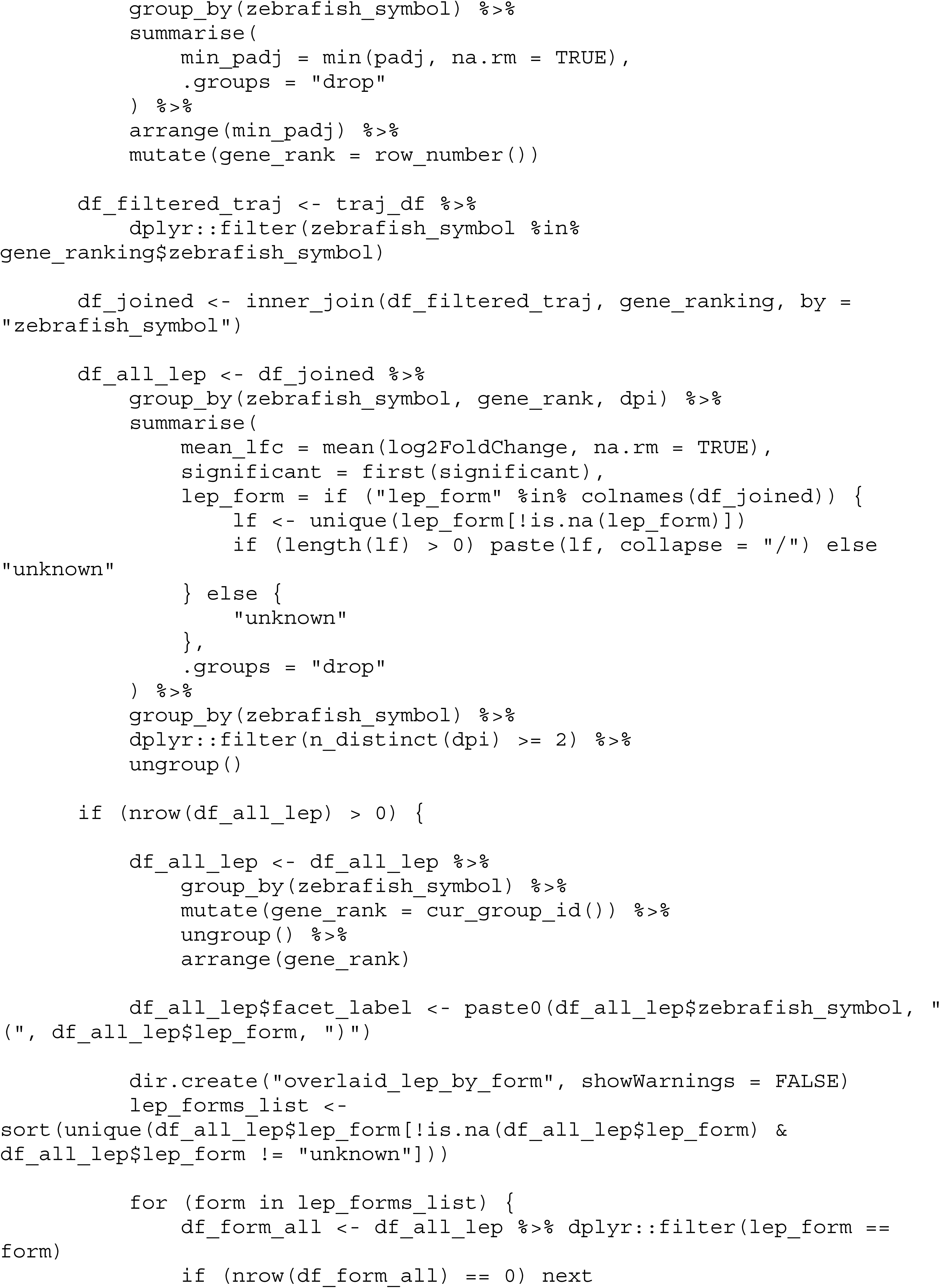

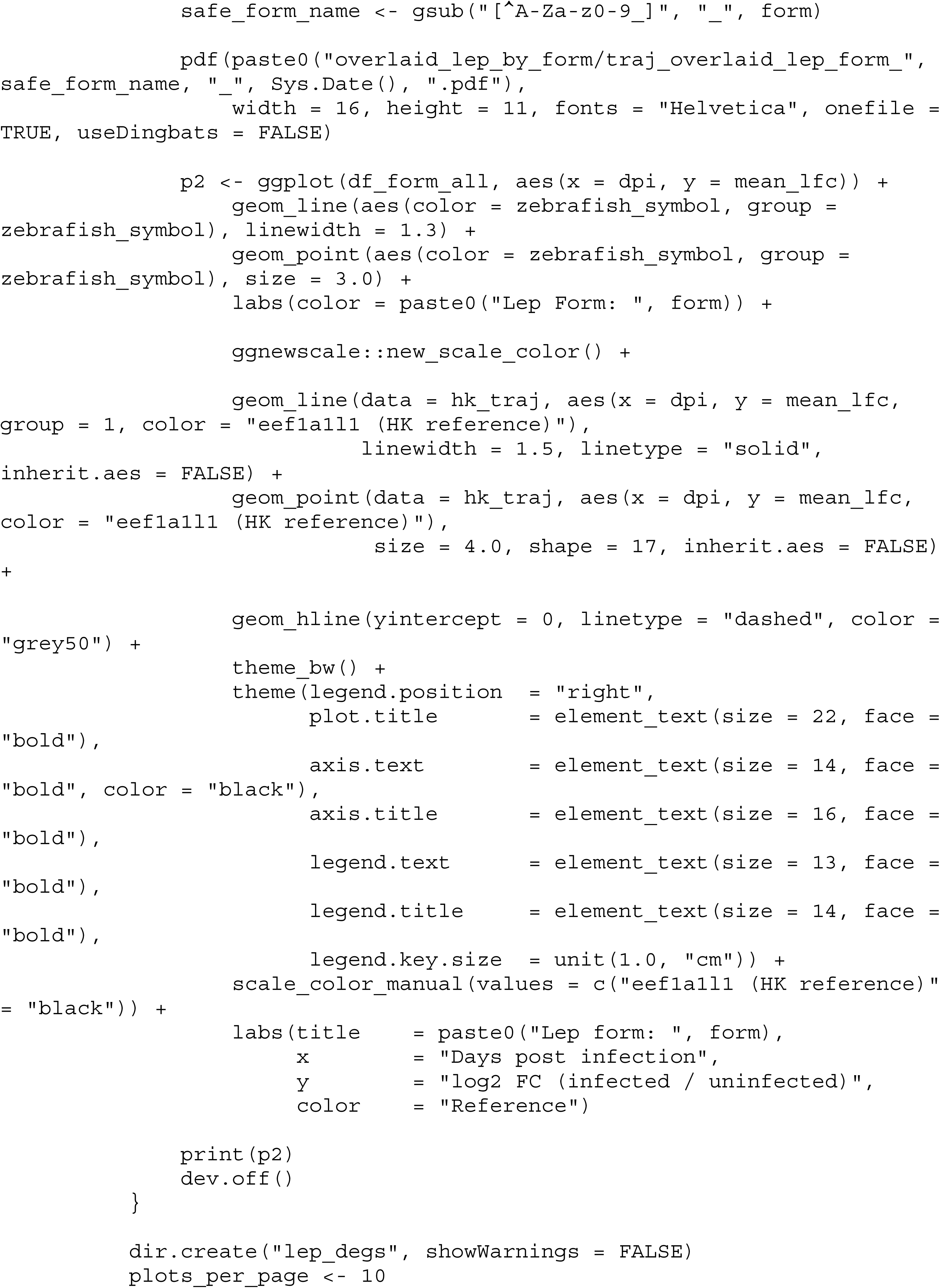

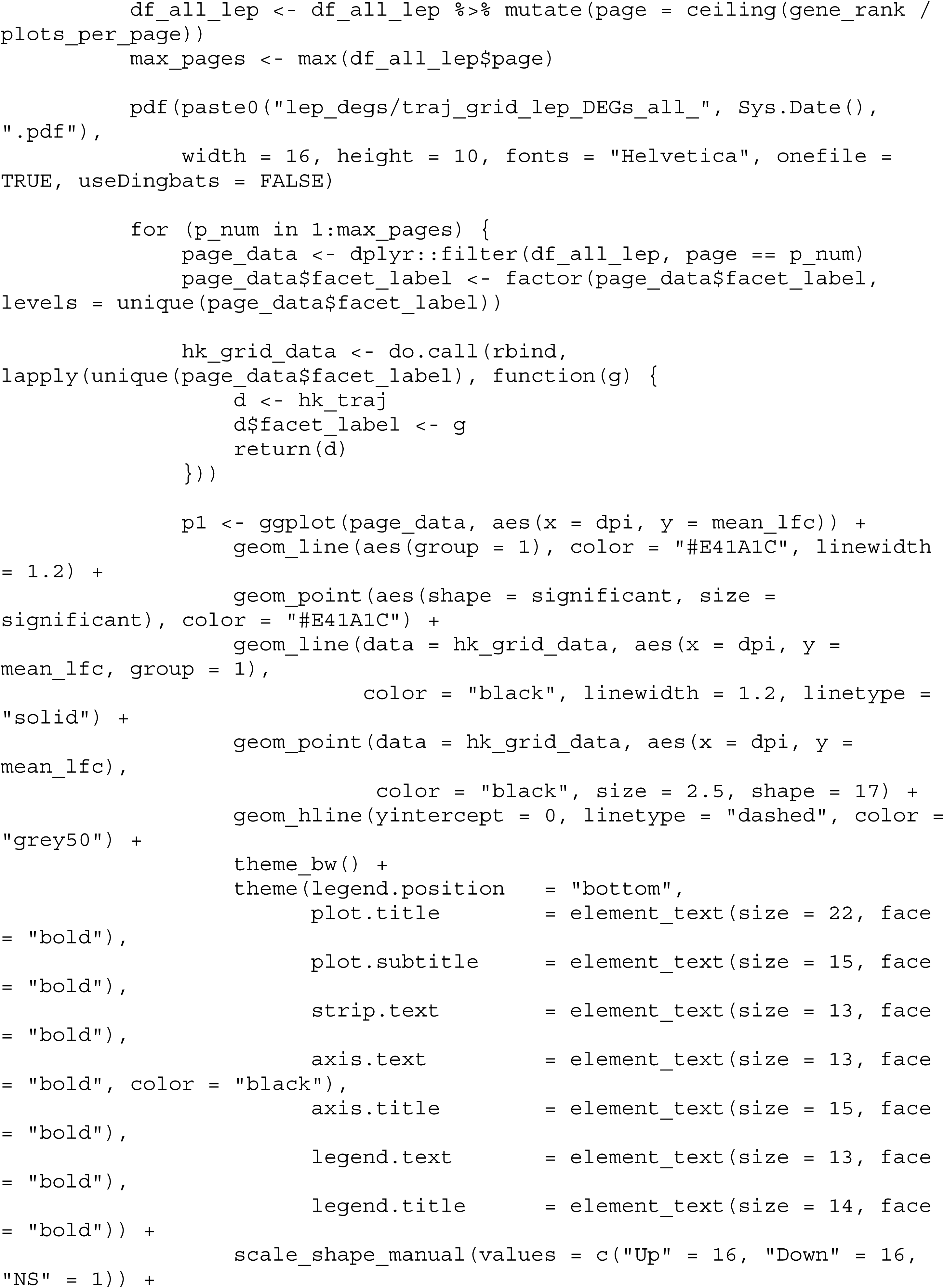

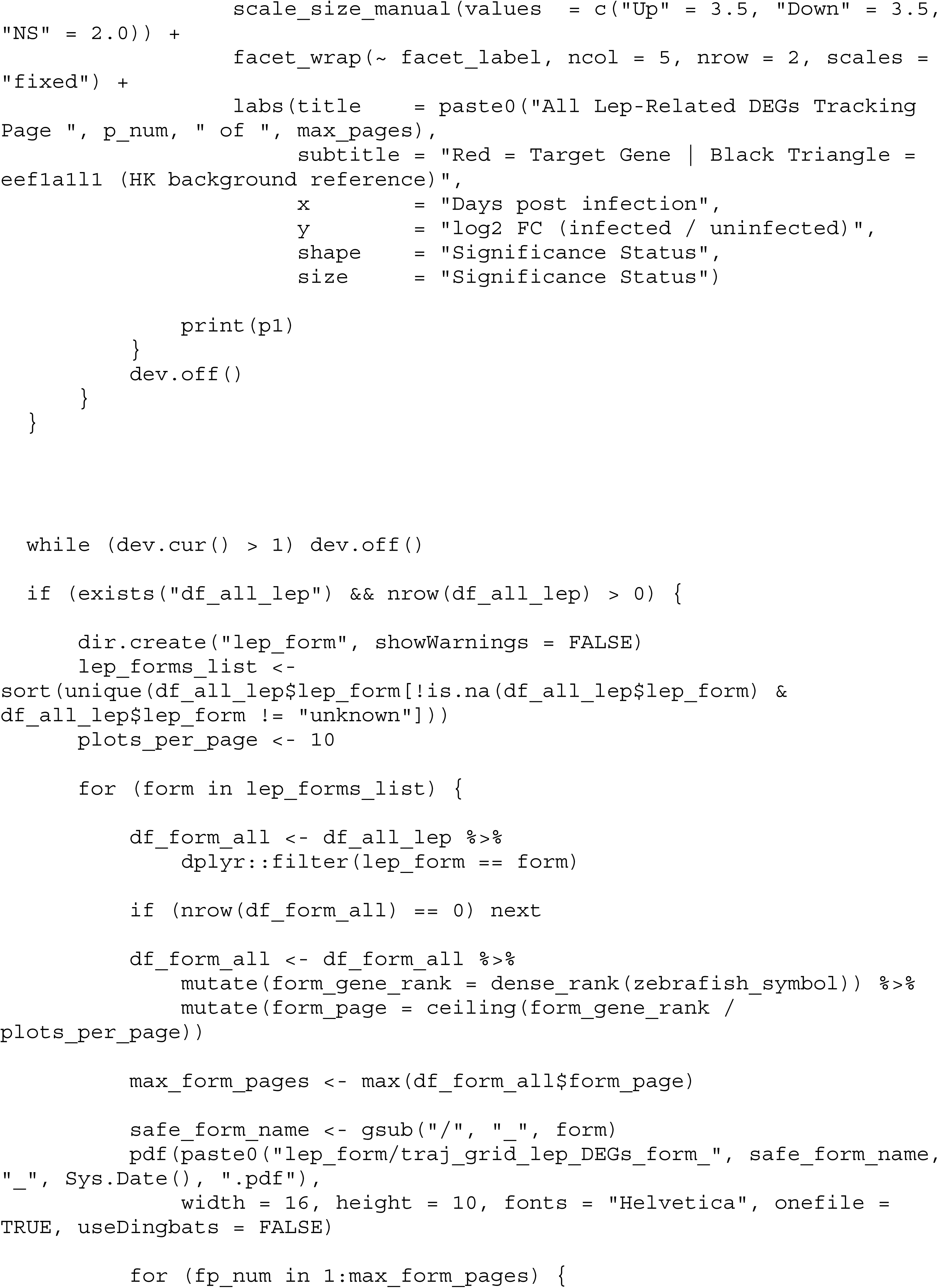

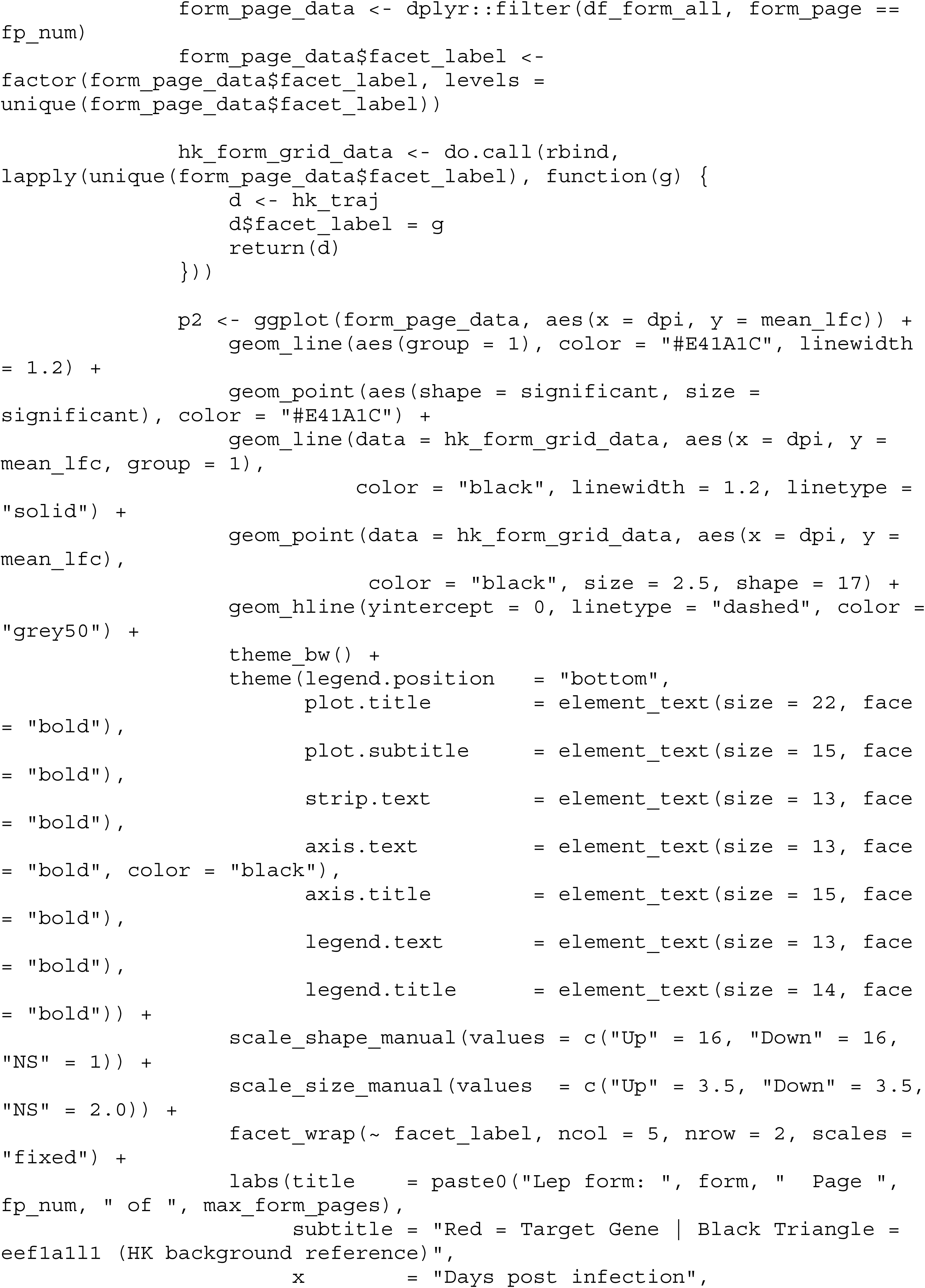

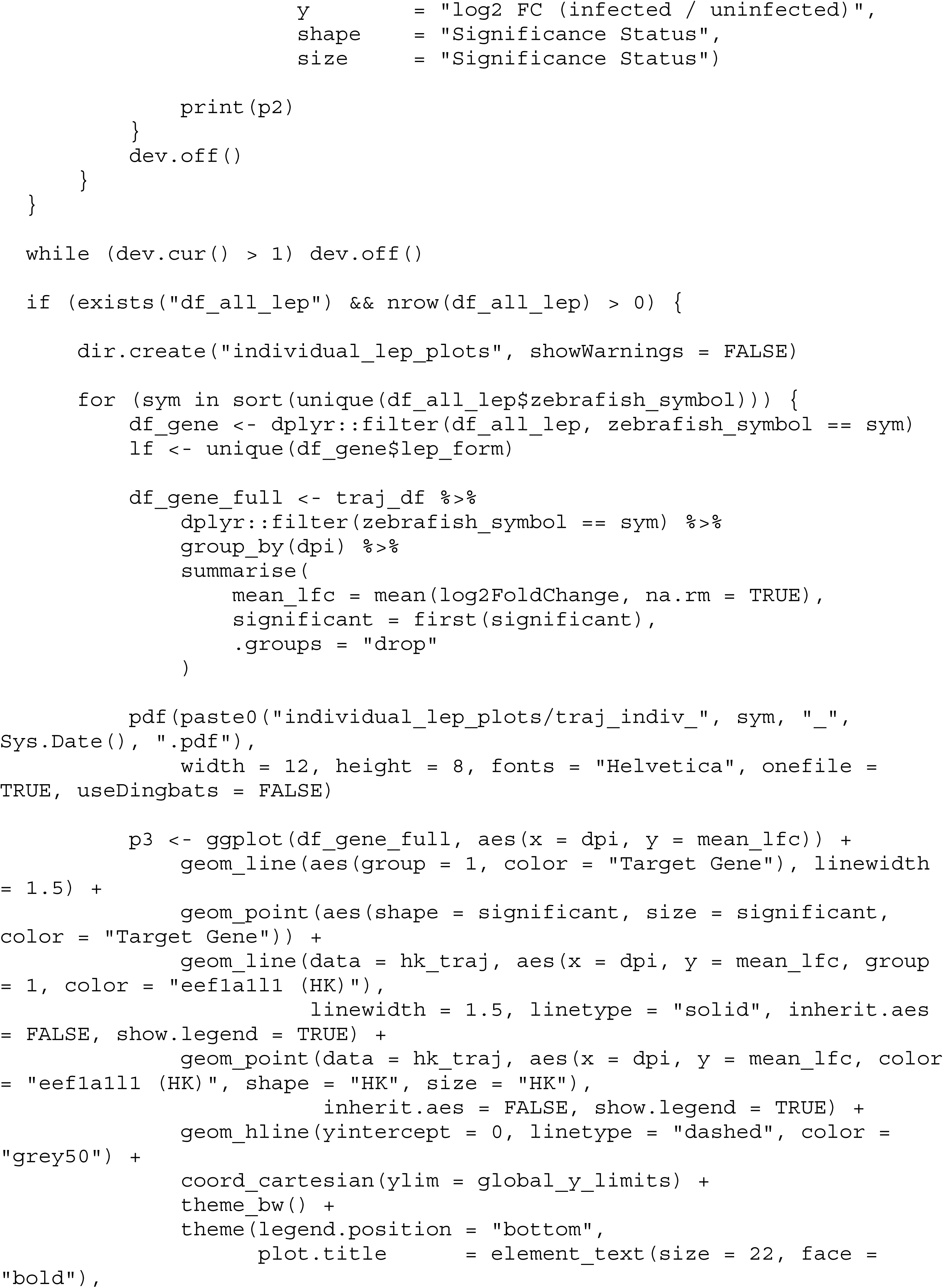

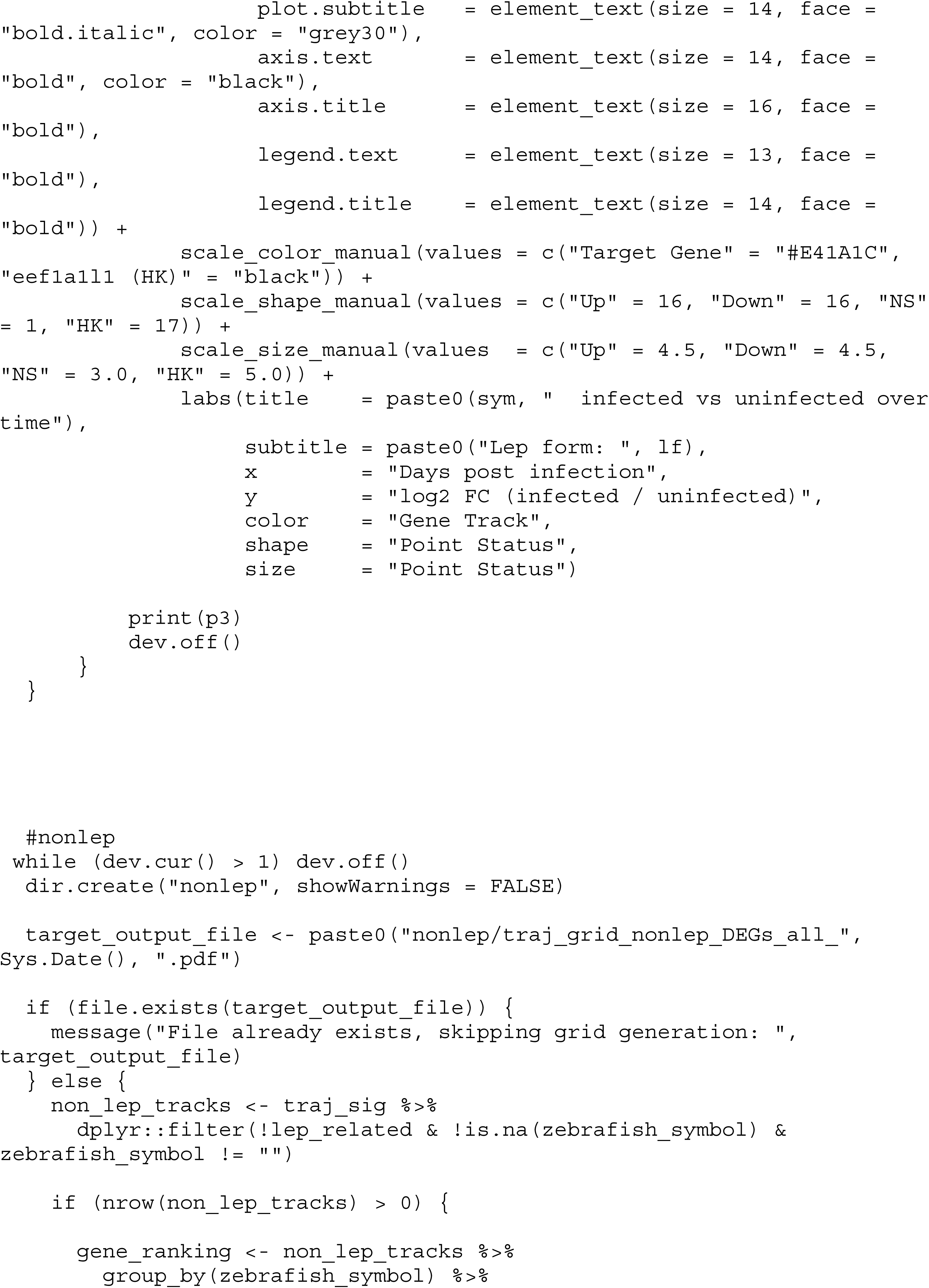

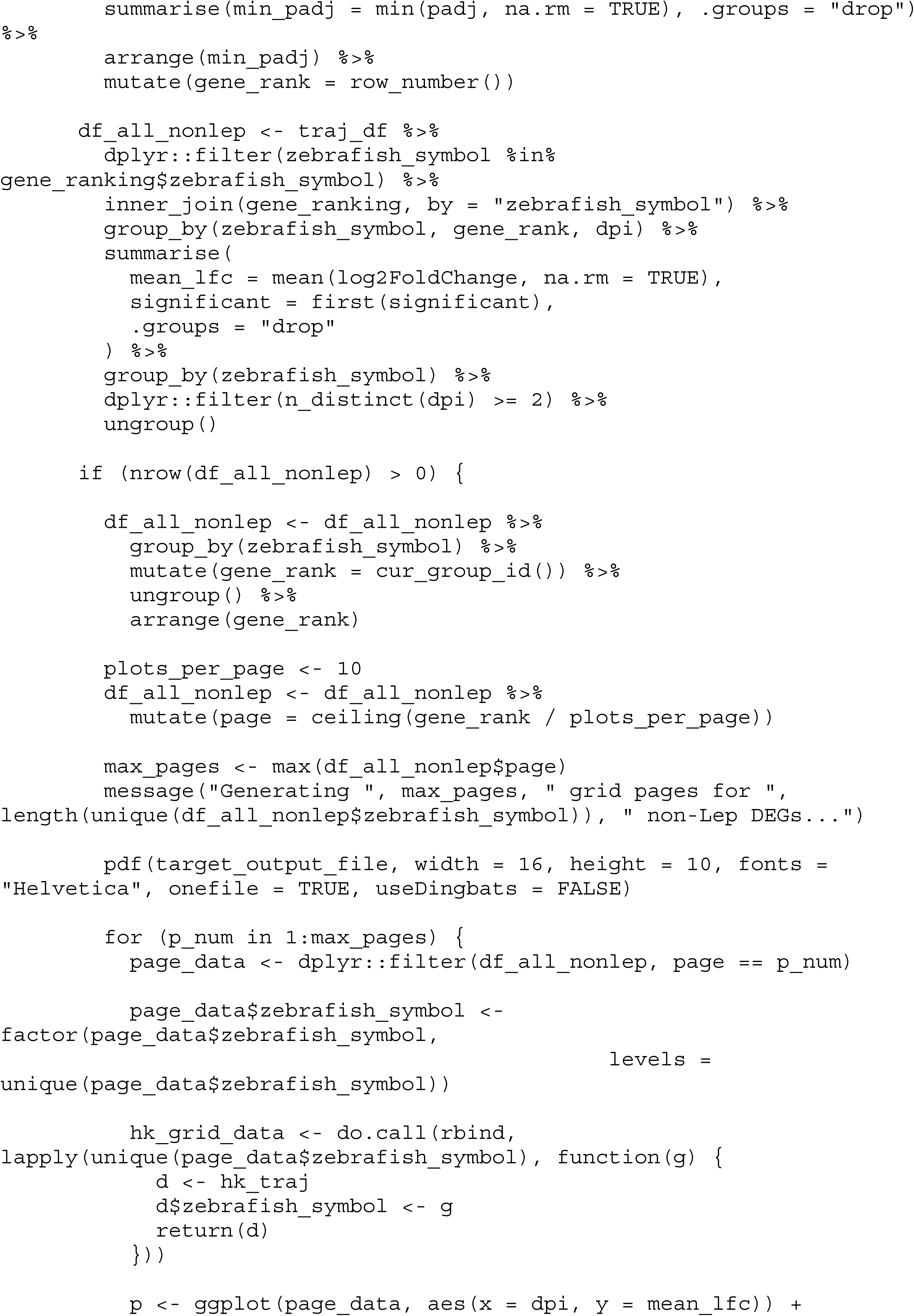

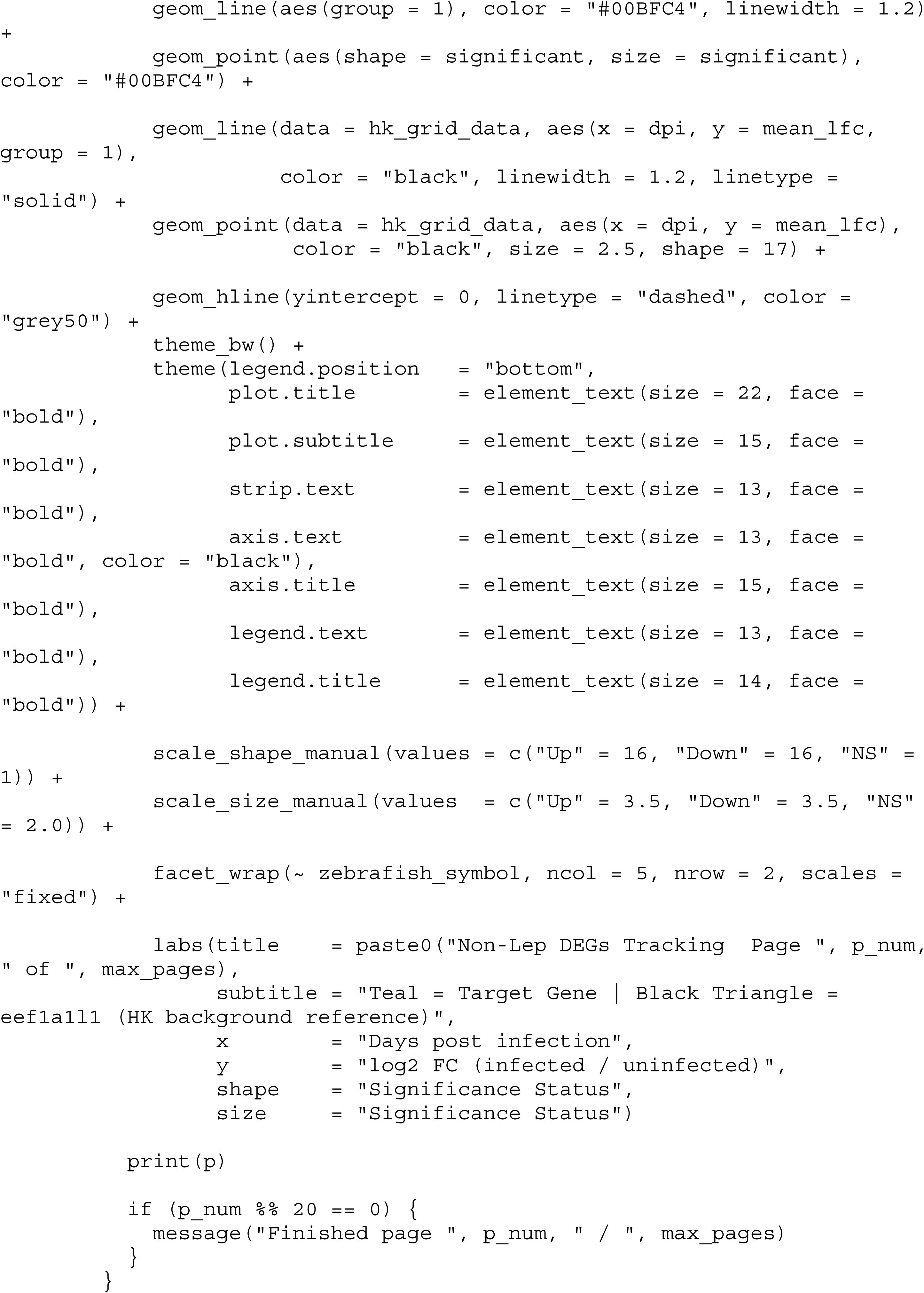

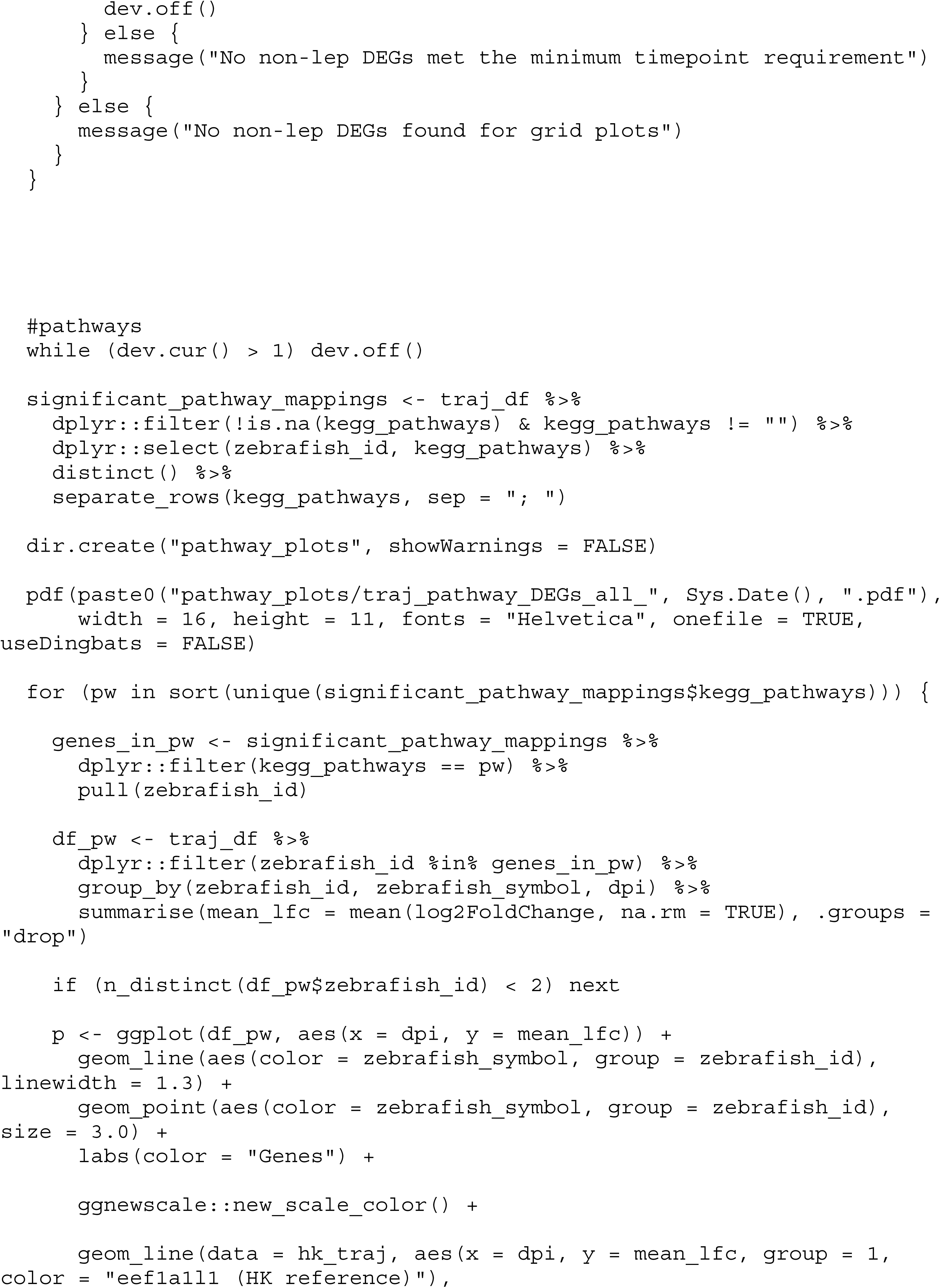

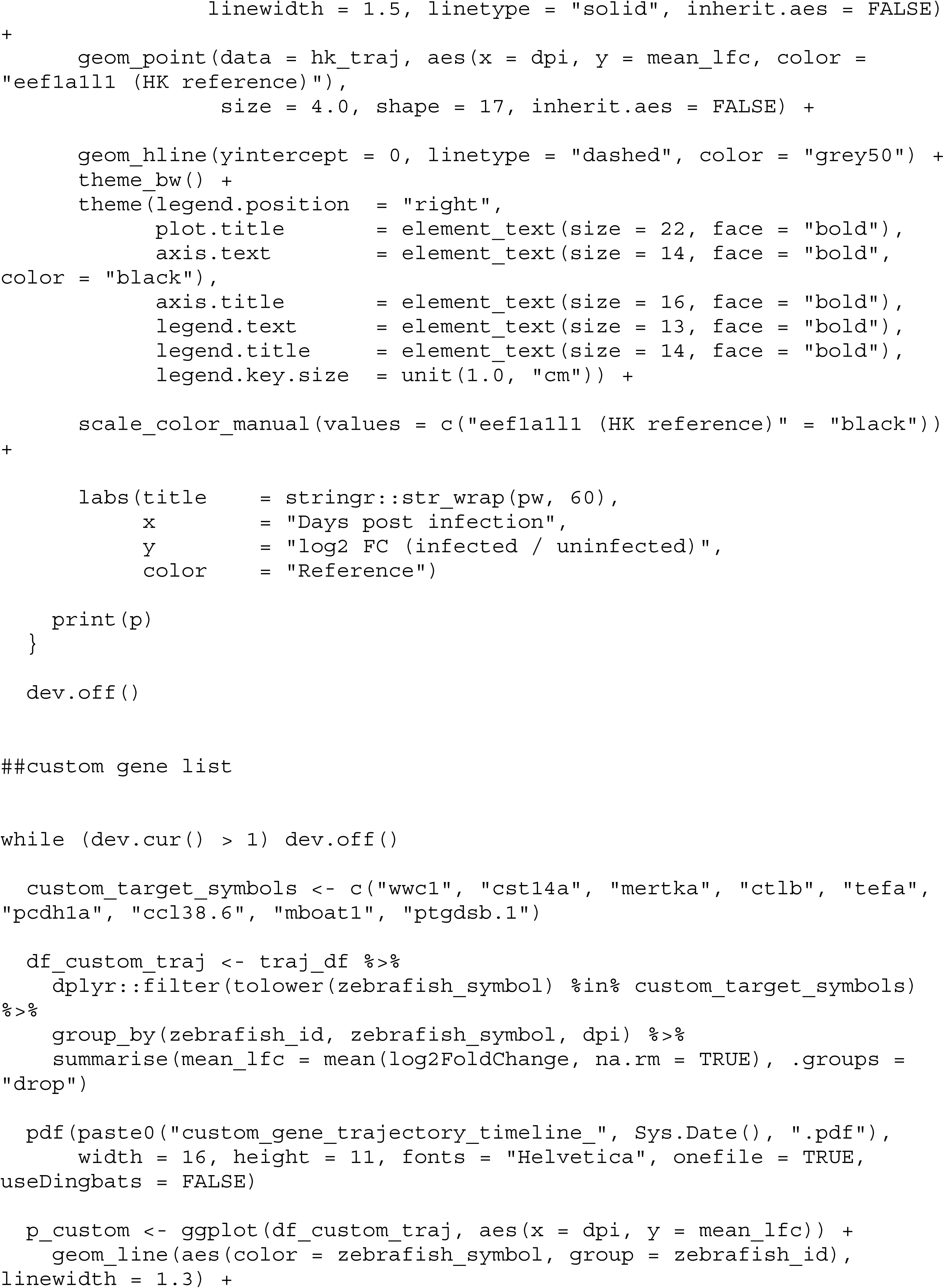

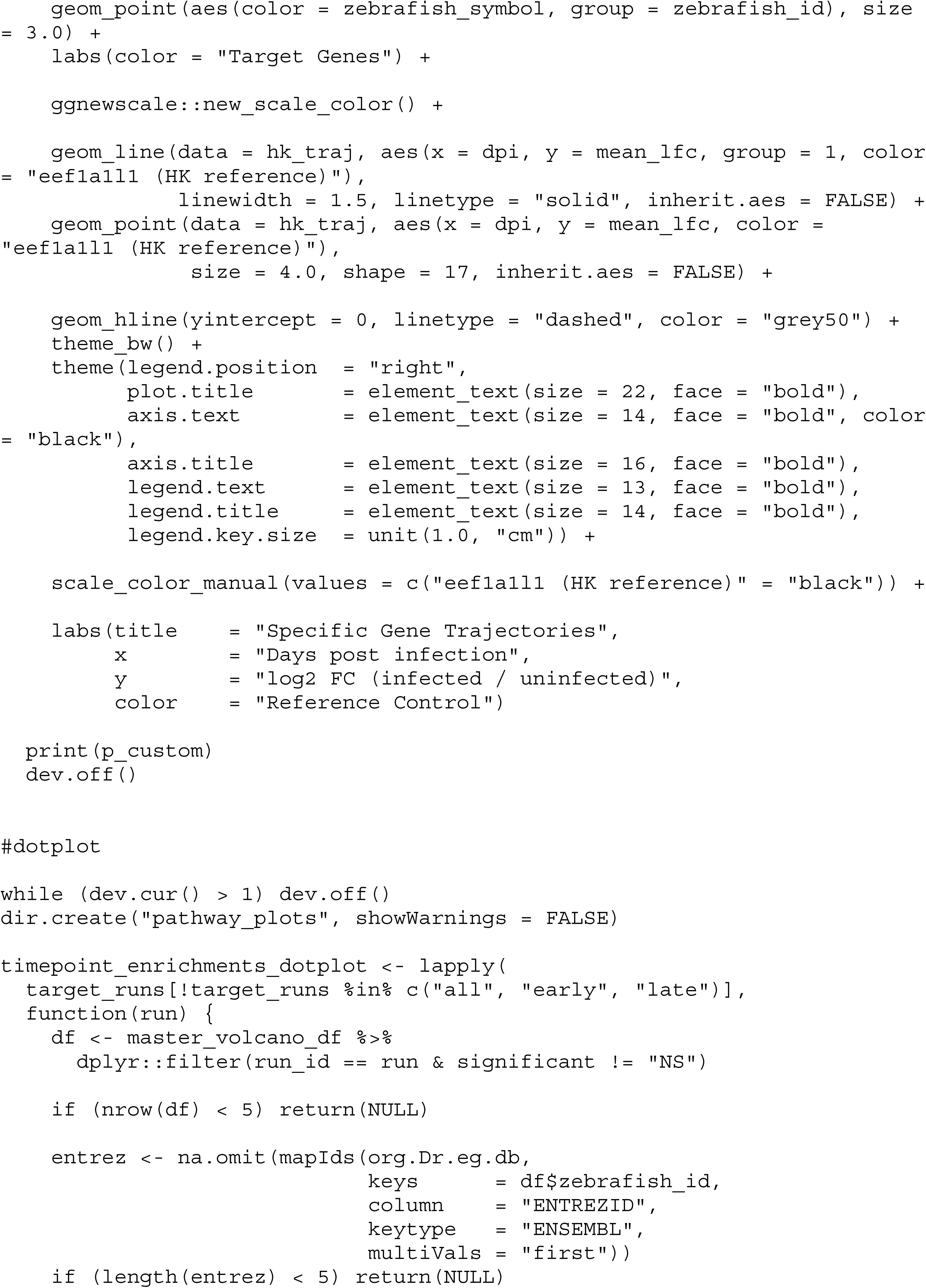

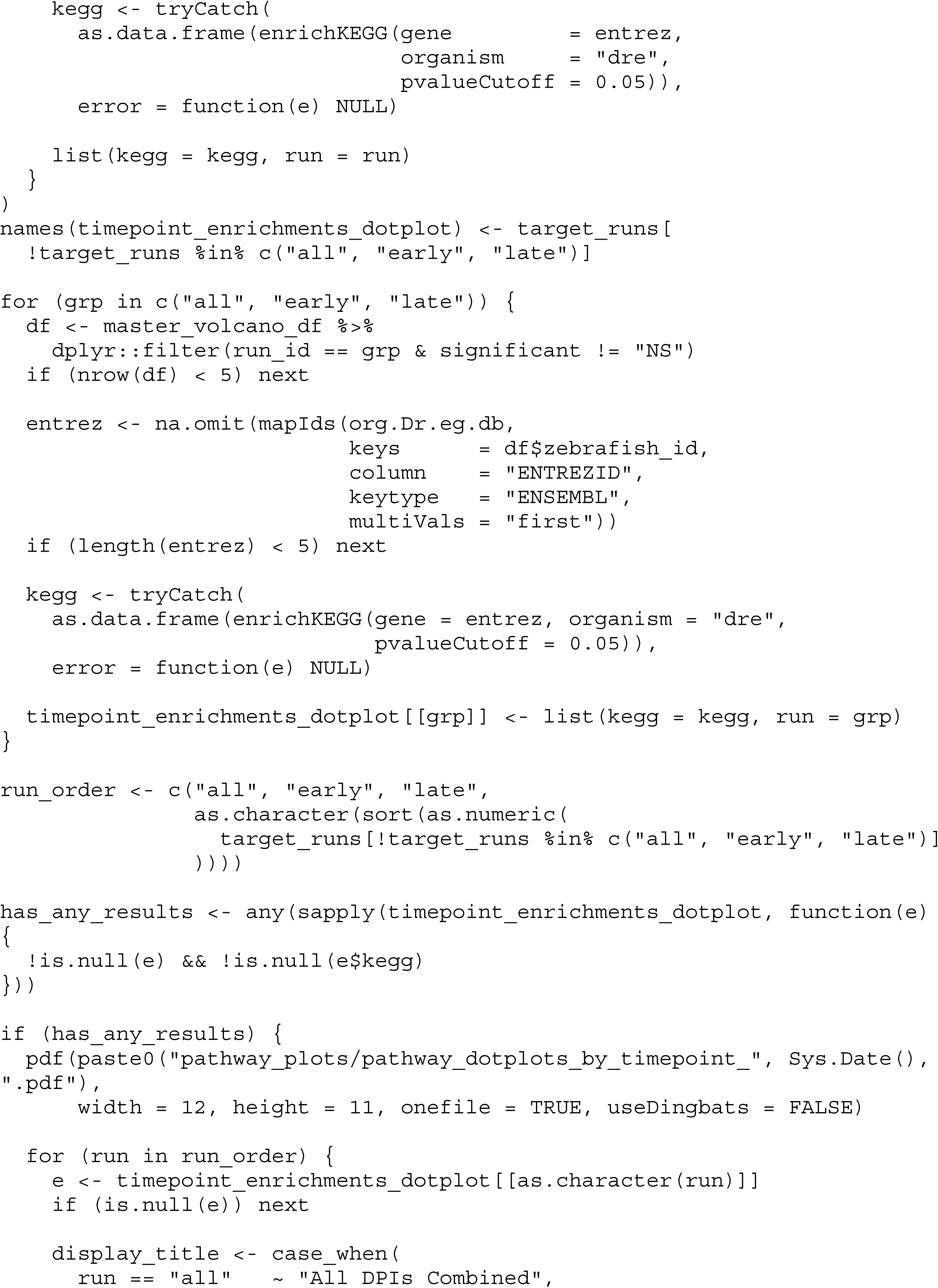

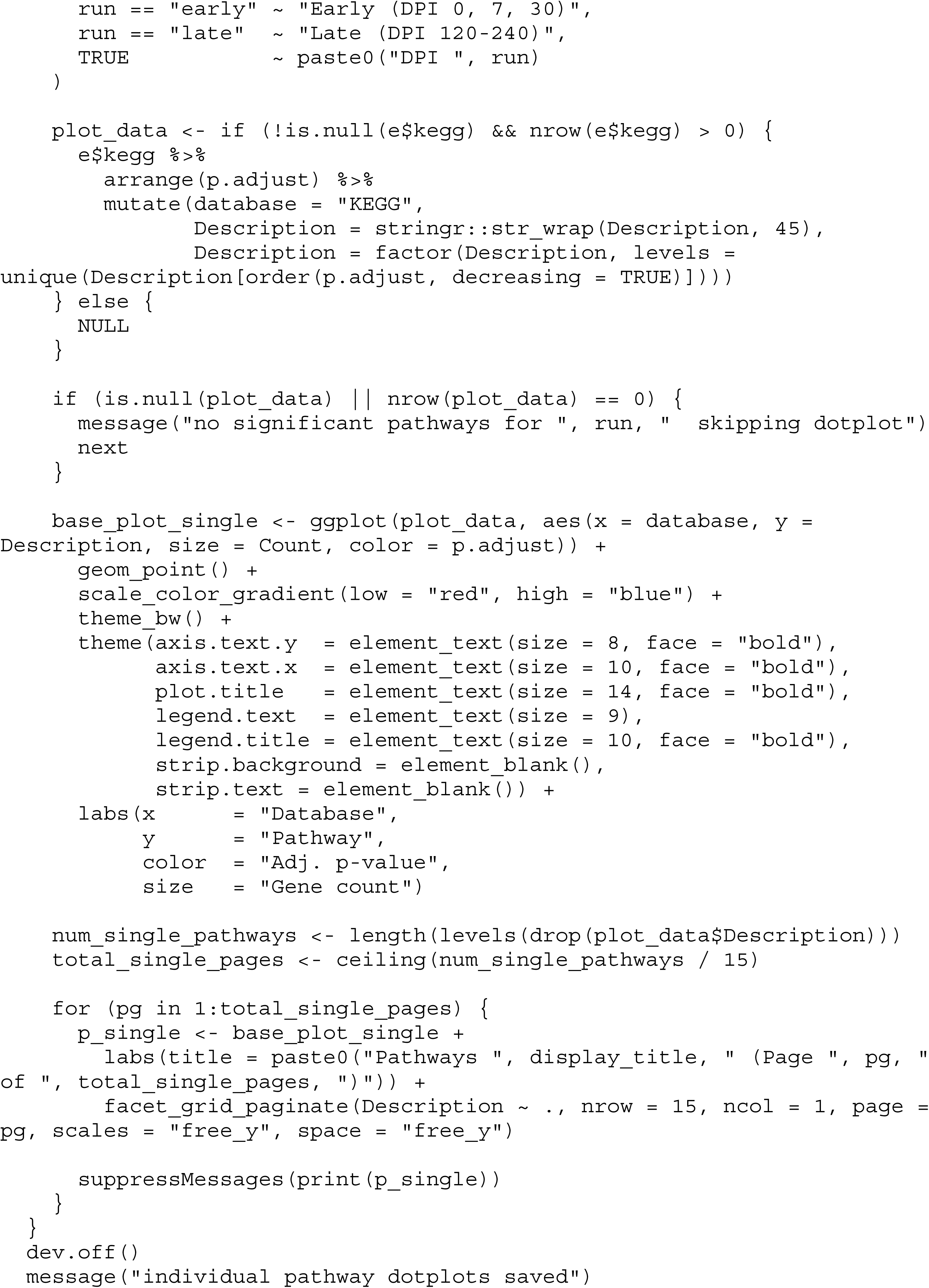

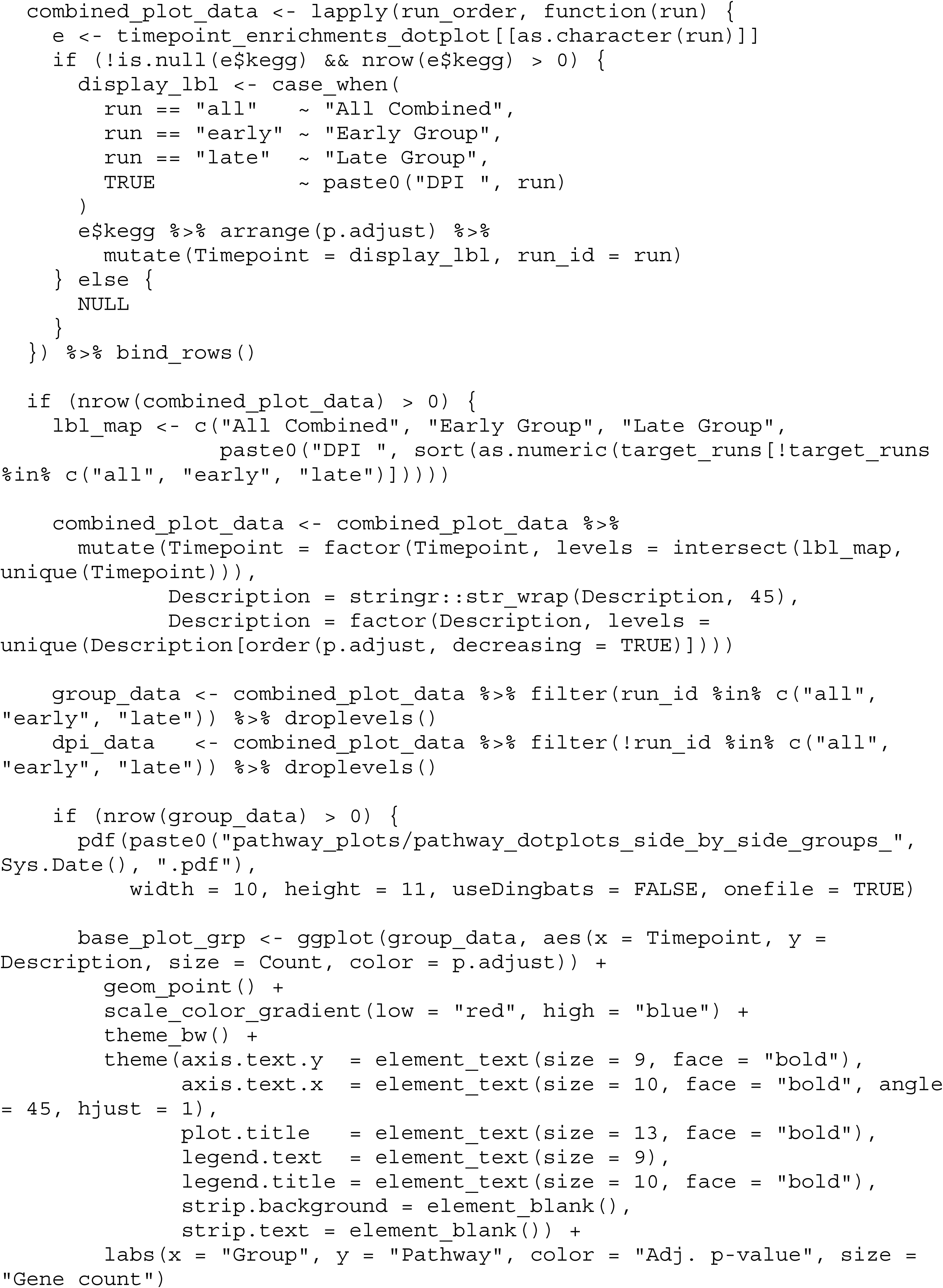

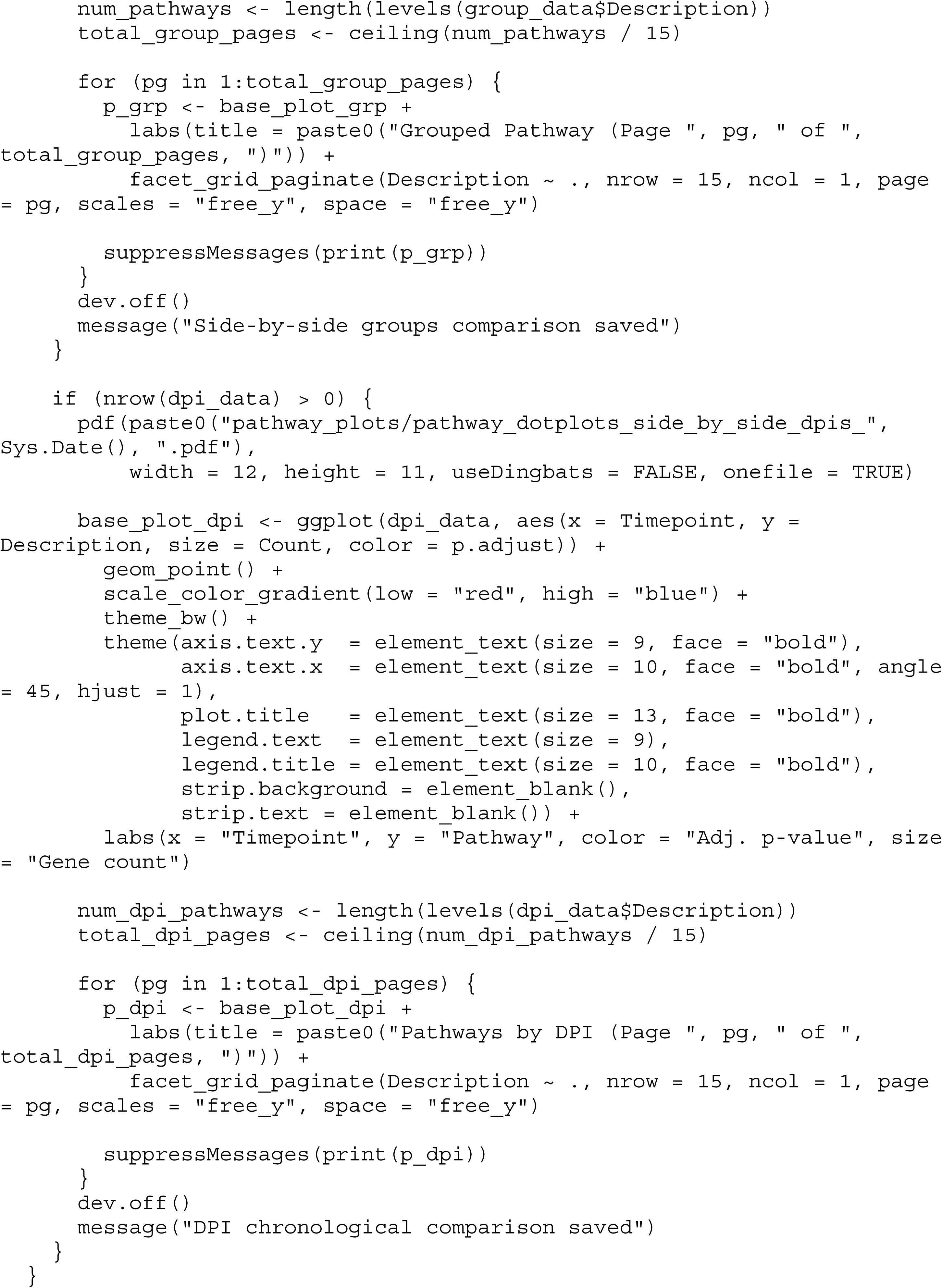

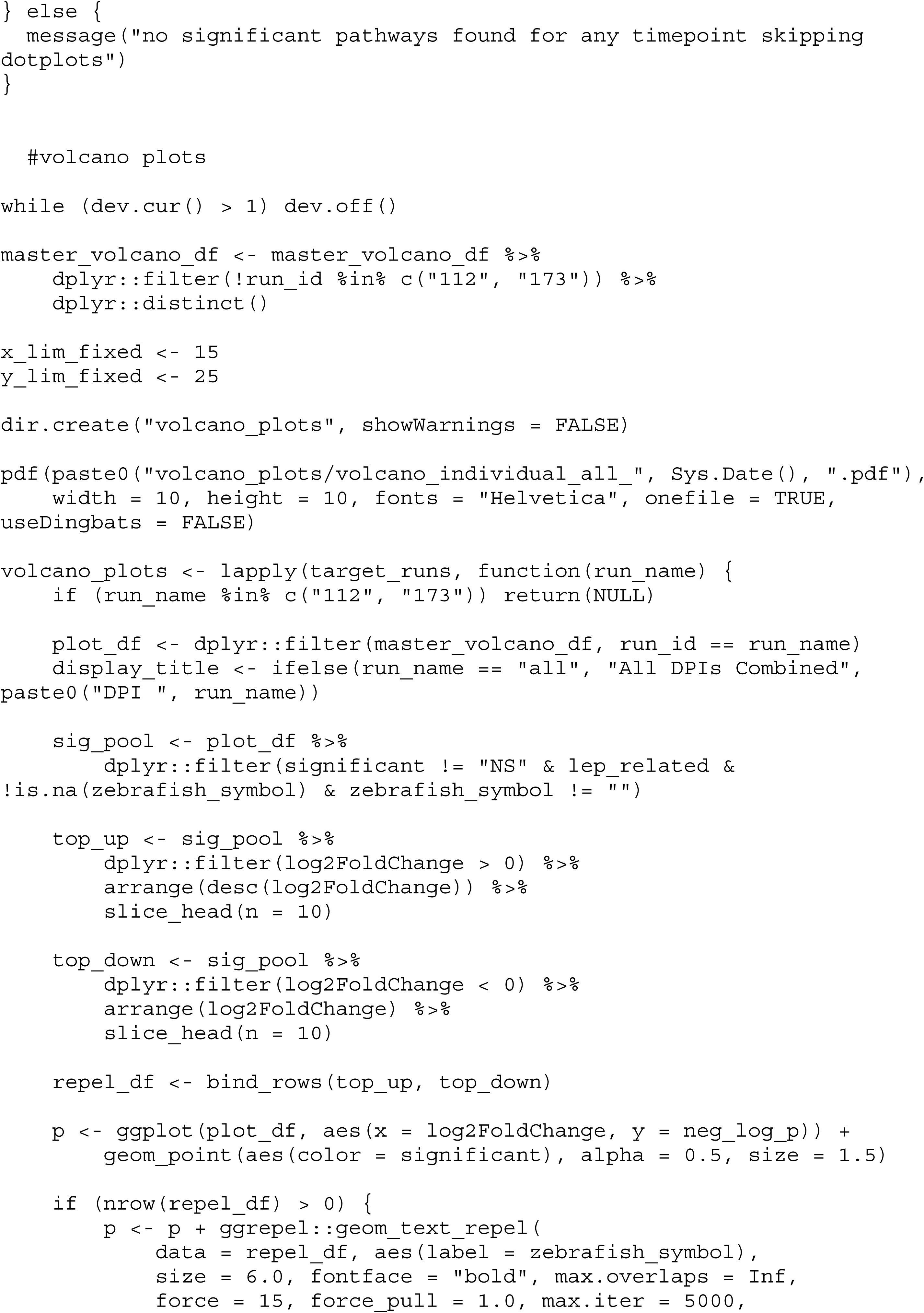

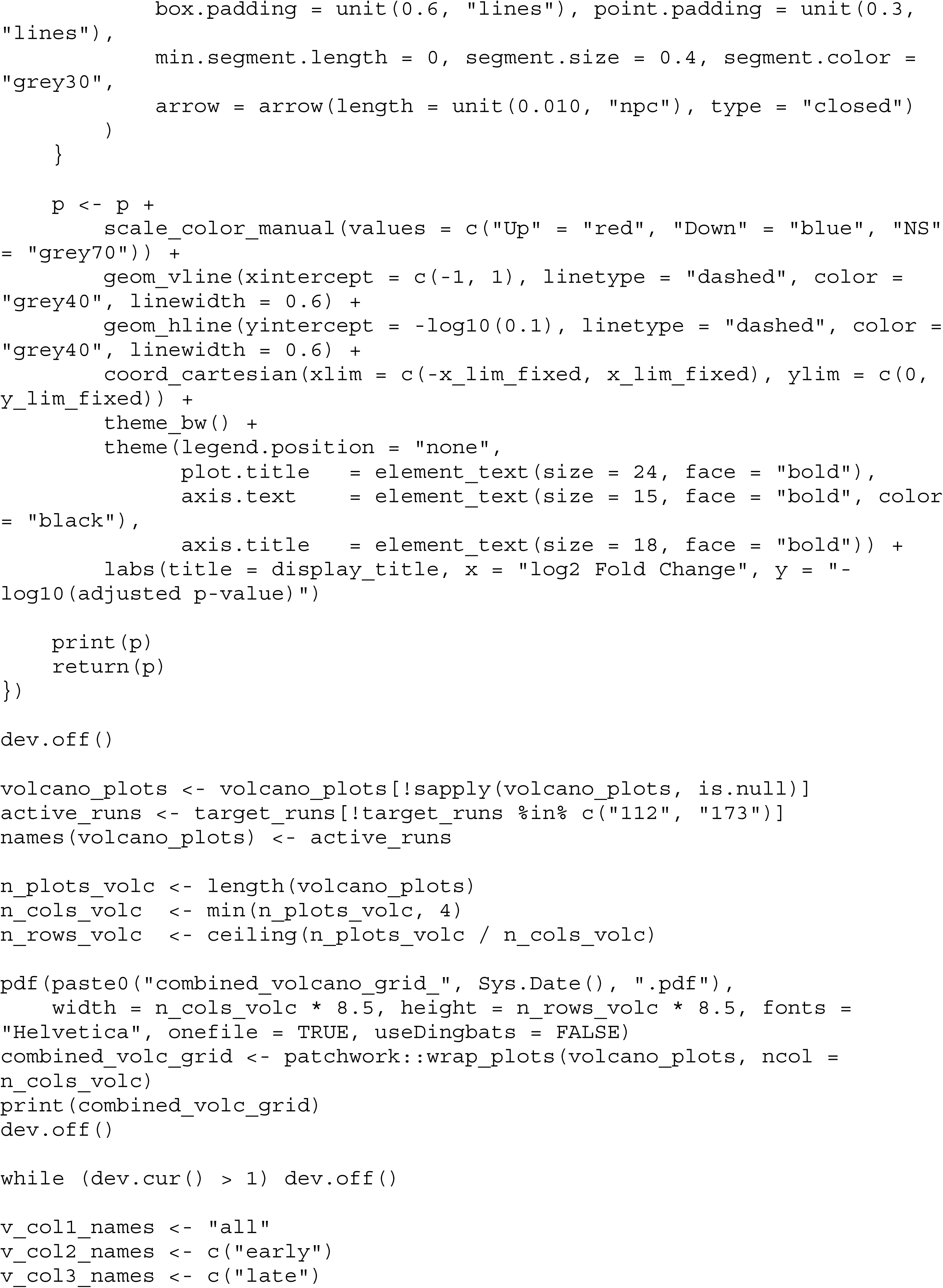

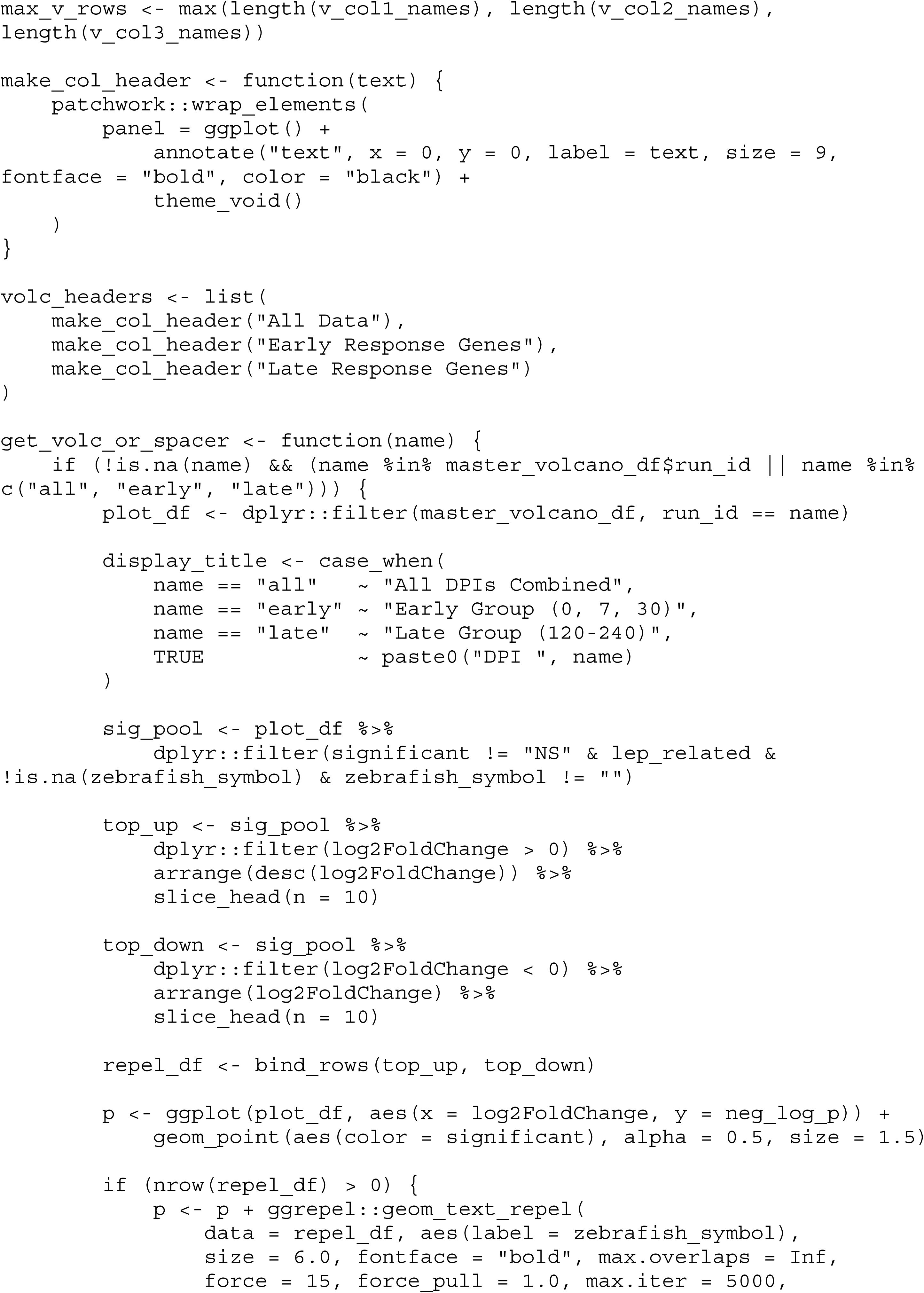

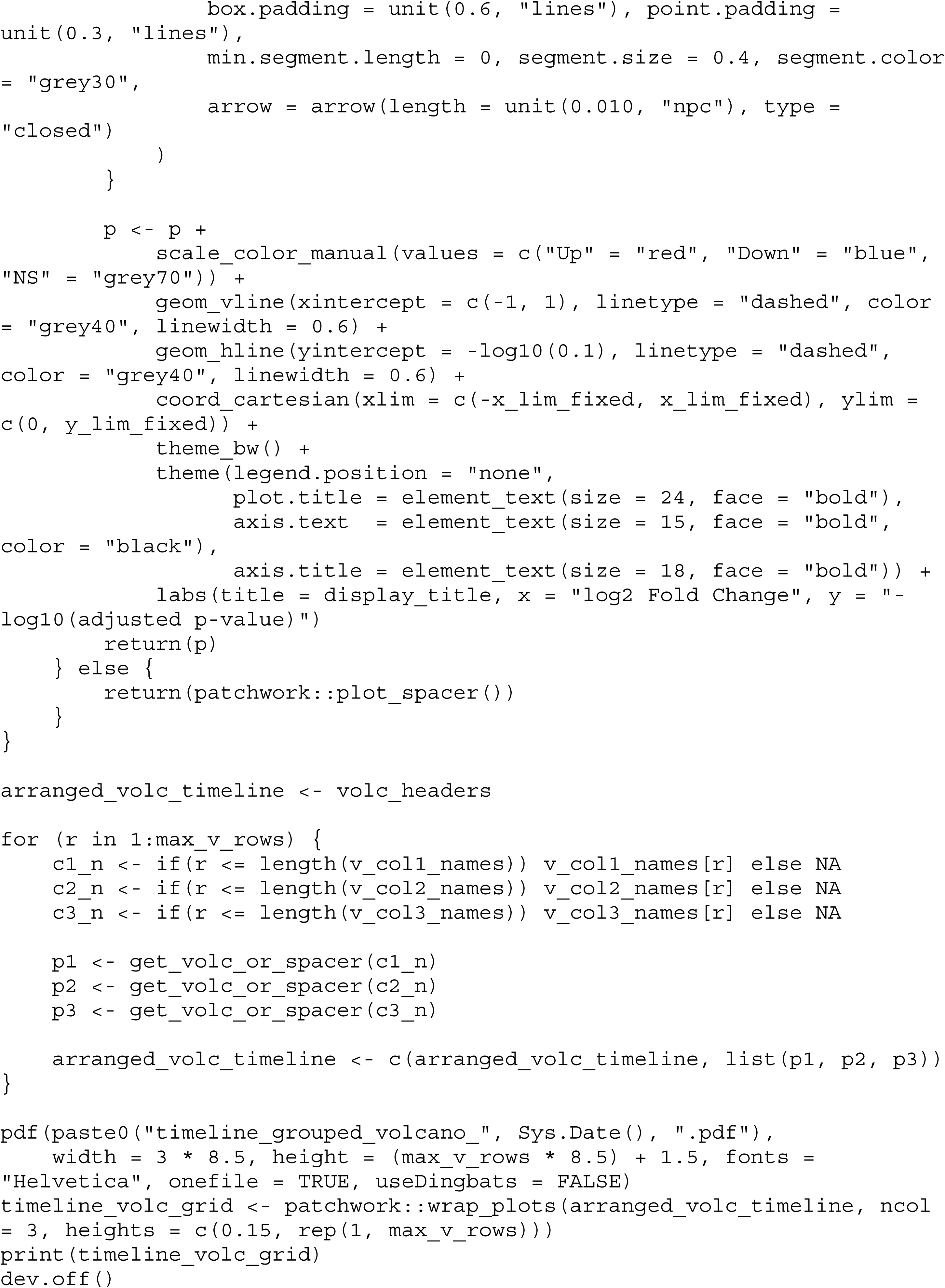

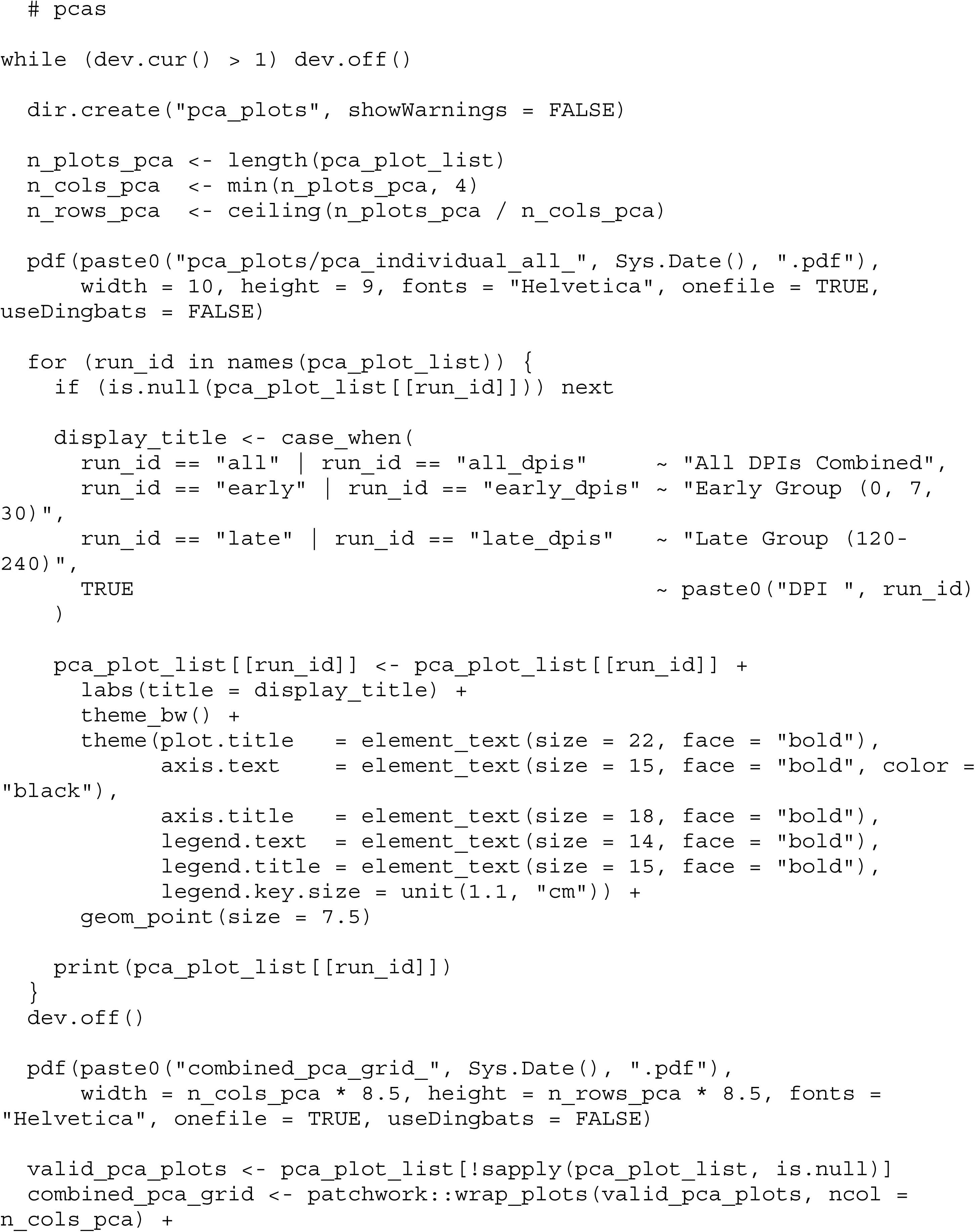

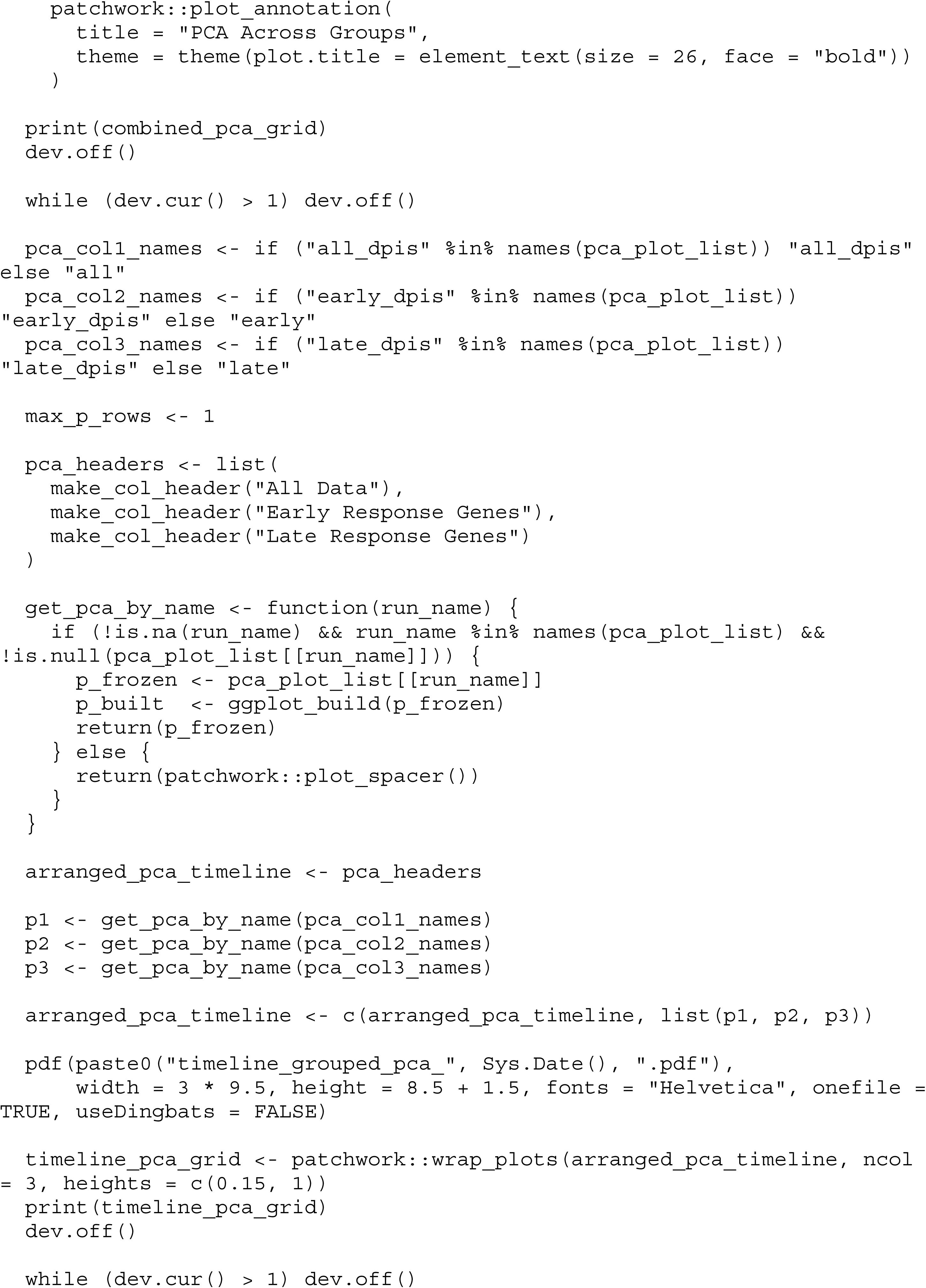

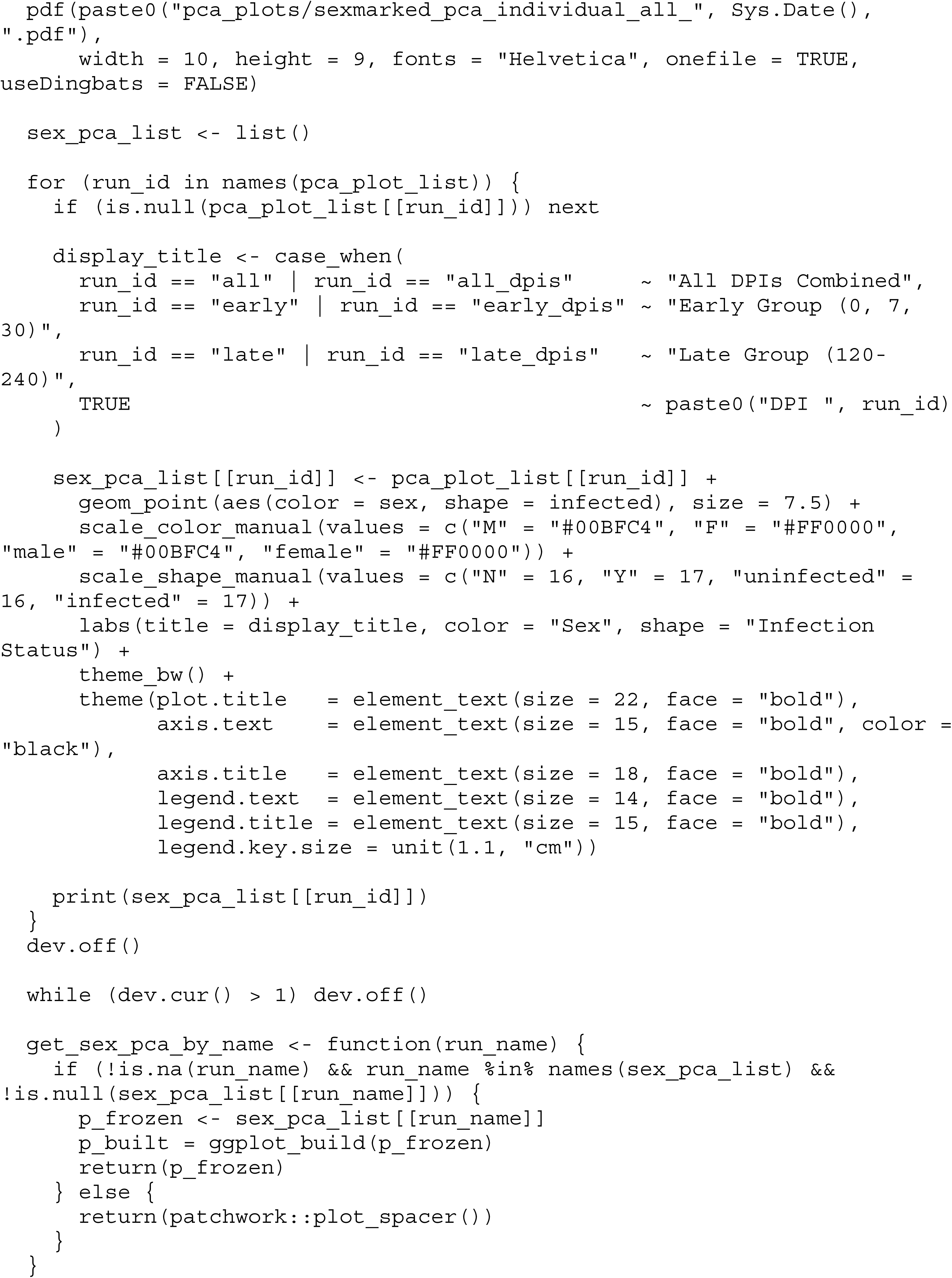

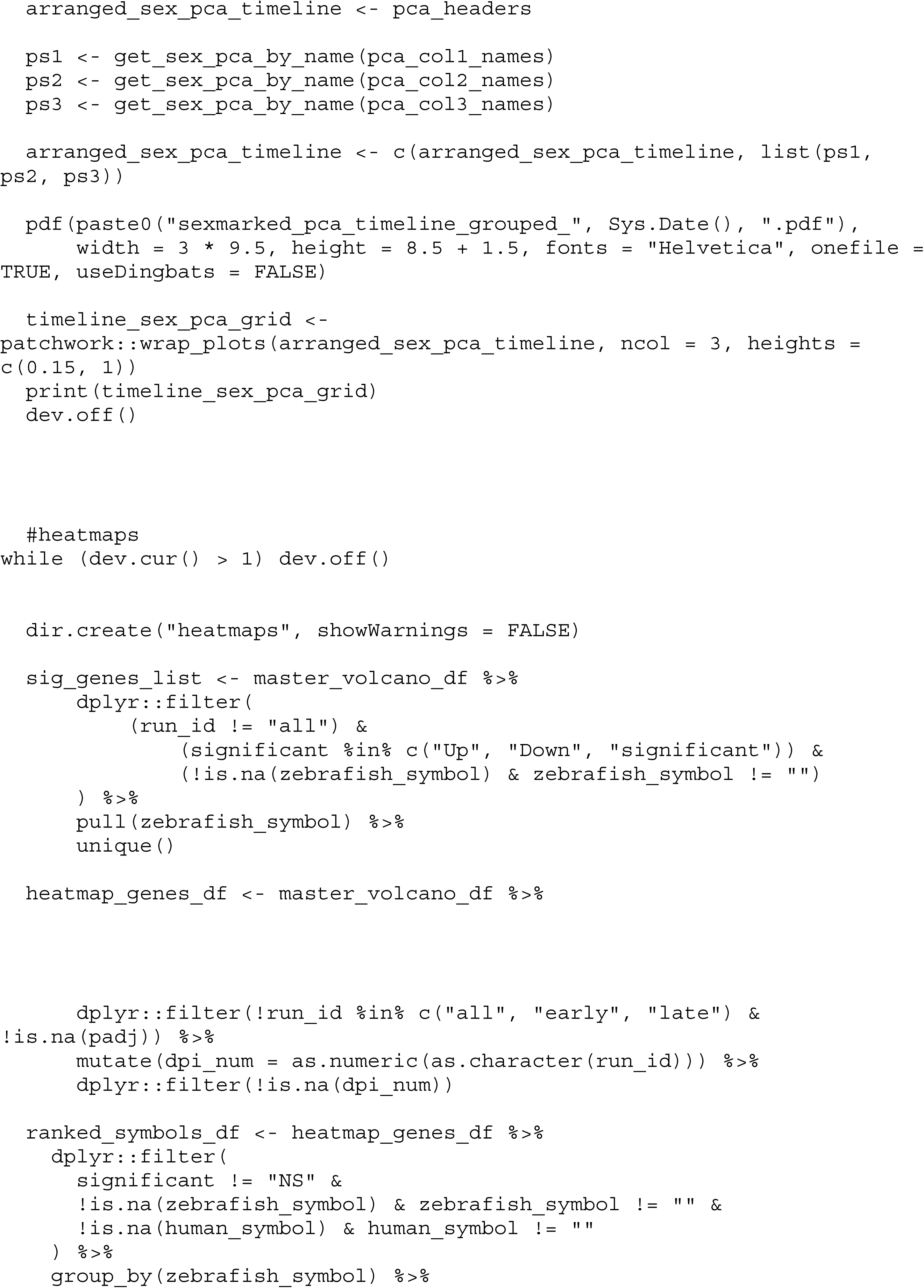

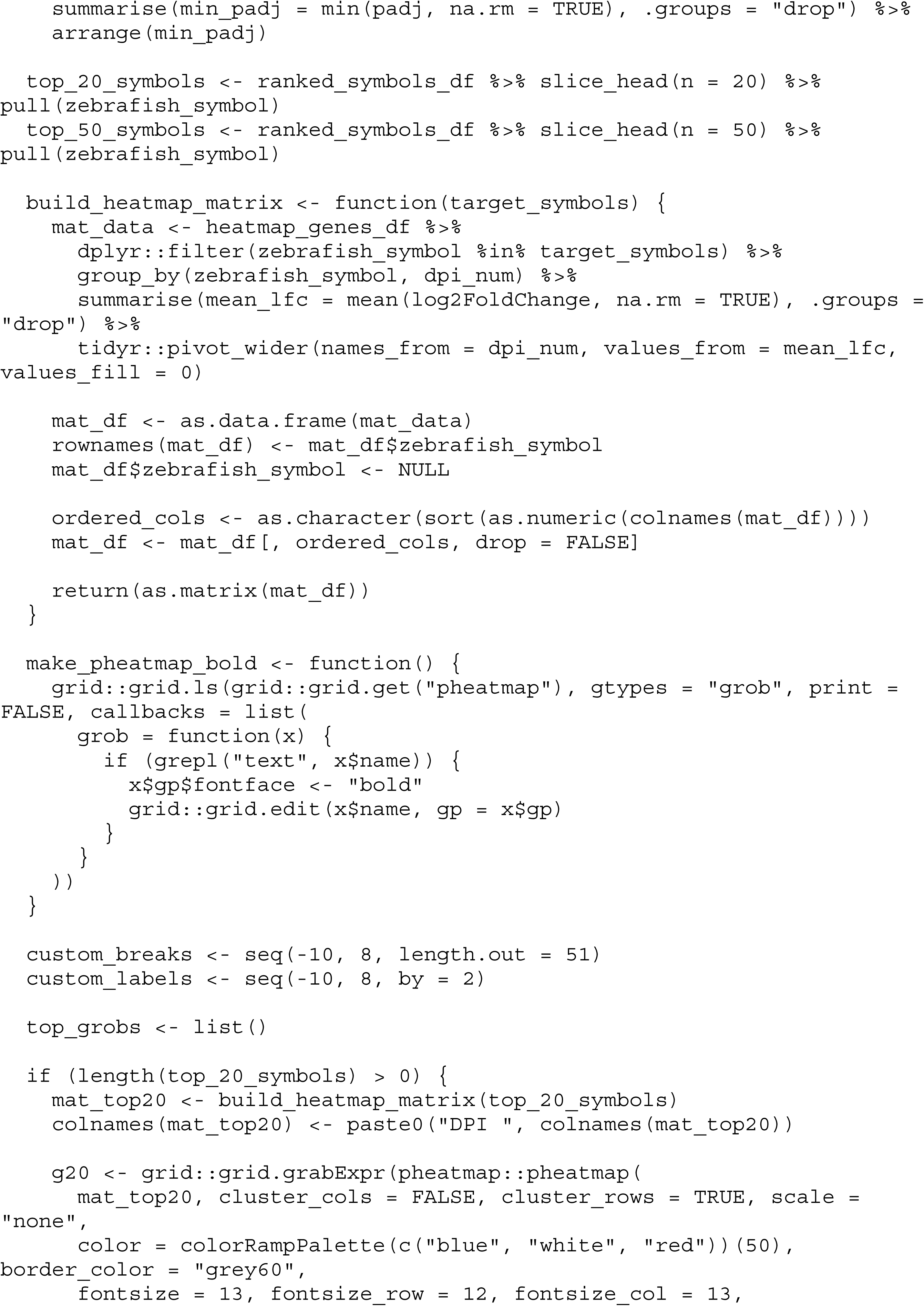

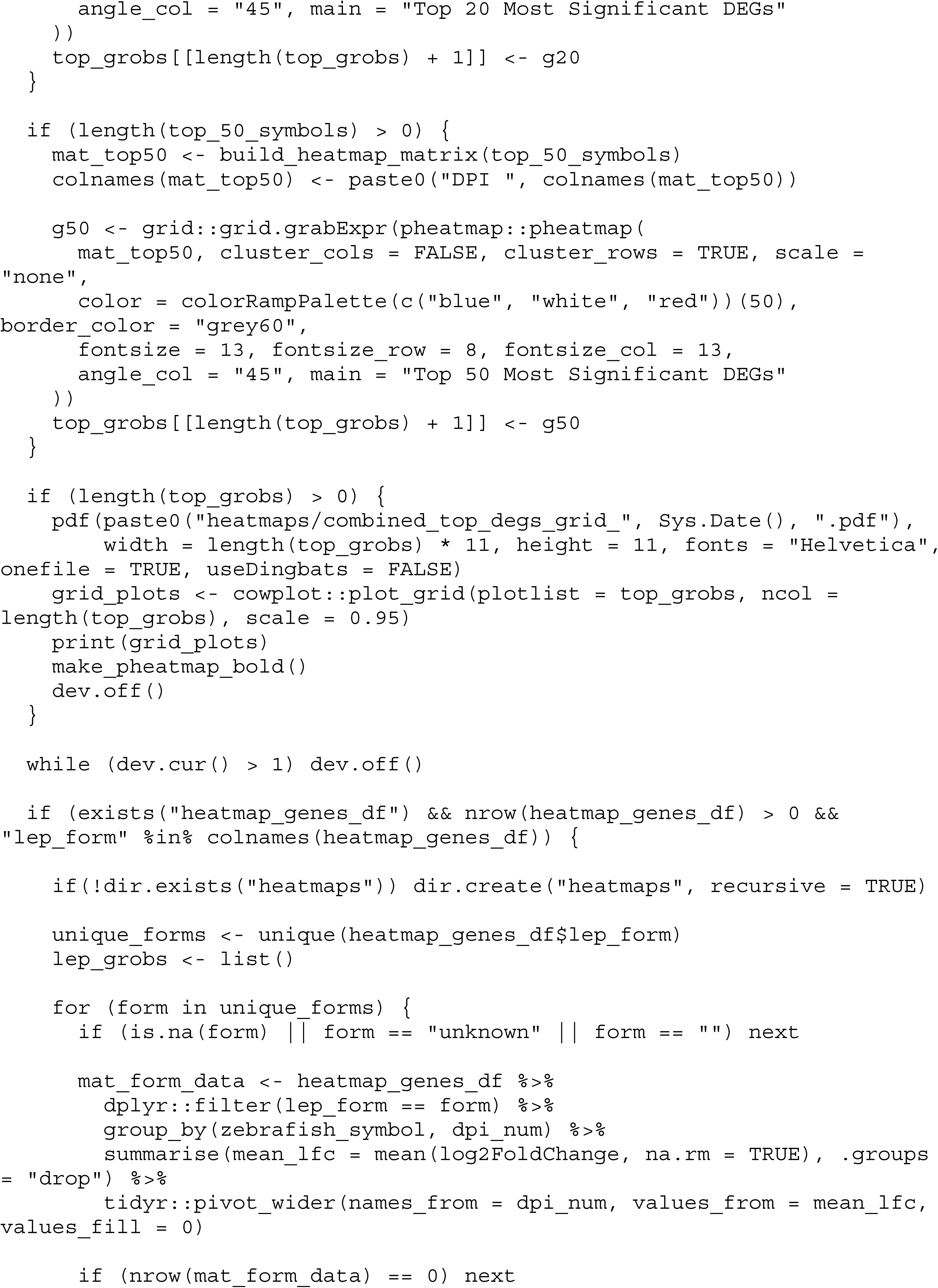

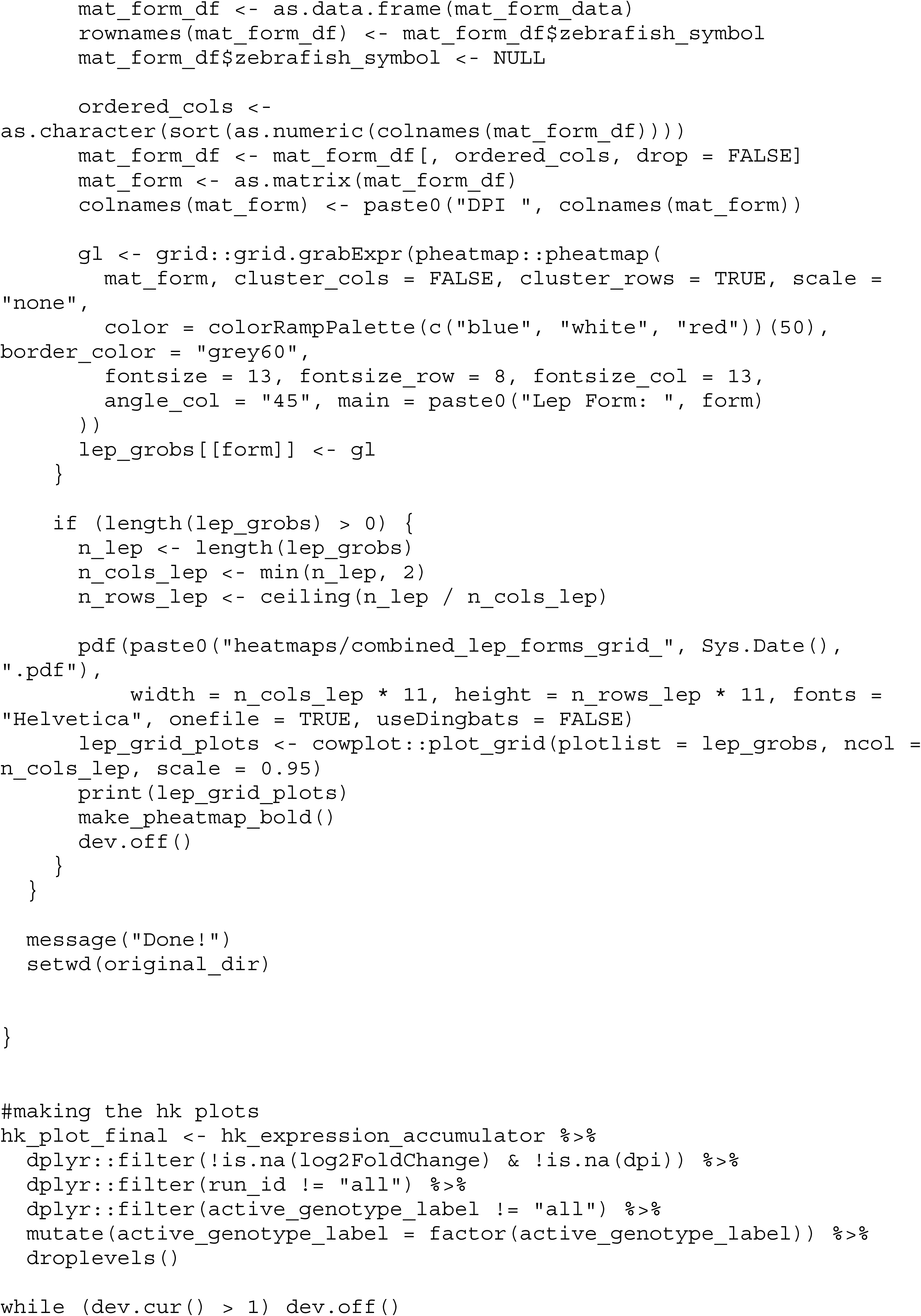

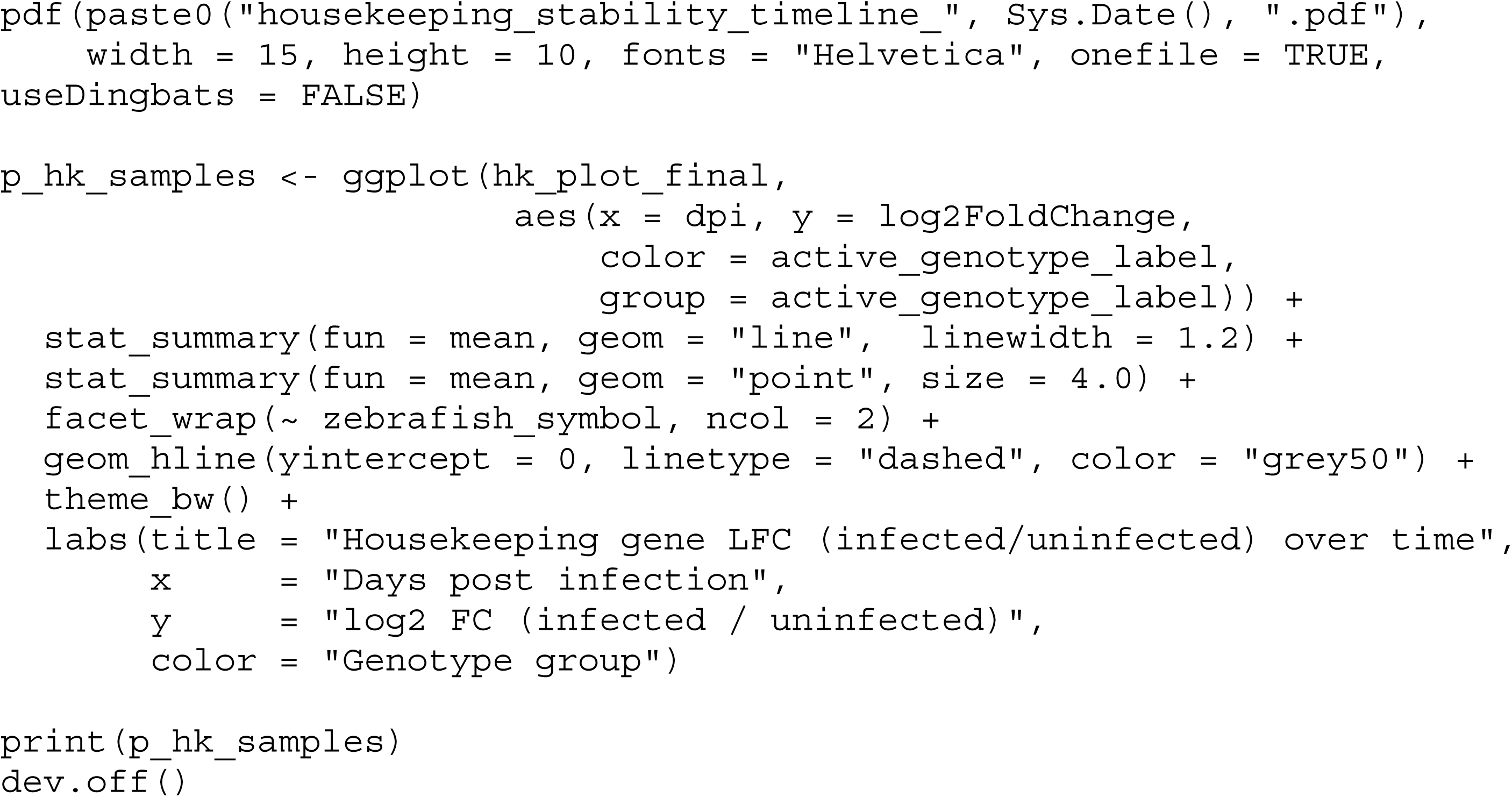

## Notes

### Competing Interest Statement

The authors have declared no competing interest.

### Funding

- Pew Charitable Trusts: 2021-A-17088
- NIH Common Fund: 1DP2NS127277
- Alfred P. Sloan Foundation: FG-2020-13447
- Searle Scholars Program
- CAPES-PrInt, Coordenação de Aperfeiçoamento de Pessoal de Nível Superior, Brazil

